# Massive programmed DNA elimination during embryogenesis in the trioecious nematode *Auanema rhodense*

**DOI:** 10.64898/2026.08.27.747507

**Authors:** Pablo Gonzalez de la Rosa, Liesl Grace Strand, Lewis Stevens, Manuela Kieninger, Joanna Collins, Margarethe Johansen, Tsz Wai (Bernice) Chan, Sally Adams, Anne Villeneuve, Andre Pires da Silva, Mark Blaxter

**Author notes:** **Corresponding author:** Mark Blaxter. **Name & Email** Pablo Gonzalez de la Rosa Liesl Grace Strand Lewis Stevens Manuela Keininger Joanna Collins Margarethe Johansen Anne Villeneuve Sally Adams Tsz Wai (Bernice) Chan Andre Pires da Silva Mark Blaxter.

## Abstract

In animals, the germline is usually set aside early in development and its genome is protected to ensure faithful transmission of genetic information to future generations. Genetic alterations in somatic cells are not inherited by progeny, and the somatic genome does not have to be protected from change in the same way. In some species, programmed DNA elimination (PDE) results in the directed loss of genetic material from somatic cells. Here, we report extreme PDE in *Auanema rhodense*, a free-living nematode with a remarkable, trioecious life history and unusual patterns of sex chromosome inheritance. We find that nearly two thirds of the *A. rhodense* germline genome is eliminated from somatic cells, with DNA lost from chromosome ends as well as within chromosomes, resulting in fragmentation of the seven germline chromosomes into fourteen somatic chromosomes. Most eliminated DNA comprises multi-megabase tandem repeat blocks. Eliminated DNA on the X chromosome harbours germline-restricted repeats that are distinct from those on the autosomes. The eliminated DNA includes many highly-repeated non-coding RNA loci but few protein-coding genes. Elimination sites are strongly associated with a palindromic sequence motif that likely directs DNA breakage and new telomere repeat array addition. Cytologically, PDE begins at the 12-cell stage of embryogenesis, is synchronous across all chromosomes, and is characterised by the formation of transient micronucleus-like bodies. Together, these findings reveal unusually extensive PDE in a free-living nematode and raise the possibility that germline-restricted repeat architecture may contribute to the atypical sex chromosome inheritance observed in *Auanema*.

## Introduction

In bilaterian animals, the germline is the sole cell population that transmits its genome to the next generation ^1^. The germline stem cell population is usually set aside very early in development, and is protected, genetically and physiologically, from influence from the soma. The soma is dispensable and can undergo genetic change without affecting the germline genome. This possibility of non-heritable change in somatic cells has been exploited widely in evolution. For example, human tissues include cells without nuclear genomes (i.e., erythrocytes) and cells where programmed genome rearrangement and mutagenesis are used to increase variation (i.e., the B-cell, or antibody, and T-cell receptors, which are generated by specific programmed rearrangement and hypermutation ^2–4^). Somatic mutation and genome rearrangement are also hallmarks of disease, especially cancers ^5^, demonstrating that even in the disposable soma, genome rearrangement needs to be precisely controlled to be useful. Ultimately, such developmentally-regulated, lineage-specific rearrangements and losses are controlled by the germline genome, which is maintained intact.

The germline hypothesis of inheritance as formulated by Weissman ^1^ was based on a large corpus of observations, including the remarkable case of early embryogenesis in a parasitic nematode of horses, *Parascaris univalens* ^6^. Boveri had noted in dividing embryos of *P. univalens* that in a subset of cells in the early embryo, the large “nuclear filaments” (the single chromosome pair of *P. univalens*) appeared to break up, yielding two sorts of fragments, one set of which was not incorporated into daughter nuclei and was subsequently lost. Only in one set of cells, the lineage that gave rise to the germline, was the single chromosome pair kept intact. Programmed DNA elimination (PDE) was also described in related nematodes, and modern genomic analysis has defined not only the full sequence of the genomes of *P. univalens* and other ascaridids ^7^, but also identified the DNA that is eliminated in the somatic cells ^7–9^. In *P. univalens,* all germline telomeres are removed, and cuts are made within the chromosome to generate 36 somatic chromosomes (n_s_=36) from the single germline one (n_g_=1). A similar pattern of PDE is found in the ascaridids *Ascaris suum* (n_g_=24, n_s_=36)*, Ascaris lumbricoides* (n_g_=24, n_s_=36) and *Toxocara canis* (n_g_=18, n_s_=36) ^8^. In these species, the sites of breakage of the germline chromosomes are inexact ^10,11^, and individual blastomeres in the embryo heal their somatic chromosomes by adding new arrays of telomeric repeat to sites that range across a kilobase or more.

PDE was recently discovered serendipitously in a free living nematode, *Oscheius tipulae* ^12^. *O. tipulae*, from suborder Rhabditina, is only distantly related to ascaridid nematodes (in suborder Spiruromorpha). The germline genome of *O. tipulae* is 60 Mb, and 0.6% of the germline genome is eliminated in somatic cells through specific elimination from all chromosome ends. Unlike in the ascaridids, there is no chromosome-internal breakage in *O. tipulae*, meaning that the somatic (n_s_) and germline (n_g_) karyotypes remain the same. The sites of elimination in *O. tipulae* are precise (the new telomeric repeat arrays are added to the same sites in all somatic cells) and are defined by a palindromic sequence motif that is required for elimination ^12,13^. PDE has also been identified in additional genera of rhabditine nematodes. In *Mesorhabditis belari,* the somatic genome is both significantly smaller (about 67% of the germline genome) and differently organised (n_g_=10 while n_s_=20), and is associated with a bipartite sequence motif ^14^. In three *Caenorhabditis* species, PDE at specific sites also trims chromosome ends and breaks chromosomes in somatic cells (such that n_g_<n_s_) ^15^.

The identification of the presence of PDE by microscopy of early embryogenesis in ten additional genera of free-living rhabditid nematodes ^16^ and the accidental discovery of PDE in *O. tipulae* during genome sequencing ^12^ inspired us to revisit the genomes of other species to explore pattern and process in the evolution of this fascinating developmental mechanism. We focused on the free-living nematode genus *Auanema,* which is relatively closely related to *Oscheius* and contains species notable for their unusual life histories. For example, *Auanema rhodense* has three sexual morphs: males, females and hermaphrodites ^17–19^ and exhibits non-Mendelian X chromosome segregation ^20^. Sex determination is partially genetic (females and hermaphrodites are XX, males are XO) and partially maternal (the decision between hermaphrodite and female sexes is conditioned by the mother’s age) ^21^. During spermatogenesis, the X chromosomes segregate equationally rather than reductionally during meiosis I, and following meiosis II, the cell division products lacking X chromosomes become residual bodies and are discarded, so that XO males produce exclusively X carrying sperm. Further, sperm produced by hermaphrodites always carry two copies of the X chromosome, while the oocytes contain none. This atypical pattern of X-chromosome transmission in the hermaphrodite is accompanied by a lack of recombination and thus the X chromosomes only recombine in females ^22,23^.

Here, we have used Hi-C and PacBio HiFi long-read sequencing to reveal that *Auanema rhodense* undergoes extensive PDE, and that the previously published assembly ^22,23^ represented only the 60 Mb somatic portion of a 180 Mb germline genome. The vast majority of the eliminated DNA is made up of huge families of tandem repeat sequences that are coordinately eliminated from somatic cells beginning at the 12 cell stage of embryogenesis, and the elimination process results in a change in karyotype from 7 to 14 chromosome pairs in somatic cells. PDE in *A. rhodense* superficially resembles that of ascaridids in that most eliminated DNA is repetitive and chromosome complement is altered, but it differs in that very few protein-coding genes are eliminated, implying that regulation of embryonic gene expression ^9^ is not the *raison d’être* for PDE in *A. rhodense*. Moreover, elimination sites are precise, with breakage and new telomere addition occurring within a palindromic sequence motif that is likely homologous to that found at elimination sites in *O. tipulae,* indicating a common evolutionary origin of PDE in *A. rhodense* and *O. tipulae* despite major divergence in the amount of DNA eliminated and the effects of PDE on somatic karyotype. Our findings suggest that acquisition of a PDE mechanism can have very different consequences for evolution of the structure and composition of germline genomes. We further speculate that expansion of repeat blocks during *A. rhodense* evolution may have enabled the unusual modifications of the meiotic program and extreme forms of meiotic drive observed in this species.

## Results

### Improved *Auanema rhodense* genome assembly and annotation

We *de novo* sequenced the genome of the *A. rhodense* APS4 inbred line ^23^ using PacBio HiFi long reads and Illumina sequencing of Arima chromatin conformation capture (Hi-C) libraries (Figure 1A; Table S1). We used DNA from staged cultures enriched for young adults to increase coverage of germline DNA. The new assembly spans 180.9 Mb, nearly three times the span of the previous assembly ^23^ (Table 1). The new assembly is also substantially more contiguous, with a contig N50 of 16.41 Mb, compared with 0.11 Mb for the previous assembly. Moreover, 92% (165.5 Mb) of the sequence was assembled into seven chromosome-level scaffolds, corresponding to the number of chromosomes defined by genetic linkage mapping and by cytological analyses of meiotic nuclei ^23^. The 11 Mb of assembled sequence that was not assigned to chromosomes was very repeat rich, and this feature likely explains the inability of the short-read Hi-C data to place it unequivocally. We confirmed the mapping of the ancestral linkage groups of rhabditid nematodes (Nigon elements) ^12,23^ on the *A. rhodense* chromosomes (Figure 1C).

**Figure 1.**
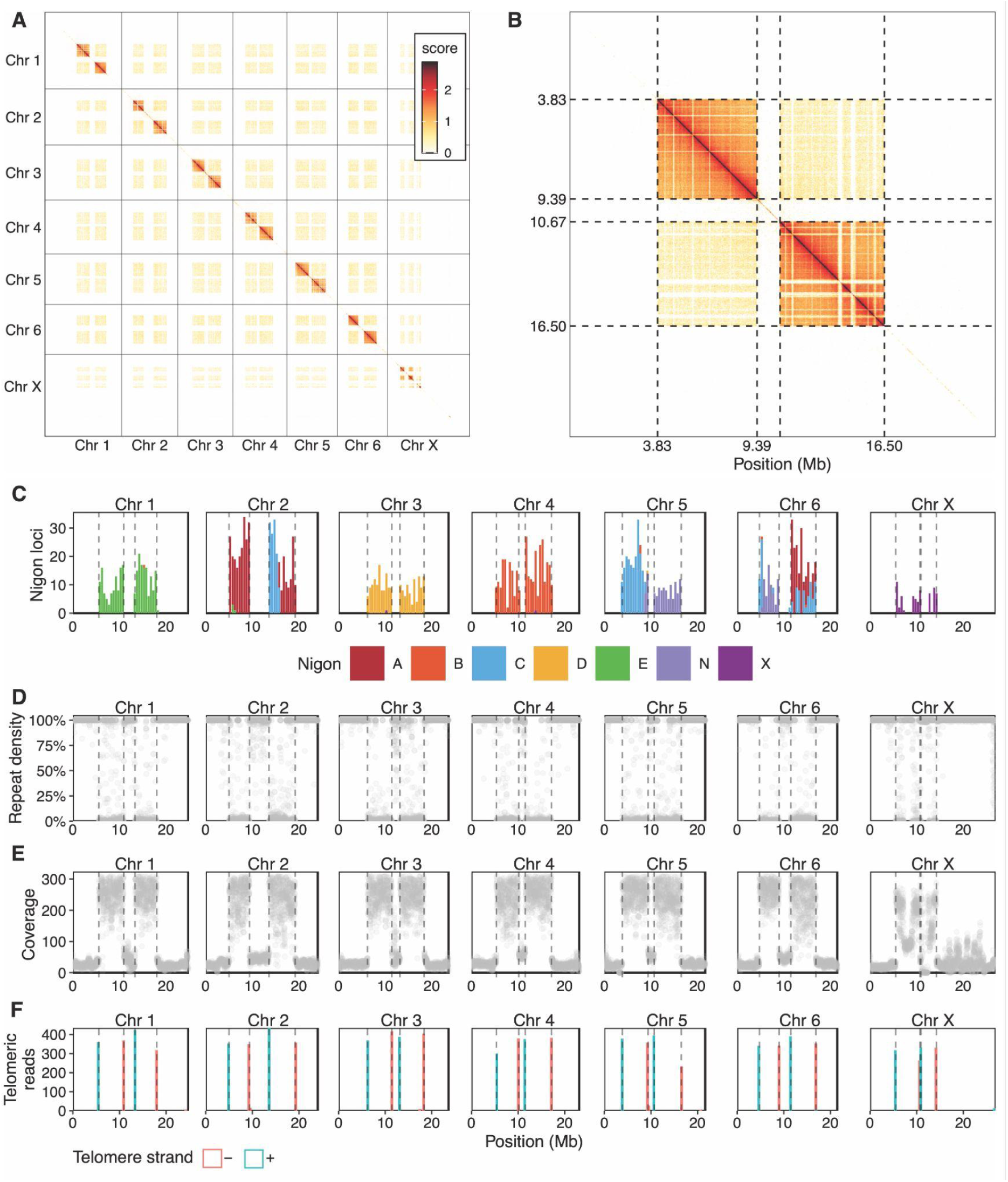
Features of the *Auanema rhodense* germline genome and evidence for PDE. **A.** Genome-wide wide Hi-C interactions at 32 kb resolution of the chromosomal scaffolds of the *A. rhodense* genome assembly. Multi-mapping reads were excluded. Grid lines delineate each scaffold. Color scale indicates interaction frequency (log scale). **B.** Detailed view of Hi-C interaction map of chromosome 5, at 32 kb resolution. Dotted lines indicate the chromosome-internal sites of telomere repeat array addition. **C.** Mapping of conserved single copy loci that have been previously allocated to the seven ancestral linkage groups of Rhabditid nematodes, Nigon elements ^12,23^, summarised per 500 kb window along each chromosome. **D.** Repeat density per 10 kb window along each chromosome (repetitive sequences identified by Red ^25^). **E.** Genome coverage supported by HiFi reads aligning uniquely with a mapping quality score of at least 10 along each chromosome. **F.** Number of mapped reads containing arrays of the telomeric repeat hexamer along each chromosome. Red bars represent mapped reads with the telomeric repeat array that has been softmasked to the right of the mapped portion of the long read while blue bars indicate those with the telomeric repeat array to the left. The absence of reads at scaffold termini reflects the fact that the assembly is not telomere-to-telomere; reads carrying telomeric repeats are therefore not expected to map to scaffold edges. The exception is a small number of reads mapping to the right-hand end of the chromosome X scaffold, suggesting this terminus approaches the true chromosome end.

**Table 1.**
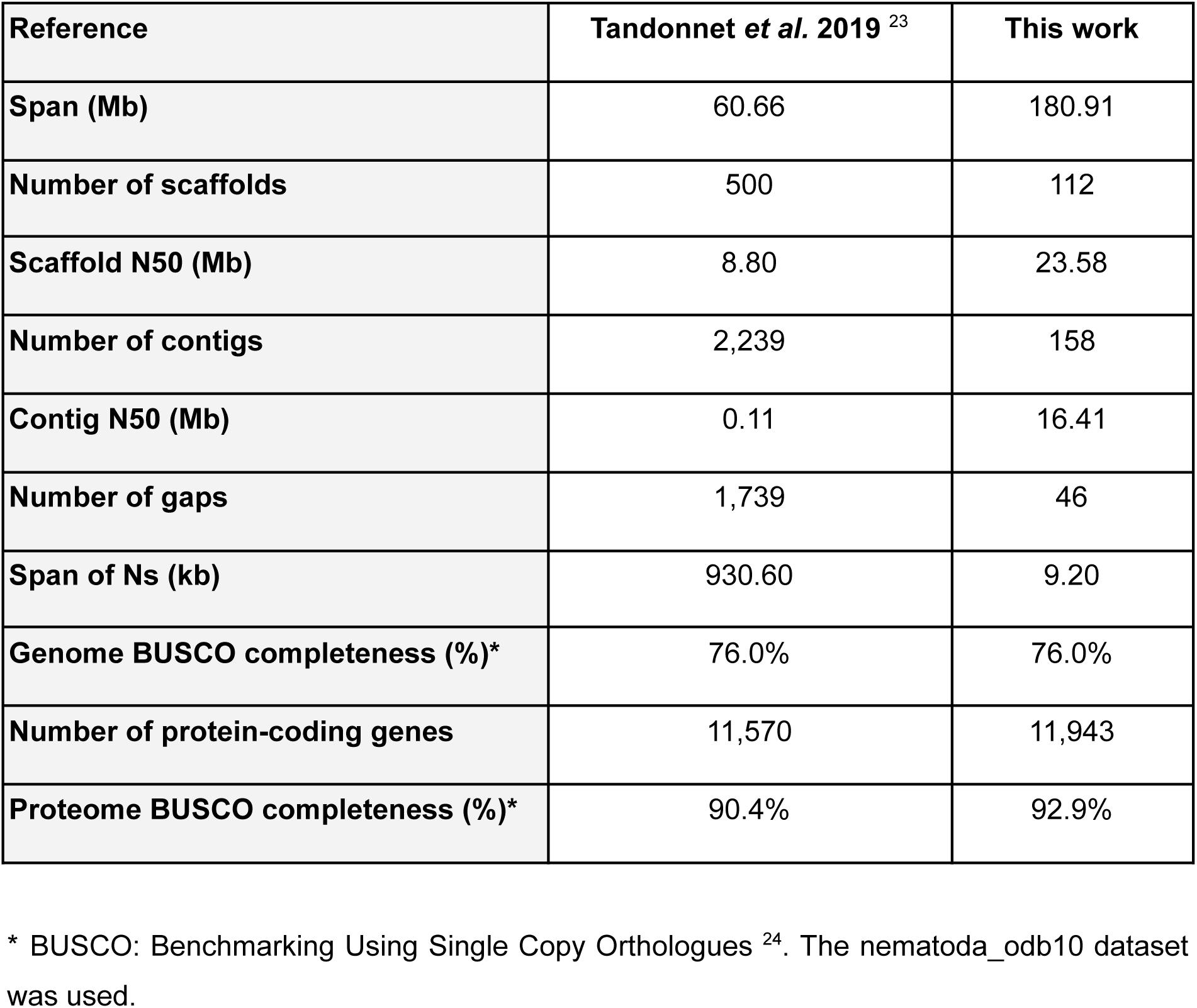
Genome assembly and annotation metrics of *Auanema rhodense*.

| Reference | Tandonnet *et al.* 2019 <sup>23</sup> | This work |
| --- | --- | --- |
| Span (Mb) | 60.66 | 180.91 |
| Number of scaffolds | 500 | 112 |
| Scaffold N50 (Mb) | 8.80 | 23.58 |
| Number of contigs | 2,239 | 158 |
| Contig N50 (Mb) | 0.11 | 16.41 |
| Number of gaps | 1,739 | 46 |
| Span of Ns (kb) | 930.60 | 9.20 |
| Genome BUSCO completeness (%)* | 76.0% | 76.0% |
| Number of protein-coding genes | 11,570 | 11,943 |
| Proteome BUSCO completeness (%)* | 90.4% | 92.9% |

\* BUSCO: Benchmarking Using Single Copy Orthologues <sup>24</sup>. The nematoda\_odb10 dataset was used.

Despite the three-fold larger size of the new genome assembly, we annotated a total of 11,943 protein-coding genes, only a slight increase from the 11,570 protein-coding genes predicted in the previous assembly (Table 1). We were able to assign functional annotation to 11,259 genes through eggNOG-mapper or InterProScan. A majority of the genes (10,136 genes; 85%) have at least one homolog in *C. elegans*, *As. suum*, *Oscheius* species or *Homo sapiens*. A total of 11,139 (93%) of the genes had expression levels of at least one transcript per million in existing RNA-seq datasets ^23^. The predicted proteome was 2.5% more complete, as assessed by the Benchmarking Using Single Copy Orthologues ^24^ (BUSCO) nematoda_odb10 dataset, than the previous ^23^ annotation (Table 1). In addition to protein-coding genes, we annotated 23,689 non-coding RNAs, with the majority (19,554) being tRNAs. Only 215 of the tRNA loci were classified as pseudogenes.

### Evidence for extensive Programmed DNA elimination in *Auanema rhodense*

The Hi-C interaction map of the *A. rhodense* assembly showed a striking checkerboard pattern, reflecting both *cis*- and *trans*-chromosomal Hi-C interactions (Figure 1A, B). In this analysis, Hi-C reads that mapped to multiple locations in the genome were excluded, and thus regions of the genome that are rich in repeats appear to have low interaction frequency in the Hi-C interaction map. Each autosome had three regions with low apparent interaction frequency, one in the centre and one at either end (Figure 1A, B), whereas the X chromosome had two large low apparent interaction regions, one at each end, and two smaller regions in the middle of the chromosome. This pattern, i.e. lack of signal in otherwise good Hi-C libraries mapped to exclude multi-mapping data, is strongly indicative of high repeat content, and these regions were indeed exceptionally repeat-rich (typically >90% per 10 kb window) (Figure 1D). There were sixteen distinct, non-repetitive regions where repeat density was typically <10% in each 10 kb window, all of which had dense *cis*- and *trans*-chromosomal interactions (Figure 1A, B). Each autosome contained two of these regions, while the X chromosome had four (three in the centre of the chromosome and one at the right end). The vast majority of protein-coding genes were located within these non-repetitive regions (Figure 1C, Figure 2A).

**Figure 2.**
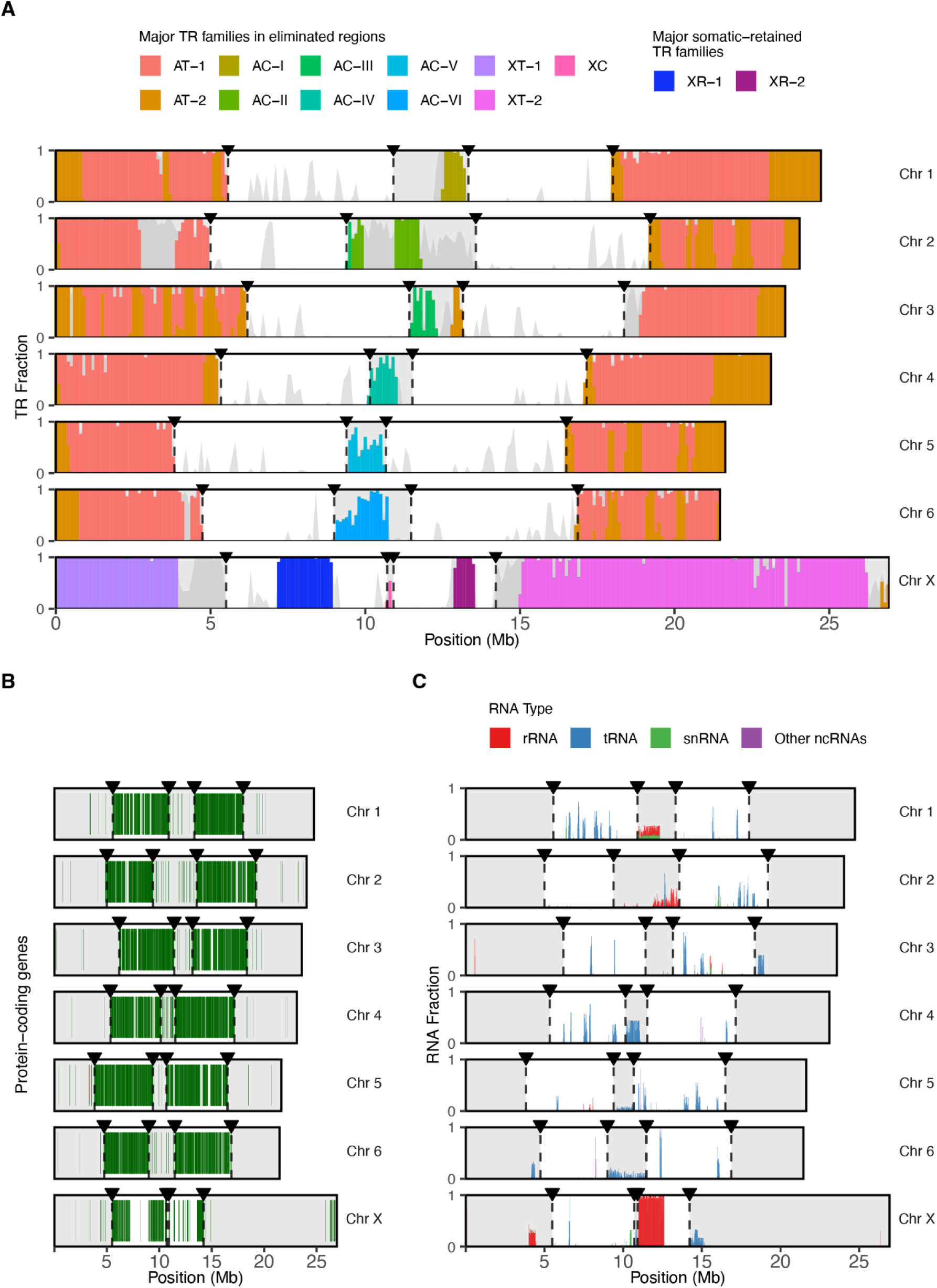
Chromosome-wide distribution of tandem repeat blocks, protein-coding genes and other features. **A.** Distribution of the dominant tandem repeat (TR) families (fractional coverage per 100 kb; total TR coverage in grey). rRNA loci are masked to avoid redundancy with panel C. TR names reflect genomic location and elimination fate: autosomal terminal (AT-1/2), autosomal central (AC-I–VI), X-terminal eliminated (XT-1/2), and X-somatic retained (XR-1/2). **B.** Distribution of protein-coding genes (dark green lines). **C.** Genome-wide distribution of RNA genes (tRNA, rRNA, snRNA, and other ncRNA) shown as stacked fractional coverage per 10 kb window. Bar height represents total rRNA occupancy of the window and segment heights show the contribution of each class. In all panels, germline-restricted regions are highlighted with semi-transparent grey overlays. Boundaries between germline-restricted and retained regions are indicated by dashed vertical lines with inverted black triangular markers (▼) at the top. Black rectangles outline each chromosome.

Long read coverage also varied along chromosomes, largely reflecting the repeat-rich and gene-rich regions described above (Figure 1E). This was not because reads could not be mapped to the repeat-rich regions, as the high accuracy of the PacBio HiFi data combined with small-scale variation in the repeat sequences meant that the majority of long reads could be uniquely mapped. The gene-rich regions had an average coverage of 236-fold, while most repeat-rich regions had only 29-fold coverage (Figure 1E). Consistent with the X chromosome being hemizygous in males, the three central gene-rich regions of the X chromosome displayed lower coverage than autosomes (an average of 146-fold). The repeat-rich regions at either end of the X (including the non-repetitive segment at the right end) had an average male coverage of 29-fold, higher than expected, likely because of repetitive content. The X had an additional region of low coverage in the centre that was not particularly repeat-rich (Figure 1D, E).

The consistent difference in coverage in different segments of the chromosomes was reminiscent of the pattern observed due to PDE in *O. tipulae* ^12^, raising the possibility that the low-coverage, repeat-rich regions represent DNA that is eliminated from the *A. rhodense* genome via PDE. To investigate this, we searched for evidence of apparent sites of telomere repeat addition within the germline chromosomes by aligning reads containing telomeric repeat arrays to the assembly (Figure 1F). We identified four sites of telomeric repeat addition within each of the six autosomes, all of which coincided with the transition between the high-coverage, gene-rich regions and the low-coverage, repeat-rich regions (Figure 1D-F). We also found four telomeric repeat addition sites within the X chromosome. Two of these sites were at the transitions between the repeat-rich, low-coverage regions on the chromosome, as observed in the autosomes. The other two bracketed a low-coverage non-repetitive region in the centre of the X. All of the 11 Mb of the assembly that we were not able to unequivocally assign to chromosomes had low coverage, suggesting that it too was eliminated from the soma. Together, these data strongly suggest that *A. rhodense* undergoes PDE, eliminating approximately 61% of its genome from somatic cell lineages (Table S2).

### Distinct large arrays of tandem repeats in the autosomes compared to the X chromosome

We identified mobile element-derived, interspersed and tandem repeats (TRs) in the germline genome. The large number of different TR sequences found were clustered at 80% identity to define TR types. The majority of the putatively germline-restricted sequences are repetitive, comprising large arrays of TRs with unit repeat sizes ranging from 167 to 504 nucleotides in length (Figure 2A; Table S3). The multimegabase TR blocks located at both ends of the autosomes are composed almost entirely of two autosome-specific TRs that are restricted to these regions. In addition to these terminal TR blocks that are shared by all 6 autosomes, the germline-restricted sequences in the centres of the germline autosomes each contain a unique set of TRs that are largely absent from the rest of the genome.

The X chromosome has a distinct pattern of TR distribution. Two-thirds of the X chromosome is made up of TR blocks. In contrast to the autosomes, the left and right TR blocks on X differ substantially from each other in size, and each has a unique composition of X-chromosome-specific TRs. In addition, whereas the autosomes each have a central TR block that is eliminated from somatic cells, the very small eliminated region at the center of the X chromosome lacks large TR blocks, while the two somatic chromosomes derived from the X chromosome both retain large TR blocks.

While the vast majority of germline-restricted TRs are satellite-type repeats, some TRs correspond to tandemly repeated blocks of elements with known functions, including arrays of 5S ribosomal RNA (rRNA), several transfer RNA loci (tRNA), and U2 spliceosomal RNA loci (Figure 2C; Table S4). The density of these non-coding RNA loci (always <20% per 10 kb window) contrasts with the density of satellite TRs (typically covering >90%).

### Few protein-coding genes are eliminated through PDE

Despite PDE eliminating nearly two thirds of the germline genome span, only 3% (360) of the protein coding genes (PCG) reside within the eliminated DNA (Figure 2B; Table S5). Most (93%; 334) of these eliminated PCGs have likely homologues in other nematode species. Half of the eliminated PCGs carry domain annotations associated with transposable element (TE) functions, including reverse transcriptases, polymerases, and helitrons, and 141 (39%) are overlapped by TE annotations (compared to only ∼0.9% of genes in the retained somatic genome) suggesting that most annotated PCGs in the eliminated DNA represent TE-derived sequences. A relatively small fraction of these 360 eliminated PCGs are expressed, judged by read mapping from published transcriptome data ^23^. Only 95 had >0.05 transcripts per million in any RNA-seq dataset, and just ten exceeded 3 transcripts per million in at least one sample.

PDE rarely eliminated a gene family entirely from the somatic genome. The 133 PCGs whose entire orthogroup is eliminated are dominated by TE-derived sequences and a divergent, *Auanema*-specific histone H3 variant cluster distinct from that of the core histone orthogroup retained in the soma. The two most biologically notable cases of eliminated genes with somatically-retained paralogs are a single actin B paralog (one of seven actin copies in the genome, the remaining six of which are somatically retained) and MRE-11, a core component of the double-strand break repair machinery, whose somatic paralog is located immediately outside the eliminated sequence on chromosome 6. Among the other expressed non-TE eliminated PCGs with somatic paralogs are an uncharacterised but conserved glutamine-rich protein orthologous to *C. elegans brp-1*, an imidazolonepropionase, an aldo-keto reductase, and six tandemly duplicated copies of a GIT2 ortholog, related to homologs involved in gonad morphogenesis in *C. elegans*. Twenty-six eliminated PCGs had no detectable orthologs in other species and had no paralogs within the *A. rhodense* somatic genome. Most were transcriptionally silent and had no functional annotations, but one, nxAuaRhod_g826, located at the boundary of an eliminated region on chromosome 6, was highly expressed and had no homology to any known gene or domain. These analyses suggest that PDE is unlikely to be used as a significant mechanism for controlling gene expression between the germline and the soma in *A. rhodense*.

### Sequence motifs are found at chromosome break sites

Given the precision of the boundaries that demarcate the switch between germline-restricted and retained genome regions (Figure 3B), we sought to identify *cis*-regulatory elements close to the chromosome breakage positions. We found a sequence motif, or Sequence For Elimination (SFE), that overlaps all 28 internal break sites in the *A. rhodense* germline sequence. This palindromic motif is 29 bases long, formed from two, relatively conserved 14-base sequences and a central base with low sequence conservation, which is flanked by four highly conserved residues. Telomere addition occurs in the centre of the SFE, between positions 14 and 15. The *A. rhodense* SFE is strikingly similar to the *O. tipulae* SFE (Figure 3A): both motifs are palindromic, flanked by AT-rich regions, and share clear sequence similarity (Figure 3A). We note that the 29 base *O. tipulae* SFE predicted here differs from one previously proposed ^13^.

**Figure 3.**
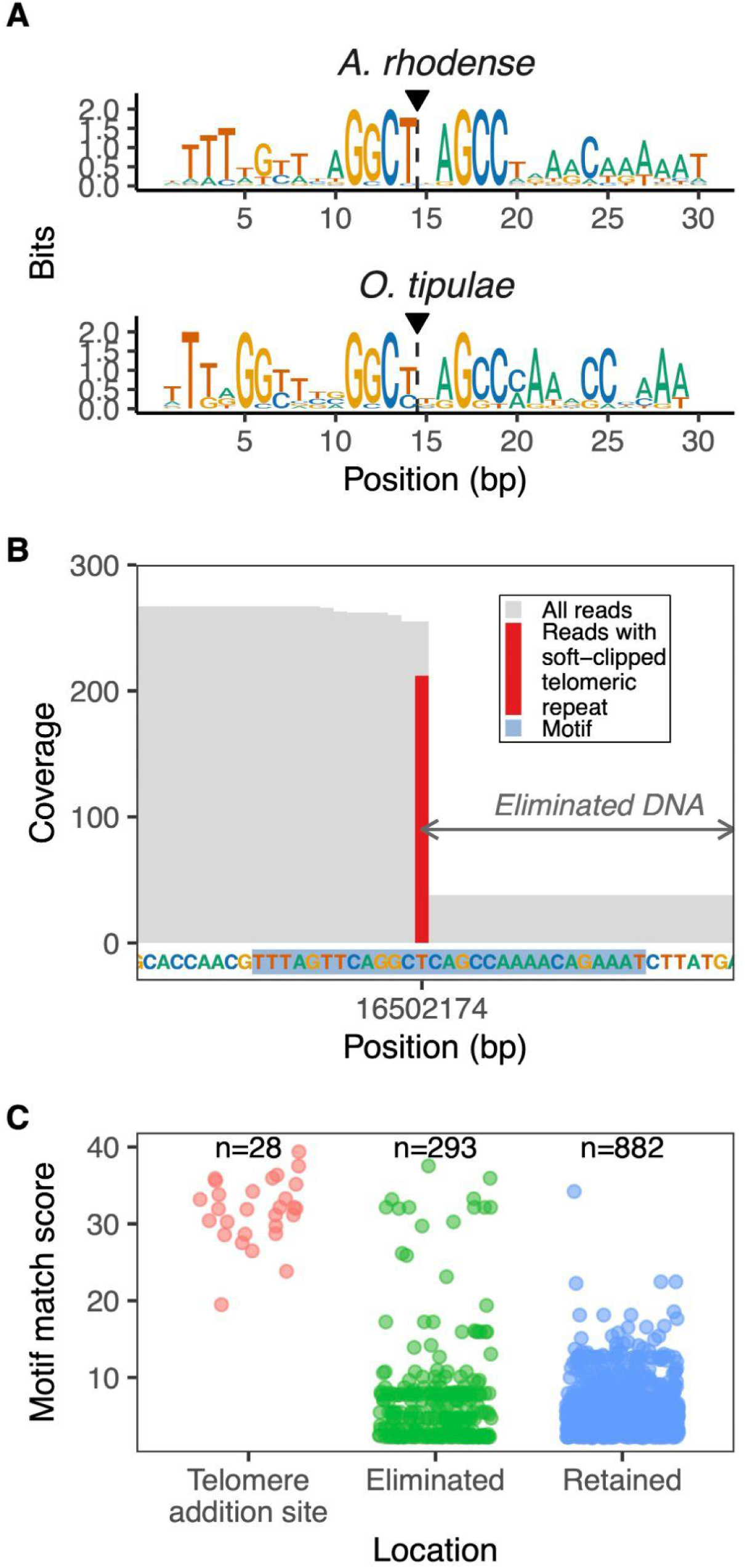
New telomeres define chromosome break sites in the genome of *Auanema rhodense*. **A.** The Sequence for Elimination (SFE) associated with telomere addition sites in *A. rhodense* compared to the *O. tipulae* SFE. The SFEs have been aligned to highlight their similarity. Vertical dotted lines, capped with a triangle (▾), mark the nucleotide position at which telomeric repeat=clipped read coverage across the boundary drops sharply in sequencing data; we interpret this coverage transition as the inferred nuclease cleavage (nucleation) site that initiates elimination of the flanking DNA. **B.** Long read coverage highlighting the precise position of soft clipped telomeric repeat arrays in most reads aligning on chromosome 5 around position 16,502,174. The part of the sequence matching the motif is highlighted (light blue) as well as the eliminated region. **C.** Genome-wide occurrence of the sequence motif. Eliminated regions correspond to those inferred to be lost from the soma whereas the retained regions are those inferred to be present in all cells.

To assess the specificity of the *A. rhodense* SFE, we searched the genome using a position-weight matrix constructed from the SFEs at all 28 break sites. This analysis indicated that strong motif matches were highly enriched at *bona fide* break sites. Within the ∼60 Mb of retained DNA, only four sites had similarity scores greater than the lowest scoring break site-associated SFE (Figure 3C), none of which showed any evidence of telomere addition in our data. Within the eliminated DNA, we found 15 sites that had scores greater than the lowest scoring break site-associated SFE. Seven of these were on the X chromosome, with five motifs with identical scores occurring in a single 14 kb region with a periodic spacing of 3.5 kb. Chromosomes 1, 2, and 3 each had a high-scoring motif match within 1.1 kb downstream of the break site (Table S6). For eight of the eliminated SFE sites with high scores, we observed a few reads containing telomeric repeat arrays that started at the central base of the motif, suggesting they can also undergo breakage and telomere addition, but are not retained in the soma because they lie within DNA that is eliminated because of another site at the true somatic chromosome end.

### Visualizing programmed DNA elimination in early *A. rhodense* embryos

To directly confirm that PDE reshapes the somatic genome in *A. rhodense*, we used immunofluorescence microscopy and fluorescence *in situ* hybridization (FISH) of sequences predicted to be eliminated in early cleavage stage embryos (Figure 4). Embryos were co-stained with DAPI to show DNA, antibodies against the repressive histone mark H3K9me3 to visualize chromatin and anti-α-tubulin antibodies to visualize microtubules. In embryos during or after the first three cell division cycles (Figure 4A), DAPI and H3K9me3 are detected only in nuclei, on mitotic chromosomes, and in polar bodies (PB, discarded products of the oocyte meiotic divisions). However, beginning at the 12-cell stage, we detected intense DAPI signals, typically associated with H3K9me3, that did not correspond to nuclei, mitotic chromosomes, or PBs. As these micro-nucleus-like (MN-like) bodies are reminiscent of those observed in other systems during PDE ^8,14^, we interpret their presence as indicative of the onset of PDE.

**Figure 4.**
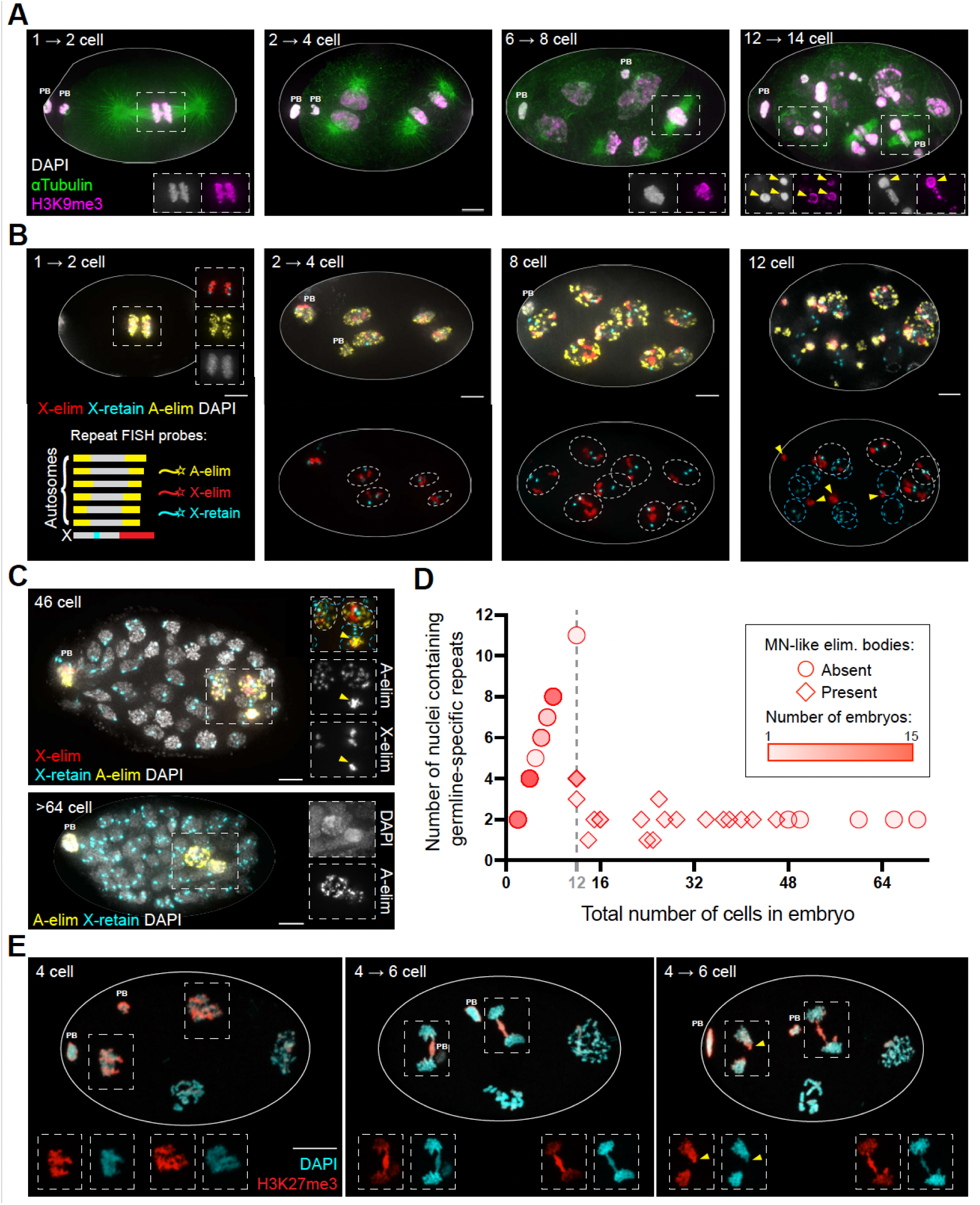
Visualizing programmed DNA elimination in early *A. rhodens*e embryos. **A.** Immunofluorescence images of embryos from the first mitosis through the 12 cell stage, stained for Tubulin, H3K9me3, and DNA; PB indicates polar bodies when they are visible. Insets show individual channels for DAPI and H3K9me3 for the indicated fields; yellow arrowheads indicate micronucleus-like (MN-like) elimination bodies, which first appear at the 12 cell stage following divisions of 4 of the 8 blastomeres present at the 8 cell stage. Scale bar = 5 µm. **B.** Fluorescence in situ hybridization (FISH) images of embryos from the first mitosis through the 12 cell stage; schematic indicates genomic regions targeted by repeat FISH probes represented by the indicated colors. In the lower row, white dashed ellipses indicate nuclei where the X (and autosomal) repeats are still present, while blue dashed ellipses indicate post-elimination nuclei. MN-like elimination bodies are present in the 12 cell embryo; yellow arrowheads indicate a subset that contain eliminated X repeats. Scale bar = 5 µm. **C.** FISH images of later stage embryos, showing that eliminated (red and yellow) repeats are absent from most nuclei and are still present in only two nuclei. Scale bar = 5 µm. **D.** Progression of DNA elimination in the early embryo (N = 74 embryos), illustrating that DNA repeats that will eventually be eliminated from somatic cells are present in all nuclei through the 8 cell stage, and that the number of nuclei containing these repeats subsequently plateaus at two as cell divisions proceed. Circles represent embryos where no MN-like bodies were present, diamonds represent embryos with these bodies. Darker colour shading reflects multiple scored embryos with a given pair of x.y values, as indicated. Dashed line indicates the 12-cell stage where DNA elimination and MN-like bodies are first detected. Points on graph were pooled from multiple FISH experiments, each using one or more FISH probes corresponding to tandem repeats destined for elimination (see Methods for probe sequences). Embryos with unresolvable or ambiguous nuclei were not scored. **E.** Immunofluorescence image of early pre-elimination embryos between the four and six cell stages, showing a lagging subset of DNA enriched for H3K27me3. Insets show individual channels for DAPI and H3K27me3 for the indicated fields. Yellow arrowhead indicates lagging DNA remaining associated with a nucleus post-division. Scale bar = 10 µm.

FISH analysis further illuminated the timing and process of PDE (Figure 4B-D). For these experiments, we used probes designed to detect X-chromosome repeat sequences eliminated during PDE (X-elim, corresponding to repeat XT-2; Figure 2A), Autosomal repeat sequences eliminated during PDE (A-elim, corresponding to repeat AT-1, present on all autosomes; Figure 2A), and/or X-chromosome repeat sequences retained in all cells (X-retain, corresponding to repeat XR-1; Figure 2A). All three FISH probes were detected in all embryonic nuclei through the 8-cell stage, indicating that no PDE had occurred during these early rounds of division (Figure 4B, left three panels). Beginning at the 12-cell stage, both X-elim and A-elim FISH signals were absent from a subset of nuclei and were concurrently detected in MN-like bodies, providing direct evidence that PDE had been initiated (Figure 4B, right panel). Both X-elim and A-elim were often present within the same MN-like body, indicating that these bodies can contain DNA derived from multiple chromosomes. In the majority of 12-cell embryos scored, all FISH probes were detected within four of the nuclei, while only the somatically-retained X-retain probe was detected in the remaining eight nuclei. Post-elimination nuclei in 12-cell embryos were also distinguishable by having faint DAPI signals, consistent with their abrupt decline in DNA content.

This “DAPI-faint” phenotype resolves by the time embryos reach 30 cells, with post-elimination nuclei in >30-cell embryos once again easily visible by DAPI staining. Notably, in >30-cell embryos, the vast majority of cells contain post-elimination nuclei, MN-like bodies are few or absent, and the number of nuclei that still contain germline-specific repeats has plateaued at two (Figure 4C-D). Given that all repeats are present in the *A. rhodense* germline genome, the retention of these repeats can serve as a marker of primordial germ cells (PGCs), and we thus infer that *A. rhodense* has two PGCs. Further, we note that X-elim and A-elim probes consistently behaved concordantly, with either both FISH signals or neither being detected in a given nucleus. As we did not detect evidence for a “partial elimination” state in which only a subset of the elimination-destined repeats had been removed, we infer that PDE for different chromosomal regions occurs in a coordinated manner.

Immunofluorescence images of early embryos stained for H3K27me3 and DNA provided additional insight into the PDE process (Figure 4E). Specifically, we detected lagging DNA in anaphase-dividing cells in a subset of embryos transitioning from the 4- to 6-cell stage. Further, this lagging DNA was enriched for H3K27me3. Notably, this H3K27me3-enriched lagging DNA was observed during the cell cycle preceding the cell cycle when DNA elimination first became evident, as our FISH analysis did not detect evidence of DNA elimination until after the 8-cell stage. We speculate that the H3K27me3-enriched lagging DNA may represent DNA that is being prepared for elimination during the following cell cycle.

## Discussion

There is a tendency to view eukaryotic genomes as constant, with most or all cells in an organism sharing an identical genome except for new mutation, and many mechanisms operating to ensure that the genome is faithfully inherited at each cell division. Here we present an extreme counter-example. Nearly two thirds of the genome of the free-living nematode *Auanema rhodense* is eliminated from somatic cells during early embryogenesis, leading to dramatically different germline and somatic genomes. The eliminated DNA consists primarily of massive TR arrays that occur in the middle and at each end of all seven chromosomes of the germline genome. We identified conserved sequence motifs at the elimination boundaries that coincide with sites of DNA breakage and new telomere addition, defining the ends of the 14 chromosomes that make up the greatly diminished somatic genome.

Comparing features of PDE in *A. rhodense* and the related rhabditine nematode *O. tipulae* reveals several commonalities, as well as important distinctions. In both species, PDE was first detected cytologically following the 8-cell stage of embryogenesis (by DNA and histone-modification staining and TR-probe FISH in *A. rhodense* and by telomere FISH in *O. tipulae*), suggesting that PDE occurs with similar timing ^7,13–15^. Whereas DNA staining detected prominent micronucleus-like elimination bodies during PDE in *A. rhodense*, these were not detected in *O. tipulae*, likely reflecting differences in the mass of DNA eliminated (see below).

The *A. rhodense* and *O. tipulae* elimination sites have two striking similarities. First, the sites of elimination in both species are precise, with almost all telomere-repeat-associated reads supporting a single site of telomere addition for each new chromosome end. Because elimination begins during or after the 8-cell stage, we can infer that the same site is used independently in each somatic blastomere. Precise elimination sites are also observed in *Caenorhabditis* species that undergo PDE ^15^, suggesting that this is likely to be a general feature of PDE in rhabditine species. The second clear similarity between *A. rhodense* and *O. tipulae* elimination sites is the presence of a conserved, 29 base palindromic motif. The motifs presumably serve as binding sites for the as-yet-unidentified elimination machinery. Consistent with this, Dockendorff et al. showed that editing the sequence of the motif using CRISPR prevented elimination from occurring at an edited site in *O. tipulae* ^13^. The similarity between the *A. rhodense* and *O. tipulae* motifs suggests that their PDE mechanisms have a common ancestor, which is consistent with the phylogenetic distribution of PDE unveiled in a recent cytological survey ^16^. Sequence motifs are also associated with the sites of PDE in *Caenorhabditis*, but their sequence and structure are notably different ^15^. The base-level specificity and presence of sequence motifs differentiate PDE in rhabditine species from that in ascaridids, where telomere addition occurs variably within 1-2 kb regions that lack sequence motifs or other identifiable features ^7^. PDE is common in rhabditine nematodes (though not universal) and additional genomic analysis of a wide range of species will be critical in resolving mode and tempo of this genetic mechanism ^16^.

The most striking difference between PDE in *A. rhodense* and *O. tipulae* is in the amount of DNA that is eliminated. While *O. tipulae* eliminates 0.6% of its genome (and PDE in the three *Caenorhabditis* species results in the discarding of 0.7-2.3% of the genome ^15^), *A. rhodense* eliminates nearly two-thirds. The bulk of this excess in *A. rhodense* is made up of TRs. In the ascaridid nematodes, PDE similarly results in the elimination of large proportions of the germline genome (7% to 90%), and the eliminated DNA is also dominated by repeats ^7^. PDE could therefore be framed as a mechanism used by *A. rhodense* and other nematodes to streamline their genomes in the soma, and to protect the soma from potentially deleterious effects of ectopic transcription or activation of mobile elements. However, we believe this puts the cart before the horse: it would be less costly to simply eliminate the repeats from the germline genome rather than evolve and deploy a complex repeat-elimination machinery in every embryo. Rather, we suggest that the eliminated regions are places in the genome where selective surveillance is weak, and thus where repeats can accumulate. In this model, repeat accumulation in eliminated regions is an epiphenomenon resulting from the prior existence of PDE. The difference among species in the amount of repeat accumulation in the eliminated regions would then be a product of the different contributions of replication slippage to genomic change in each.

Regardless of how or why the tandem repeat blocks have accumulated in the germline chromosomes of *A. rhodense*, their existence may have been exapted to new function, with particular consequences for the mechanisms of chromosome inheritance during the reproductive cell division program of meiosis. One intriguing possibility is that a subset of these *A. rhodense* tandem repeat blocks might function analogously to *C. elegans* meiotic “pairing centers” (PCs), which are specialized chromosomal domains enriched for short (11-12 bp) repeated motifs that associate with a family of rapidly-evolving, Zn-finger PC-binding proteins and play key roles in enabling homologous chromosomes to identify their appropriate pairing partners ^26–30^. While the tandem repeats that are shared among both arms of every *A. rhodense* germline autosome are unlikely to aid in homolog recognition, we speculate that the repeat blocks unique to the centers of each autosome and the X chromosome might facilitate partner choice during the process of homolog pairing. It will be interesting to determine whether there are proteins that bind specifically to the repeats that are specific to each chromosome. Related to this, it is noteworthy that there are several Zn-finger proteins encoded in the eliminated portion of the *A. rhodense* genome.

The unique repeats found on the *A. rhodense* X chromosome may also play a role in the highly unusual non-Mendelian segregation patterns, involving extreme forms of meiotic drive in both spermatocyte and oocyte meiosis, exhibited by the X chromosome during meiosis in this species ^22^. Having a unique repeat composition and chromatin structure could confer distinct properties to the X chromosomes that could contribute to their distinct behaviour. For example, special properties conferred by their unique repeat composition could potentially drive the X chromosomes to undergo equational separation at meiosis I while the autosomes undergo canonical reductional segregation, or to be preferentially retained in or discarded from the meiotic products destined to become the gametes ^22^. We speculate that repeat accumulation, and in particular, differentiation of the X chromosome from the autosomes, may have co-evolved in *A. rhodense* in tandem with both the emergence of its unusual trioecious life history and its atypical patterns of sex chromosome inheritance.

## Acknowledgements

The genomic and transcriptomic data reported in this study were produced by the long read team in Sanger’s Scientific Operations, to whom huge thanks. We thank members of the Blaxter laboratory for comments on the manuscript, and Alan Tracey for guidance in use and interpretation of GAP5 analyses. This research was funded in part by the Wellcome Trust Grant 220540/Z/20/A. Pablo Gonzalez was supported by a PhD Fellowship from the Darwin Trust of Edinburgh. A.P.-d.S. and S.A. were supported by grants from the Biotechnology and Biological Sciences Research Council (BB/Y512904/1) and The Leverhulme Trust (RPG-2019-329). B.C. was funded by the Doctoral Training from BBSRC (Midlands Integrative Biosciences Training Partnership). This work was also supported by NIH Grant R35GM126964 to A.M.V. and by NIH Grant 1S10OD01227601 from the NCRR to the Stanford Cell Sciences Imaging Facility (RRID:SCR_017787). L.G.S. was supported by NIH Training Grant T32GM007790.

For the purpose of Open Access, the author has applied a CC BY public copyright licence to any Author Accepted Manuscript version arising from this submission.

## Data Availability

All raw sequence data are available in the European Nucleotide Archive. The *A. rhodense* genome assembly is available in the ENA under accession GCA_964057225.1. For details of raw data accession numbers, please see Table S1.

## Supplementary Information

Document S1: Supplemental Tables S1 to S9.

## Methods

### Strain culture and DNA extraction

All analyses were carried out on the inbred *A. rhodense* strain APS4 ^23^. This strain was derived from the inbreeding of strain SB347, isolated from a deer tick in Rhode Island, USA ^31^. Nematodes were cultured on solid nematode growth medium (NGM) comprising NaCl (3 mg/ml), peptone (2.5 mg/ml), agar (19 mg/ml), cholesterol (5 µg/ml), CaCl (1 mMol), MgSO_4_ (1 mMol), KH_2_PO_4_ (25 mMol), and AmphotericinB (8 µl/ml). These were spotted with an overnight culture of *Escherichia coli* OP50, following standard nematode culturing procedures ^32^. Mixed stage cultures were harvested directly from plates, while for staged cultures we performed a bleach treatment to isolate eggs and then harvested the plates when most animals were adults 2-3 days later. Individual nematode cultures were rinsed from NGM plates using ice cold M9 minimal buffer (12.8 mg/ml Na_2_HPO_4_•7H_2_O, 3 mg/ml KH_2_PO_4_, 500 µg/ml NaCl, 1 mg/ml NH_4_Cl, 2 ml Glycerol (20%), 2M MgSO_4_, 0.1M CaCl_2_) and pooled in 15 ml falcon tubes. Pooled nematodes were pelleted, then washed twice with 4 ml and 10 ml ice cold M9 minimal buffer by centrifugation. To remove *E. coli* from pooled nematodes, a density gradient sucrose float method was performed using a 5:7 ratio of 60% sucrose solution to M9 minimal buffer. Nematodes were recovered from the interphase of the sucrose/M9 gradient and washed a further two times with M9 minimal buffer. The floating layer of nematodes was collected and washed in an M9 buffer. Aliquots of 100 µL were transferred with minimal supernatant to 1.5 mL LoBind Eppendorf tubes, snap frozen in liquid nitrogen, and kept at -80°C until required for extraction.

To enrich populations for adult nematodes, we bleach treated a pellet of mixed-stage nematodes to obtain eggs ^32^. After bleaching, the eggs were incubated at 25°C for 24 hours at 80 rpm. The arrested L1 larvae that hatched from these eggs were then plated on NGM with *E. coli* and left to feed until the majority reached adulthood. *E. coli* was removed by density gradient sucrose float and the resulting collection flash-frozen in liquid nitrogen and kept at -80°C until DNA extraction.

### DNA extraction and genome sequencing

For PacBio HiFi sequencing, high molecular weight DNA was extracted from 20 to 30 mg of snap-frozen nematodes using a modified version of the HMW DNA kit (Qiagen). A lysis buffer mastermix was prepared containing 200 µl PBS, 20 µl Proteinase K (Qiagen), 4 µl RNAse A (Qiagen) and 150 µl Buffer AL (Qiagen) per sample. Nematodes were homogenised in a lysis buffer using BioMasher II disposable micro tissue homogenisers (Nippi inc). Tissue lysis was performed overnight at 25°C. The remaining steps in the process were performed in accordance with the kit manufacturer’s handbook for fresh and frozen tissue extraction. DNA was sheared to ∼15 kb using a hydroshear instrument, end-repaired and processed for PacBio SMRTbell v2.0 library preparation according to manufacturer’s instructions. Purified DNA was quantified using the Qubit 2 fluorometer (ThermoFisher Scientific) and DNA fragment analyses were performed by capillary electrophoresis with a Femto Pulse system (Agilent Technologies) utilising a 165 kilobase ladder. DNA sample purity was assessed using the Nanodrop spectrophotometer (ThermoFisher Scientific). Libraries were sequenced on PacBio Sequel IIe instruments.

To generate Hi-C data, ∼30 mg pellets of frozen nematodes were processed using the Arima Hi-C version 2 kit following manufacturer’s instructions, and sequenced on one eighth of an Illumina Novaseq S4 lane using paired end 150 base sequencing.

### RNA extraction and sequencing

RNA was extracted with TRIzol® Reagent from Life Technologies (REF: 15596026). Nematodes were snap frozen in liquid nitrogen. One ml of TRIzol® Reagent was added and the samples underwent 3 freeze thaw cycles with liquid nitrogen. The RNA was extracted following the manufacturer’s protocol for RNA extraction. RNA was eluted in 100µl MagMAX^TM^ Total RNA Elution Buffer (REF: A41043). The RNA was digested with DNase using the TURBO DNA-*free*^TM^ Kit from Invitrogen (REF: AM1907). Quality control was performed with the NanoDrop One Photospectrometer (Thermo Fisher Scientific) and RNA integrity was analysed with the TapeStation 4150 (Agilent) using an RNA Screen tape. Illumina paired-end libraries were prepared and multiplexed in a HiSeq 4000 with a read length of 150 bases.

### Genome assembly and curation

We estimated the genome coverage of each dataset with GenomeScope.FK (https://github.com/thegenemyers/GENESCOPE.FK) ^33^ by counting kmer frequencies with FastK (https://github.com/thegenemyers/FASTK). To generate primary assemblies, we removed reads that started or ended with more than three copies of the telomere repeat (TTAGGC if at the end of the read or its reverse complement if at the start of the read) with a custom python script. The readsets depleted of telomeric reads were then assembled with Hifiasm (v0.16.1) ^34^ adjusting the `--hom-cov` according to the estimated coverage. We removed haplotigs and heterozygous overlaps from these assemblies with purge_dups ^35^ providing cutoff values inferred from the kmer frequency plot. We used BlobToolkit (v1.3.4) ^36^, to identify for removal contaminant bacterial sequences from these purged assemblies. The mitochondrial genome was identified and annotated with MitoHifi ^37^ using as reference the *C. elegans* mitochondrion genome (NC_001328.1). The nuclear genome was scaffolded with YaHS, in `no_break` mode, for scaffolding ^38^. Finally, each assembly was analysed and manually improved using rapid curation ^39^. The chromosome level scaffolds were named according to sequence size with exception of the one corresponding to the X chromosome.

### Repeat and non-coding element genome annotation

We re-annotated the *Oscheius tipulae* CEW1 (GCA_013425905.1) assembly. Long terminal repeats (LTR) were identified with the LTRdigest pipeline ^40^. Briefly, we identified candidates with genometools (v1.6.1; https://github.com/genometools/genometools), and kept only regions that had similarity to LTR proteins as assessed by hmmsearch (HMMER v3.3.2) ^41^ against the protein profiles from LTRdigest. Transposons were identified in each genome by running TransposonPSI (v1.0.0; https://transposonpsi.sourceforge.net/) in nucleotide mode. A wider diversity of interspersed repeat elements was predicted by running RepeatModeler (v2.0.1) ^42^ with the `-ltr` flag to include the prediction of LTR elements. To create a comprehensive library of repeats, we combined the nucleotide sequences from these predictions with the interspersed repeats previously deposited in Dfam_3.0 ^43^ and associated with Nematoda. We reduced the redundancy of this repeat library by collapsing all sequences that had >= 80% identity with each other into a single representative sequence. We used vsearch (v2.21.1) ^44^ to perform this sequence clustering. This custom non-redundant library was classified by RepeatModeler’s utility tool, RepeatClassifier. To remove from our library high confidence proteins predicted from high copy number repeats, we searched this non-redundant library against the proteomes of *C. elegans* (PRJNA13758) and *P. pacificus* (PRJNA12644), which were downloaded from Wormbase Parasite ^45^, using tBLASTn ^46^. We removed repeat sequences that had hits to the reference proteomes with an e-value < 0.0001 and which could not be classified by RepeatClassifier. This custom library was given to RepeatMasker (v4.1.2-p1) issued with flags `-xsmall -gff` to identify repeat instances in each genome. We also ran Tandem Repeat Finder (TRF v4.09) ^47^, tRNAscan-SE (v2.0.5) ^48^, RNAmmer (v1.2) ^49^, and infernal (v1.1.4) ^50,51^ to annotate other high frequency sequences and non coding elements. A final repeat annotation was created by combining the results from RepeatMasker, TRF, RNAmmer, tRNAscan-SE, and infernal, and removing redundant annotations with rtracklayer (v1.48.0) ^52^. The regions identified as repetitive were softmasked in the corresponding assemblies using bedtools (v2.30.0) ^52,53^.

Due to their large size, tandem repeats were divided into subsequences not exceeding 100 kb in length. These subsequences were then annotated using Tandem Repeat Finder v4.09 ^47^. To group these individual TR instances into biologically meaningful families, TRs were first grouped by their exact period size, and then further clustered based on consensus sequence similarity using a k-mer profiling approach (k=5). We calculated Jaccard distances between circular, strand-invariant k-mer profiles and applied average-linkage hierarchical clustering with a similarity threshold of 80%. Families spanning multiple period sizes (e.g., multimers) were merged if their representative consensus sequences demonstrated high similarity. To resolve redundant or nested TR annotations and ensure each genomic locus was uniquely assigned to a single feature, we implemented a base-level “winner-take-all” resolution strategy. TR families were prioritized based on their total genomic span, such that dominant families (those with the largest cumulative footprint) were assigned to genomic positions first, effectively masking overlapping lower-priority variants. This priority-based subtraction was extended to structural RNA elements (rRNAs, tRNAs, and snRNAs), which were given precedence over TR annotations to minimize false-positive repeat calls in RNA-rich regions.

Names were assigned to the most prevalent TR families based on their characteristic genomic distributions and elimination fate. These included Autosomal Terminal eliminated repeats (AT-1 and AT-2), Autosomal Central eliminated repeats (AC-I to AC-VI), X-Terminal eliminated repeats (XT-1 and XT-2), a minor X-Central eliminated repeat (XC), and X-Retained somatic repeats (XR-1 and XR-2). Genome-wide density was calculated as the fractional occupancy of these resolved families within 100 kb (repeat) or 10 kb (RNA) windows using the GenomicRanges package in R ^54^.

### Protein coding gene annotation

The softmasked assembly was annotated with protein coding genes via two different Braker2 (v2.1.6) ^55^ approaches and the best performing gene set was chosen per species. One approach relied on Braker2 trained with the BUSCO Nematoda odb10 proteins. The other approach relied on Braker2 trained from transcript splice sites identified from the RNA-seq reads mapped against the corresponding genome with STAR (v2.7.2b) ^56^ using the two-pass mode. As Braker2 relies on training Augustus (v3.4.0) ^57^, GeneMark-ET and GeneMark-EP and also considers predictions from GeneMark-ES (GeneMark Suite v4 May, 2021) ^58^, we included each of these predicted gene sets in our assessment. The longest protein per gene was extracted using AGAT (v0.8.0; https://github.com/NBISweden/AGAT) and the gene set completeness and redundancy was assessed with BUSCO (v5.2.2) ^24^. We annotated the protein domains in the longest protein predicted of each gene with interproscan (v5.39-77) ^59,60^. The Braker2 final annotation based on protein homology yielded the most complete gene set (Table S7). This gene set was functionally annotated with interproscan, as in the previous chapter, and also using the eggnog-mapper web service ^61^.

### Expression analysis

Gene expression was assessed using previously published RNA-seq data ^23^ derived from L2 stage nematodes fated to become hermaphrodites or females, L2 nematodes switched from hermaphrodite fate to female using dafachronic acid, adult males and a population of mixed sex and stage nematodes (Table S8). RNA-seq reads were mapped to the genome with STAR (v2.7.2b) ^56^ using the two-pass mode.

### Identification of PDE boundaries

We identified the boundaries of the germline-restricted sequences by searching for all the supported chromosome ends in our assemblies. We identified the chromosome ends by extracting all the telomeric reads from the PacBio sequencing read sets and then mapping. This is an inverse version of the approach described in the “Genome assembly and curation” section. We oriented these reads so that the telomeric sequence was found at the start and we removed the telomere repeat sequence. We then mapped these trimmed telomeric reads against the assembly with minimap2 (v2.24) ^62^ using the `-ax map-hif`i parameter and collected the start position if the read mapped in the forward strand or the end position of the alignment if it mapped on the reverse. Only alignments with mapping quality (MAPQ) higher than 30 were considered. We summarised in R the number and orientation of these coordinates. Finally, we manually curated these coordinates by exploring the whole read set alignments in GAP5 ^63^ at each of the candidate regions. Additionally, we validated the same positions with BleTIES, a software toolkit for detecting DNA elimination in ciliates with long reads ^v0.1.11 64 64^.

### Motif identification and search

We extracted 100 bp centred at each of the PDE boundaries, taking only one window where the eliminated DNA spanned less than 10 bps. For identification of and search for the motifs we used tools from the MEME suite (v5.1.1) ^65^. Specifically, we identified the enriched motifs using MEME with parameters `-nmotifs 6 -evt 0.0001 -maxw 90` ^66^. We used FIMO with the `--thresh 1e-5 --max-strand` parameter to identify all the regions across the genome with similarity to the corresponding species-identified motifs ^67^. To align the position specific weight matrices corresponding to the main motif, we used the Bioconductor package motifStack (v1.40.0) ^68^. We visualised as sequence logos the aligned motifs using ggseqlogo (v0.1) ^69^.

### Orthology inference

To identify genes with homology to genes in *C. elegans* and *A. suum*, we downloaded the annotation files and assemblies of these species from Wormbase Parasite ^70^ and used AGAT (v0.8.0) to obtain the longest isoform per gene ^45^. We similarly obtained the longest isoform prediction for each of the five *Oscheius* species. We then ran OrthoFinder ^71^ with these proteomes using Diamond BLASTp ^72^ as the sequence search program, and multiple sequence alignment with MAFFT ^73^ to infer the gene trees with IQ-TREE ^74^. For the Markov clustering step we kept the default inflation value of 1.5. For data comparing presence/absence and identifying coordinate overlaps with the eliminated regions, we used RStudio, the tidyverse (https://www.tidyverse.org/) and GenomicRanges ^54^ packages of R Bioconductor (https://www.bioconductor.org/).

### Immunostaining and FISH

Cytological experiments were performed on embryos laid by *A. rhodense* adult hermaphrodites 40-48 hours post-dauer. For embryo collection, 20-40 gravid adults were placed onto unseeded NGM plates and allowed to lay for one hour. Embryos and adults were then collected by washing with M9 containing 0.1% tween. Embryos and adult worms were allowed to settle onto charged slides (Fisher Scientific, Cat# 1255015), then adult worms were cut open to extrude in-utero embryos, and excess liquid was removed before proceeding to fixation.

Embryos were fixed in 1% PFA in egg buffer with 0.1% tween for 5 min at room temperature, immersion in liquid N_2_, and transfer to -20 °C methanol for 5 min ^75^. For immunostaining (adapted from ^75^) fixed slides were transferred to PBST (3x 5 min washes) and blocked for 30 min in 0.7% BSA in PBST. Slides were incubated overnight with primary antibodies diluted in block; the following primary antibodies were used at the indicated dilutions: anti-α-Tubulin-FITC (clone DM1A, Thermo Fisher Scientific cat#F-2168, RRID: AB_476967, 1:1000), anti-H3K9me3 (Abcam Cat# ab8898, RRID:AB_306848, 1:500). After 3x 10 min washes in PBST, slides were incubated with secondary antibody (goat-anti-rabbit 555; Thermo Fisher Scientific Cat# A-21428, RRID:AB_2535849, 1:200) and DAPI (5 µg/ml) for 2.5-3 hours at RT, washed in PBST (3x 5 min) and mounted with Vectashield (Vector Laboratories, Cat#H-1000) using No. 1.5 coverslips (Fisher Scientific, Cat# 12541019). For FISH (adapted from ^76^): fixed slides were transferred to 2x SCCT (3x 10 min washes) followed by preincubation in 50% formamide in 2x SCCT at 37 °C for 1-2 hours. Then, 1.5 µl each of 10 µM direct-labelled ssDNA oligo probes (IDT, sequences provided below) were added to 40 µl hybridization solution (50% formamide, 10% dextran sulfate in 2x SCCT) and applied to samples; slides were then denatured on a heat block at 91 °C for 2 minutes and hybridized overnight at 37 °C. Then, slides were washed in 2x SCCT (3x 5 min) and PBST (2x 5 min) before incubating with DAPI in PBST for 30 min at RT. After washing in PBST (3x 5 min), slides were mounted in Vectashield using No. 1.5 coverslips.

FISH Probe sequences were:

AT-1_Cy3 (A-elim): ACATACCAAGTGCACATCACGTGTACAT/3Cy3Sp/

XR-1_Cy5 (X-retain): ACTGTCATAGAATGTTCATATAGGTGT/3Cy5Sp/

XT-2_488 (X-elim): ATGTAACACGCTGAAATTCGTTGCATT/3ATTO488N/

XT-1_Cy3: ATGCCTTTAACACCATATACTCACCGAT/3Cy3S

All images in **Figure 4** were acquired on a DeltaVision OMX Blaze microscope with a 100 x 1.4 NA widefield objective and z-stacks spaced at 200nm. Images were then deconvolved and registration corrected with SoftWoRx and, where necessary, stitched together using the ‘Grid/Collection stitching’ FIJI plugin ^77^. All displayed images are maximum intensity projections through the entire embryo.

## Supplementary Materials

**Table S1.**
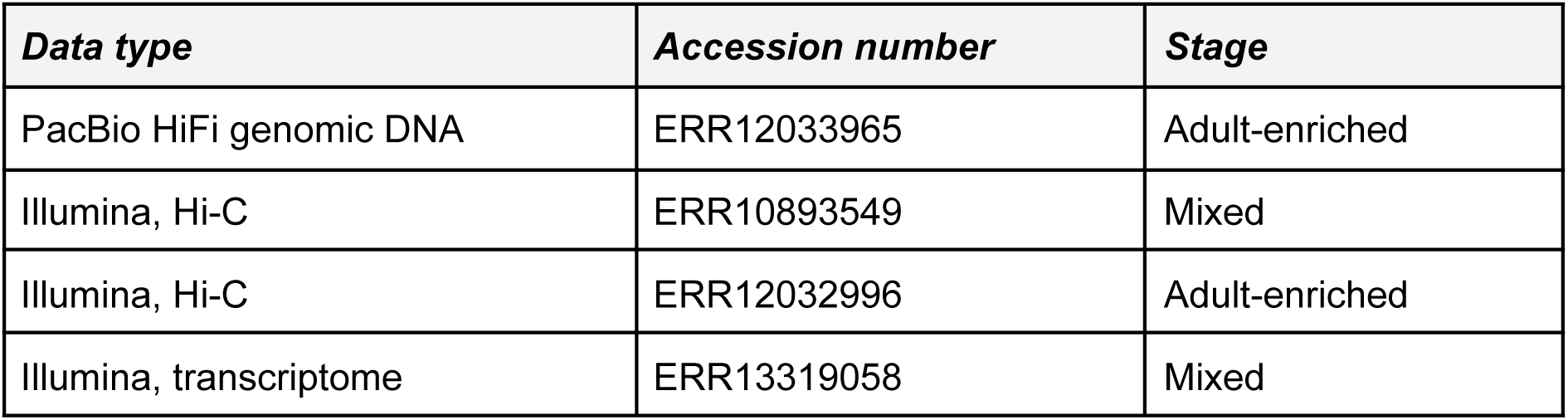
Accession numbers for genomic and transcriptomic data from *Auanema rhodense*.

**Table S2.**
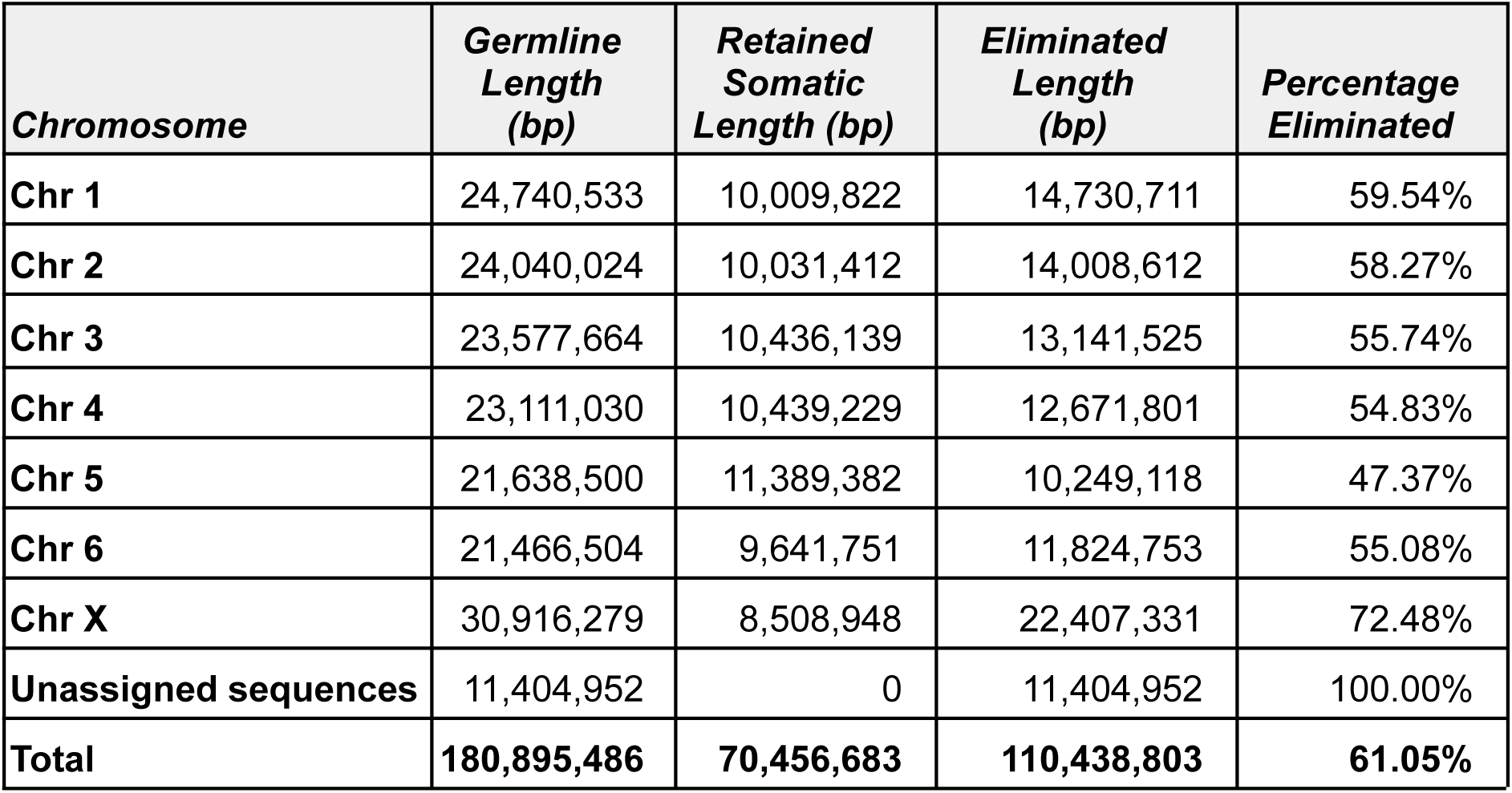
Quantification of DNA eliminated per chromosome during programmed DNA elimination in *Auanema rhodense*.

**Table S3.**
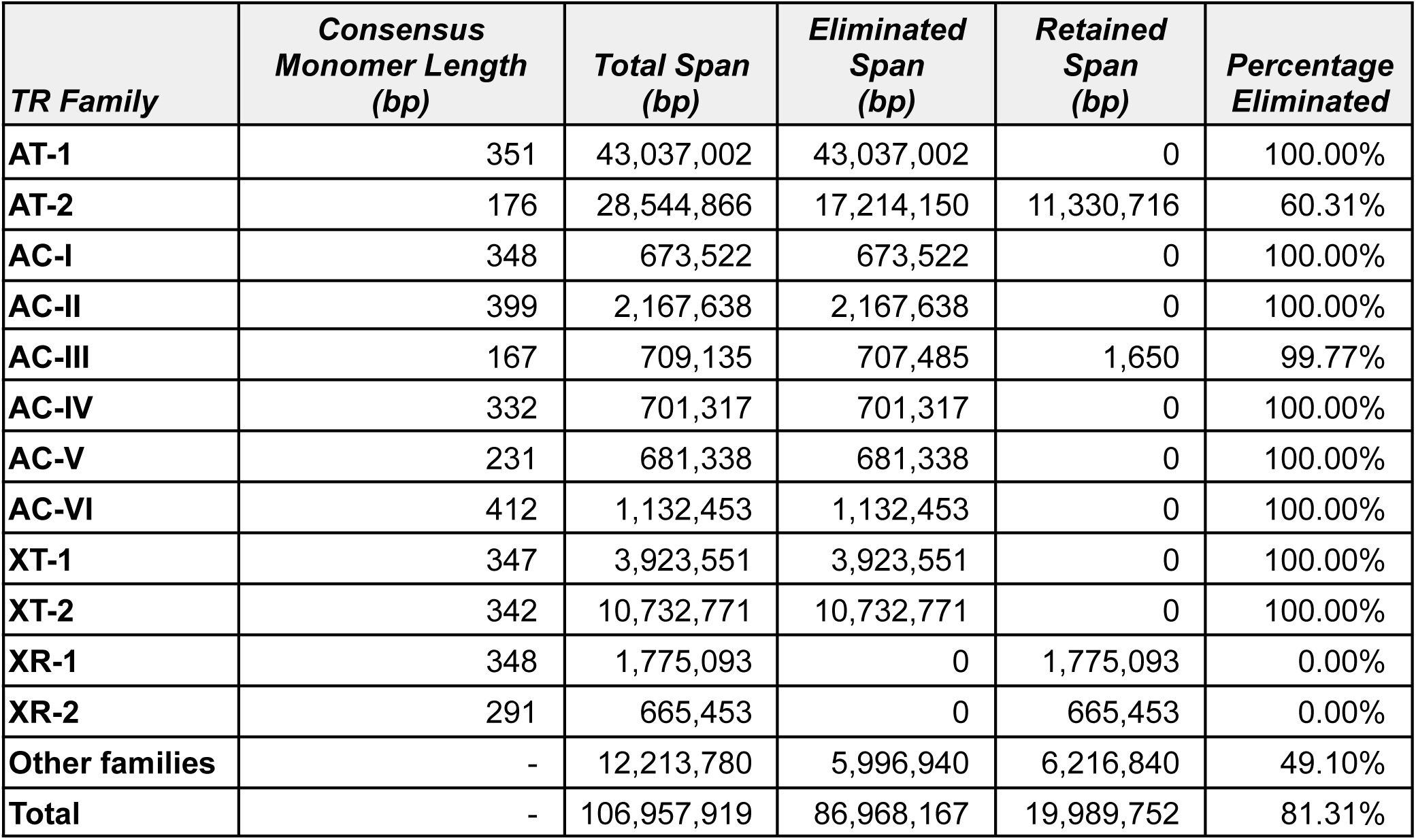
Tandem repeat families in *Auanema rhodense*.

**Table S4.**
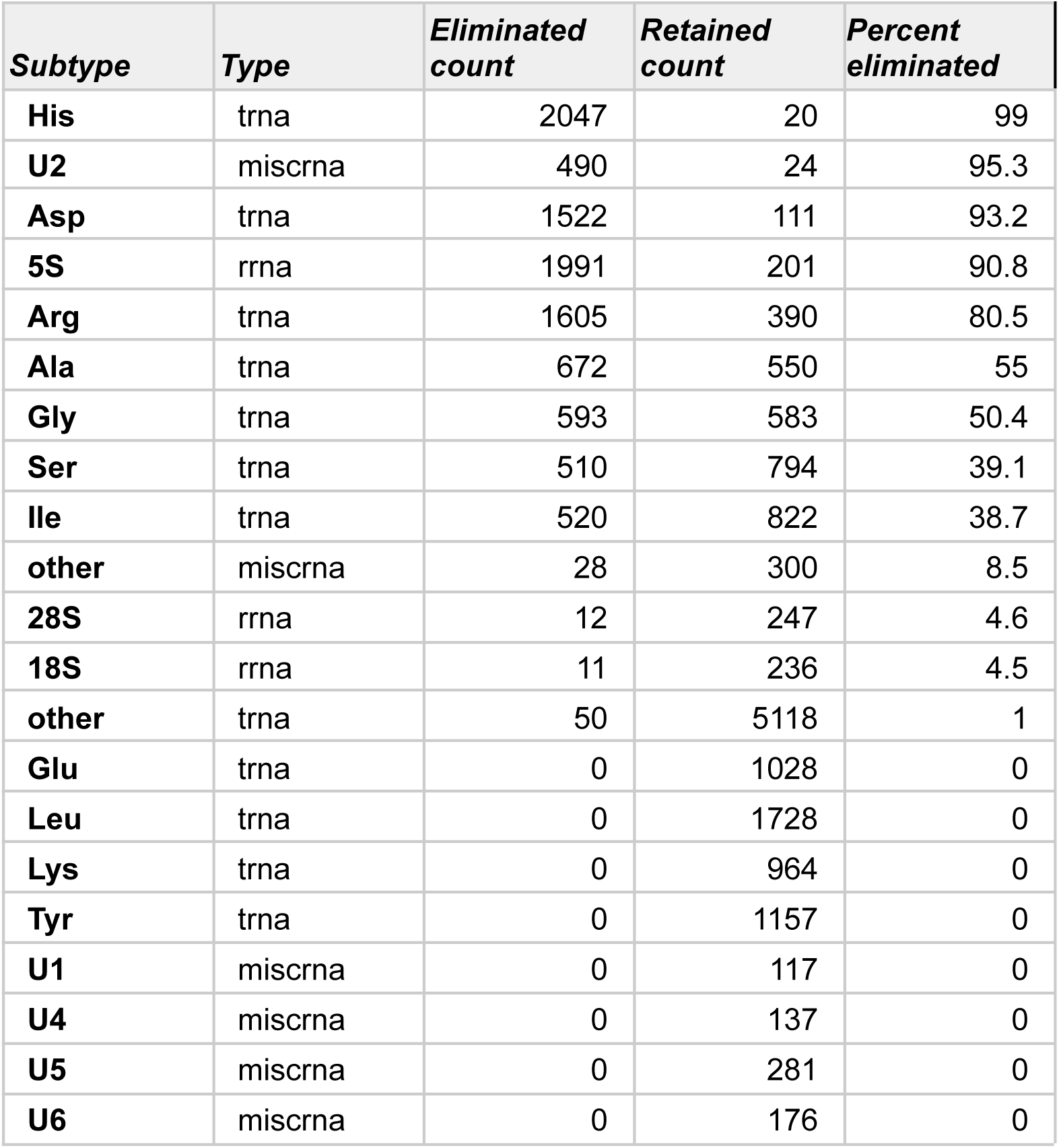
ncRNA genes present in eliminated DNA in *Auanema rhodense*.

**Table S5.**
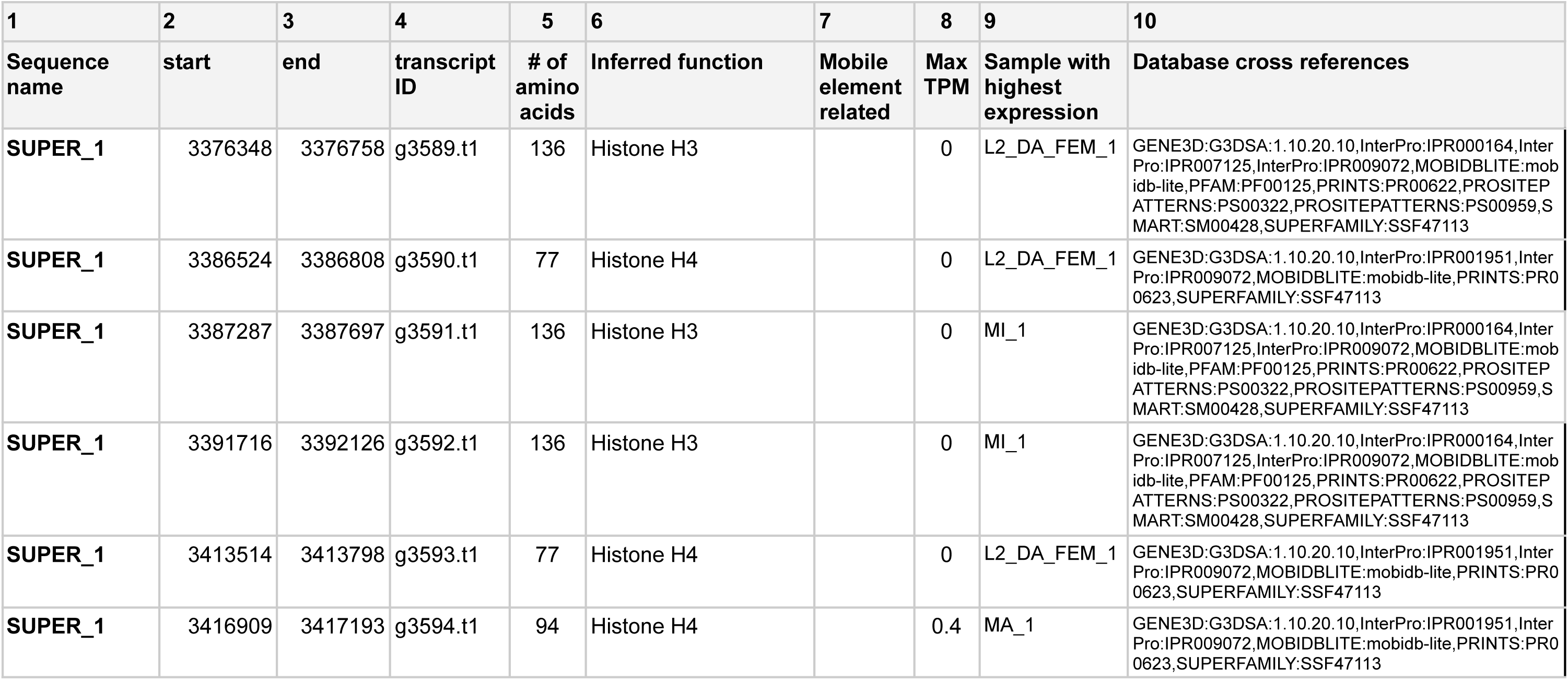
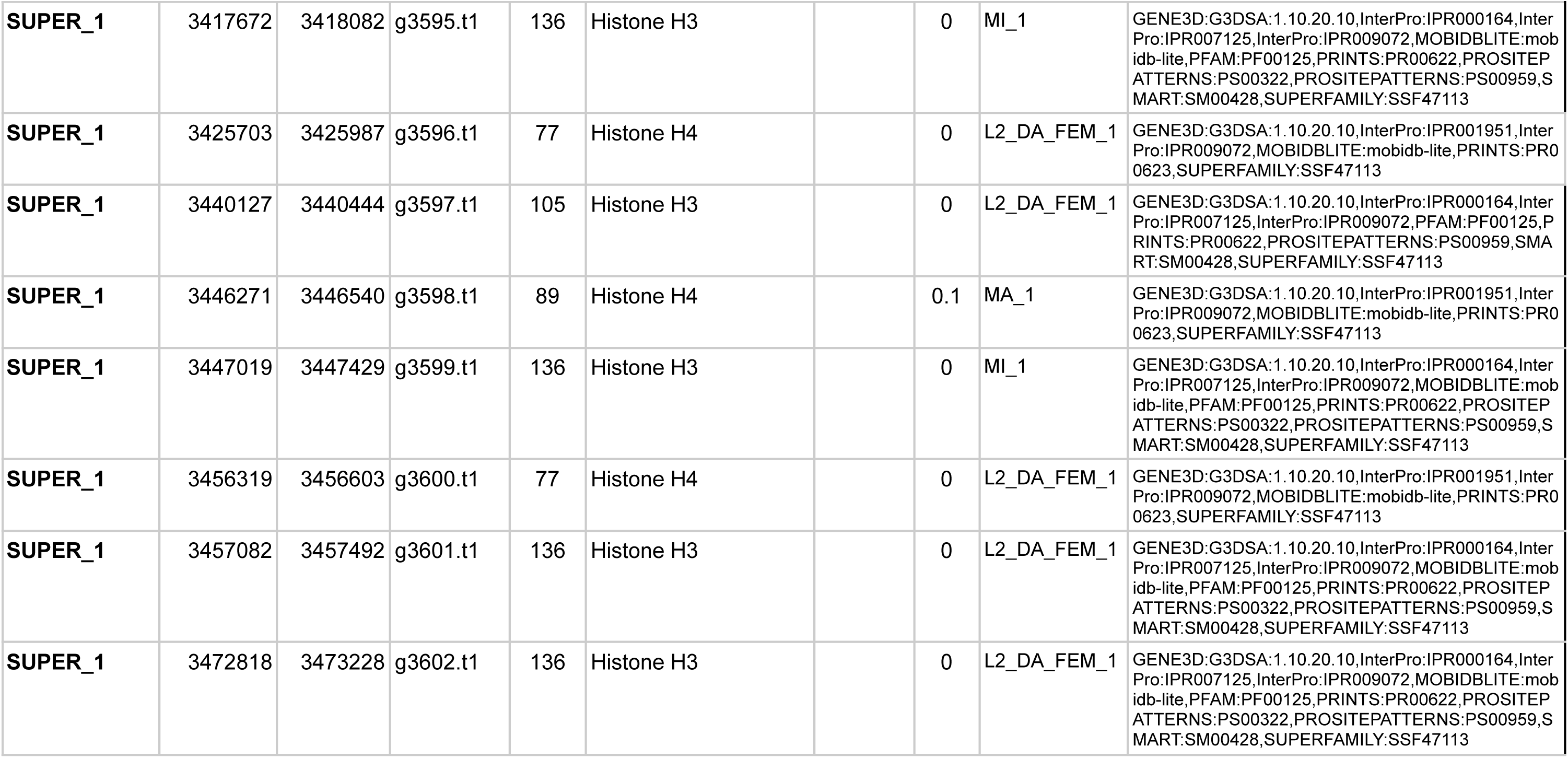
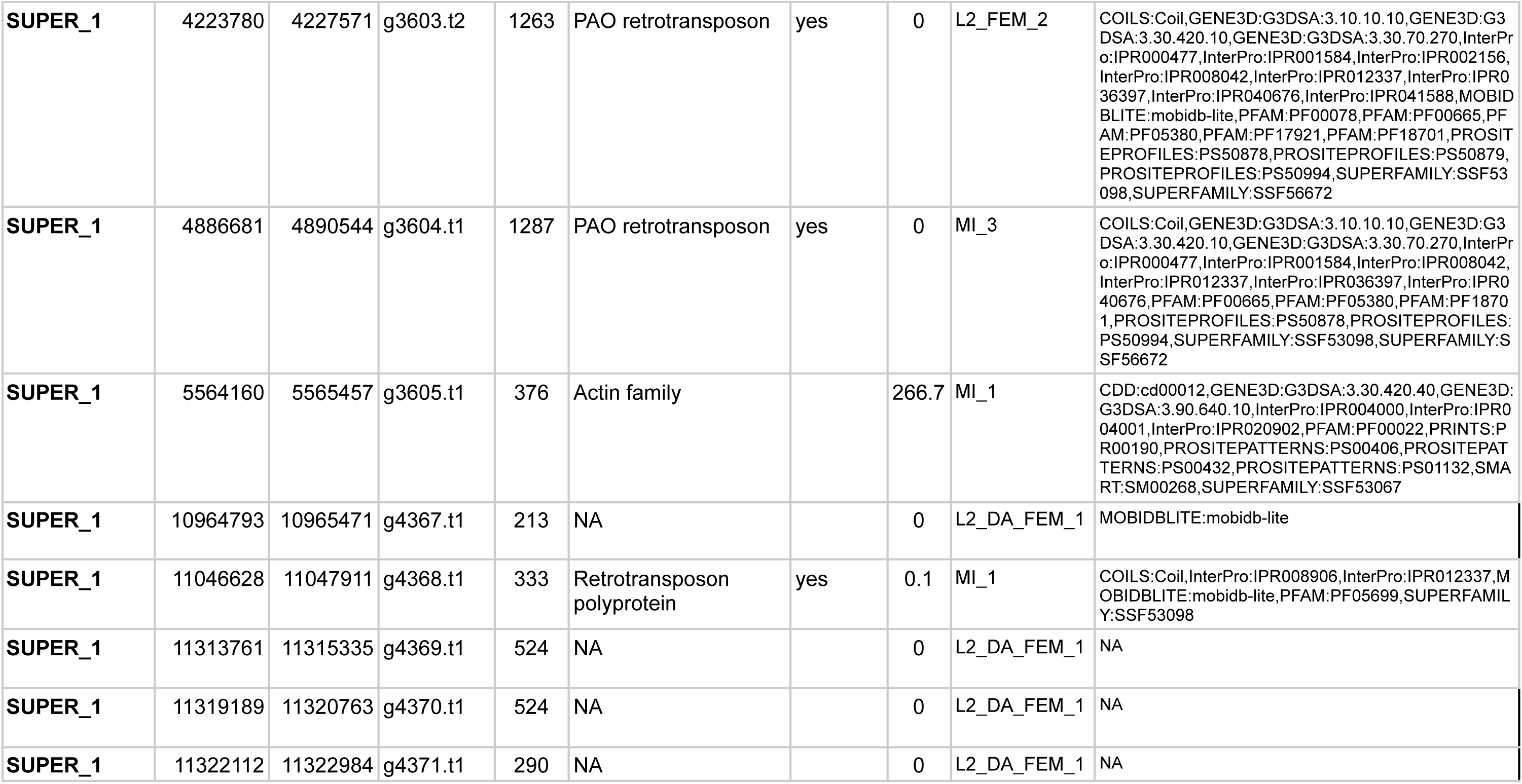
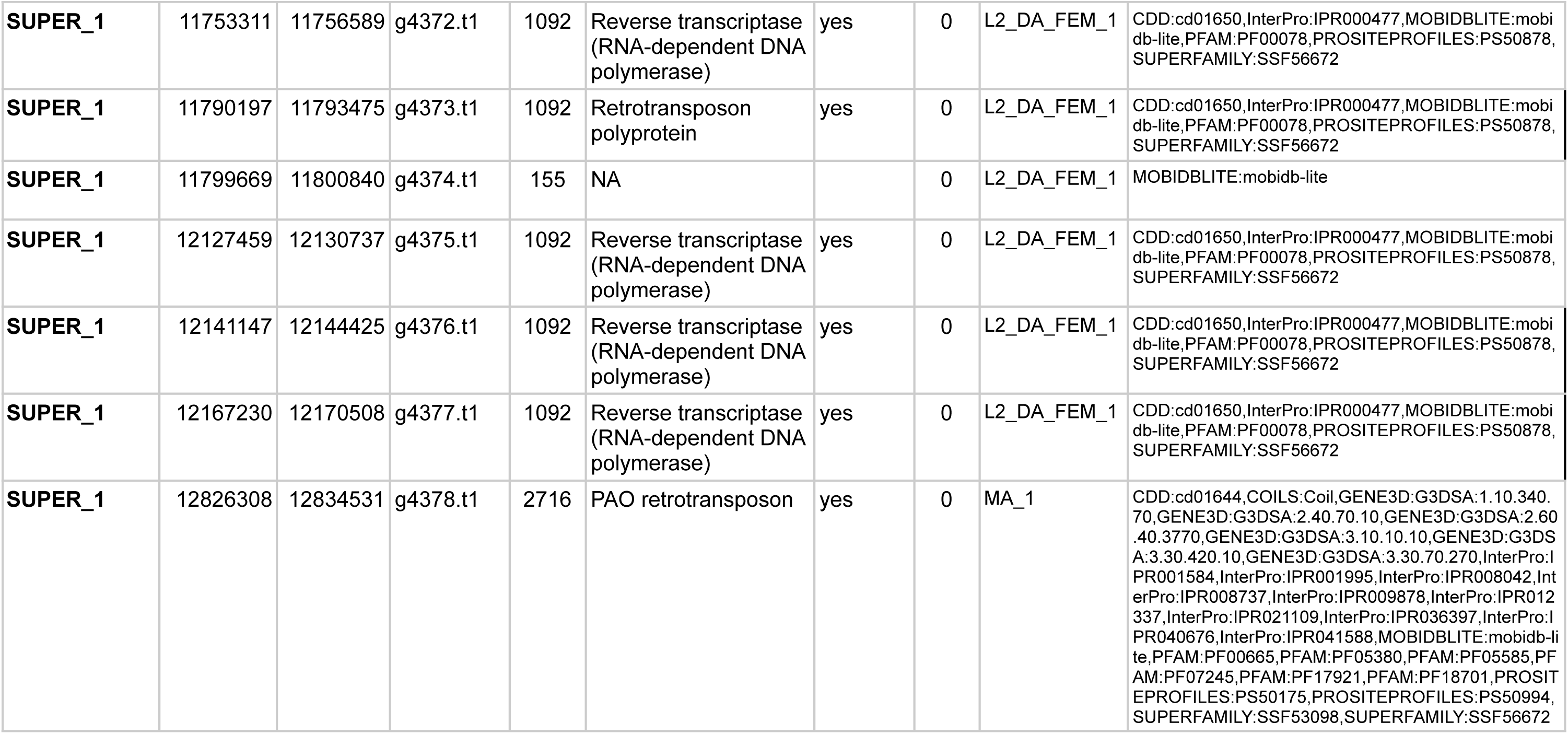
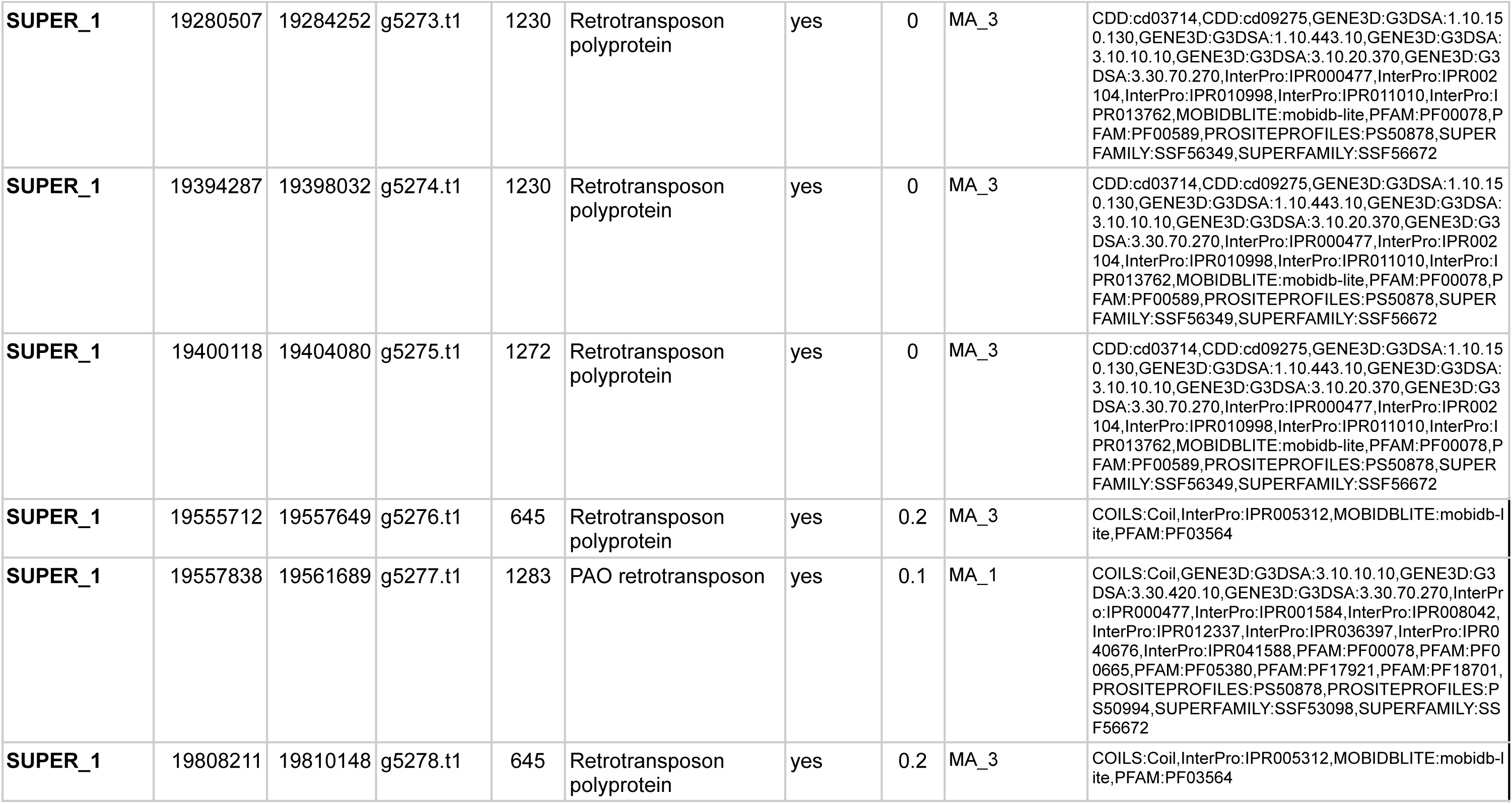
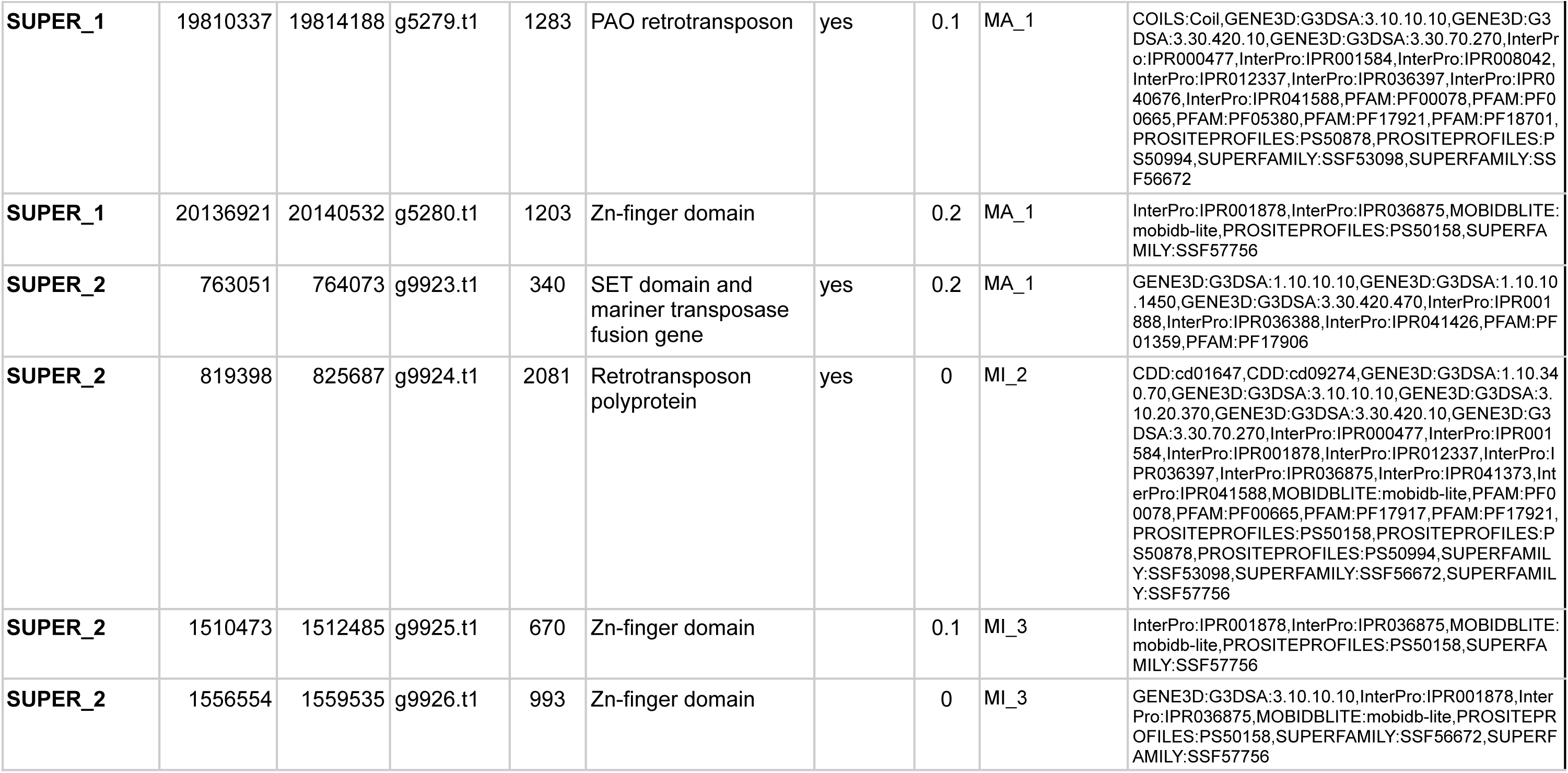
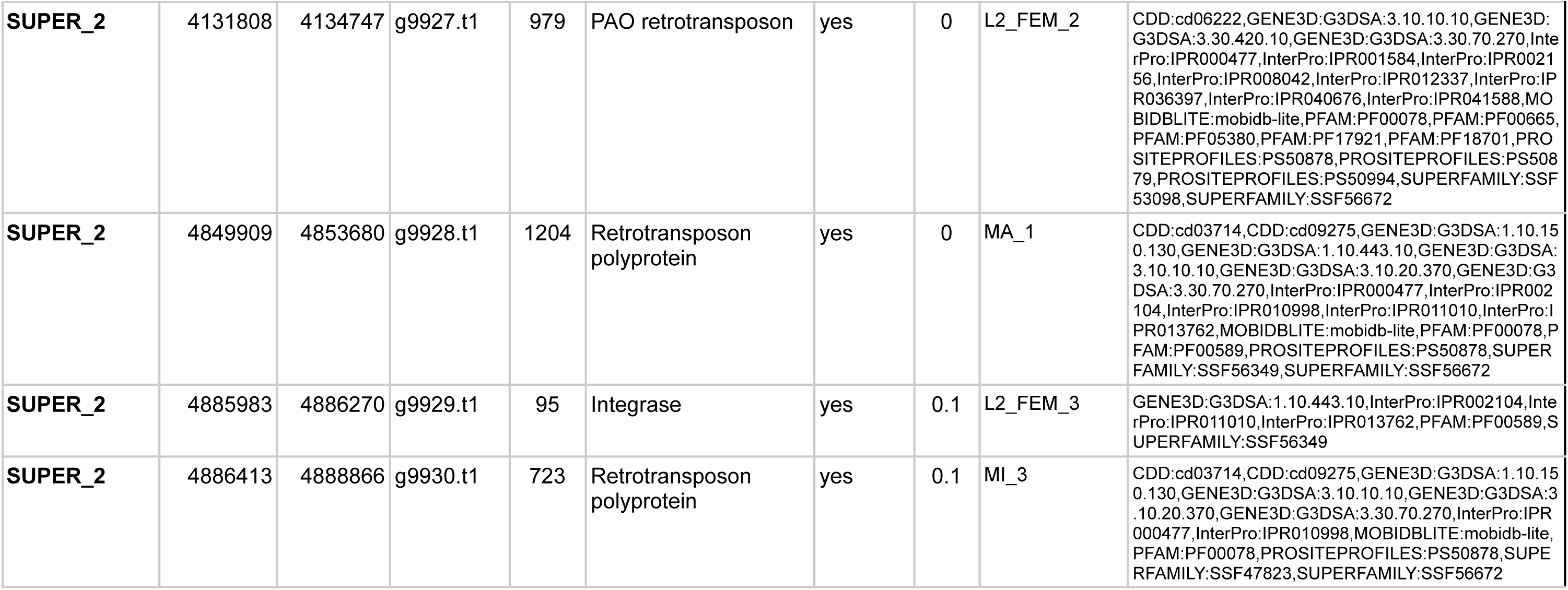
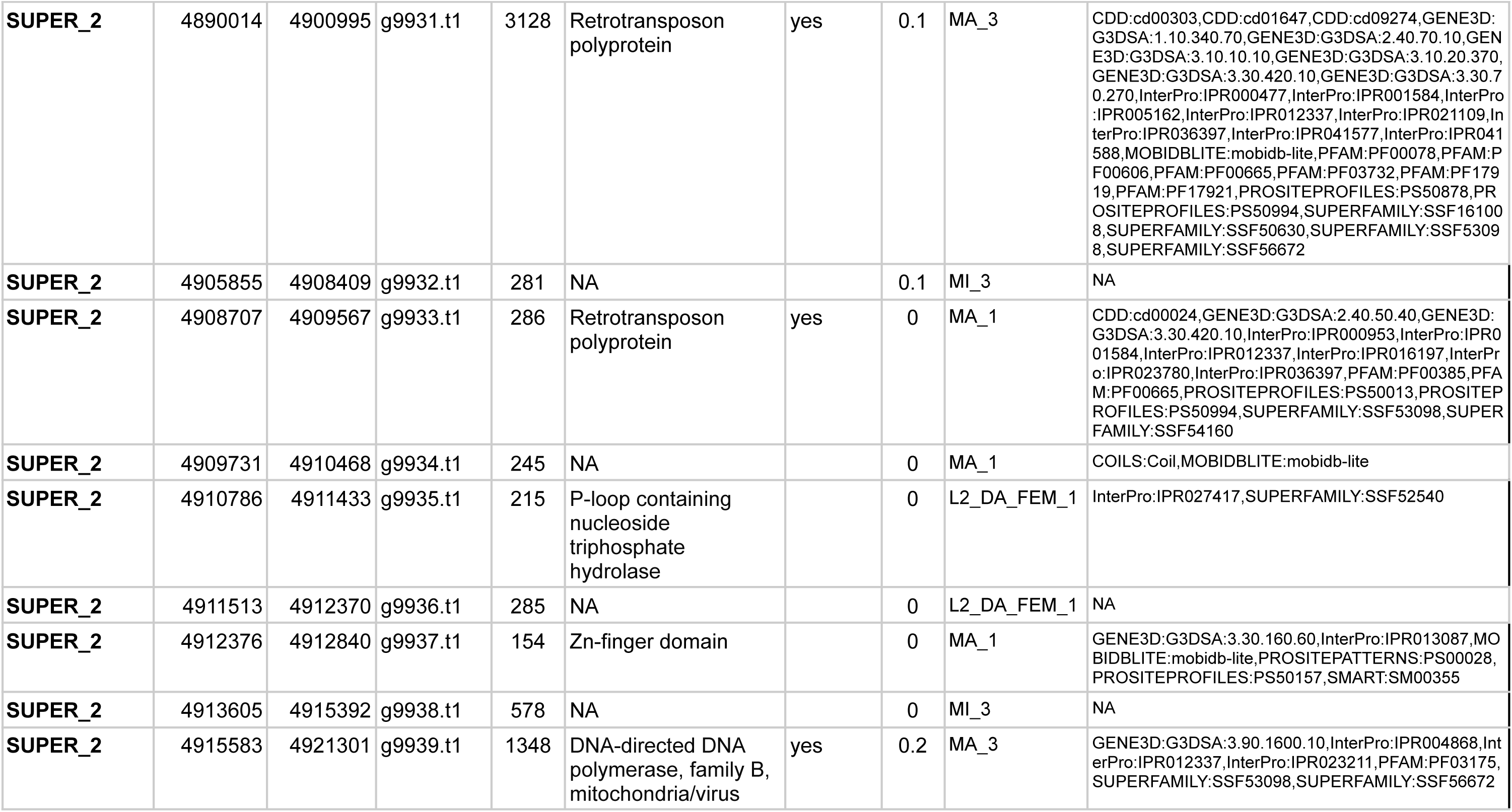
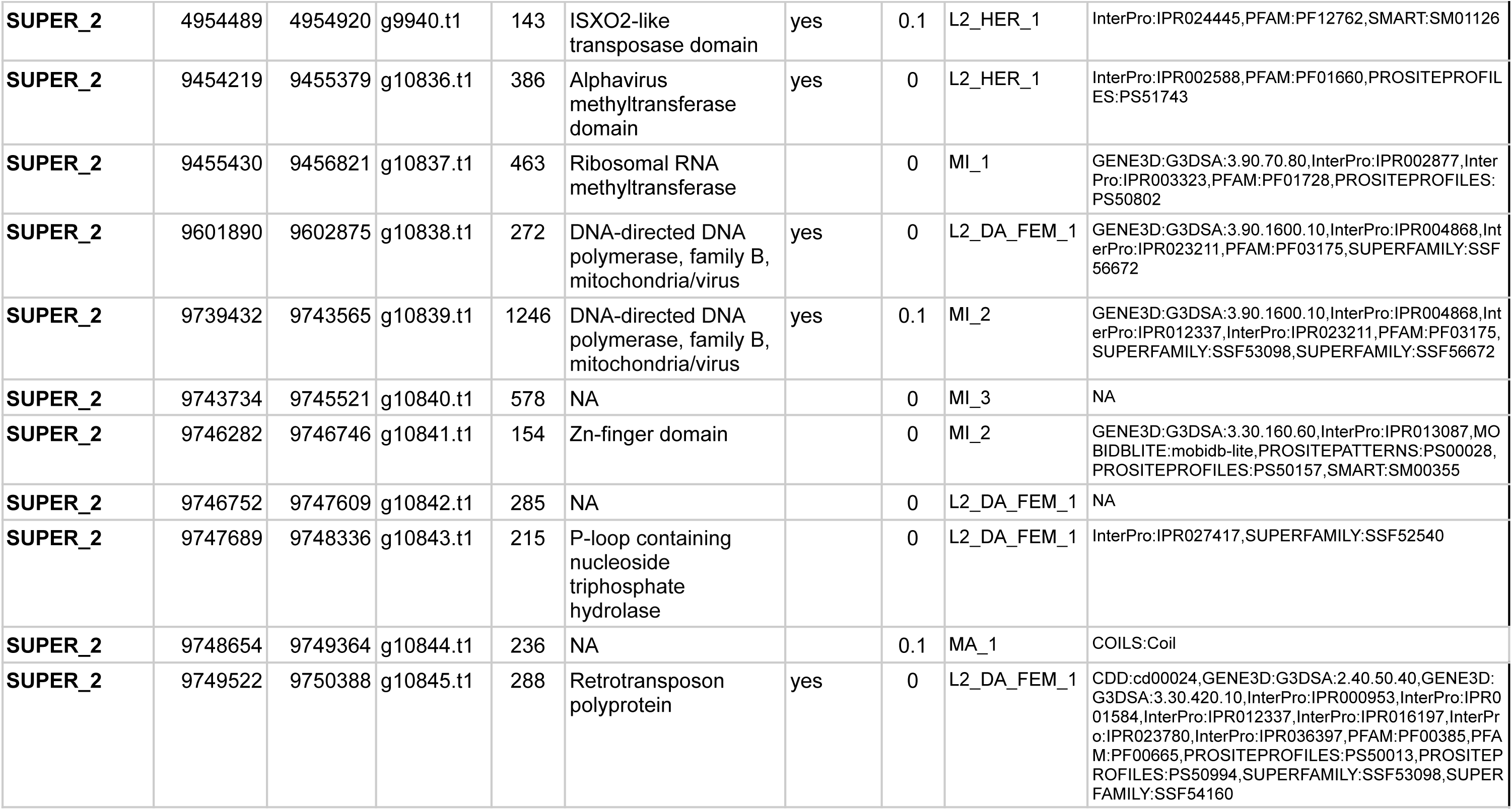
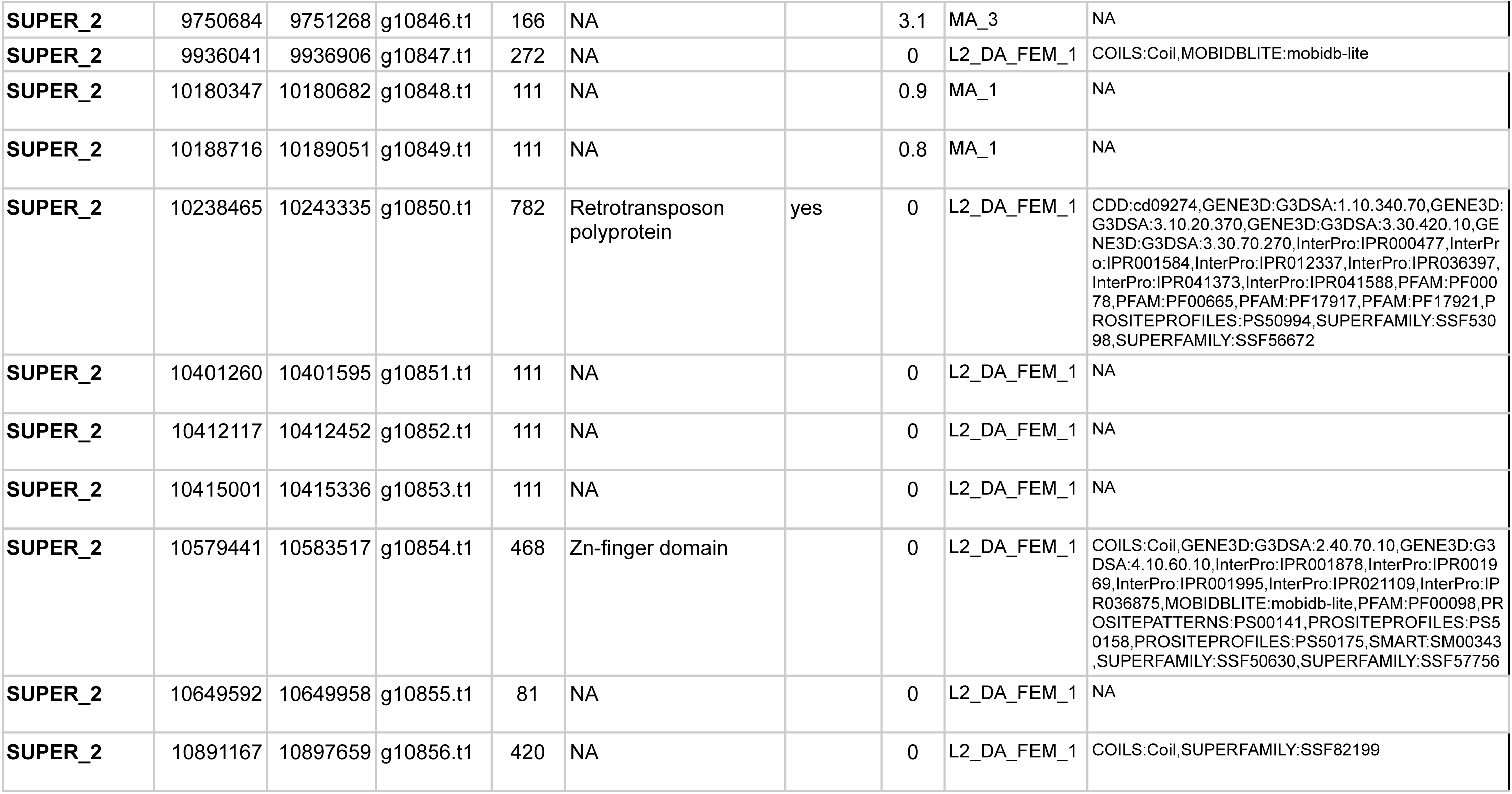
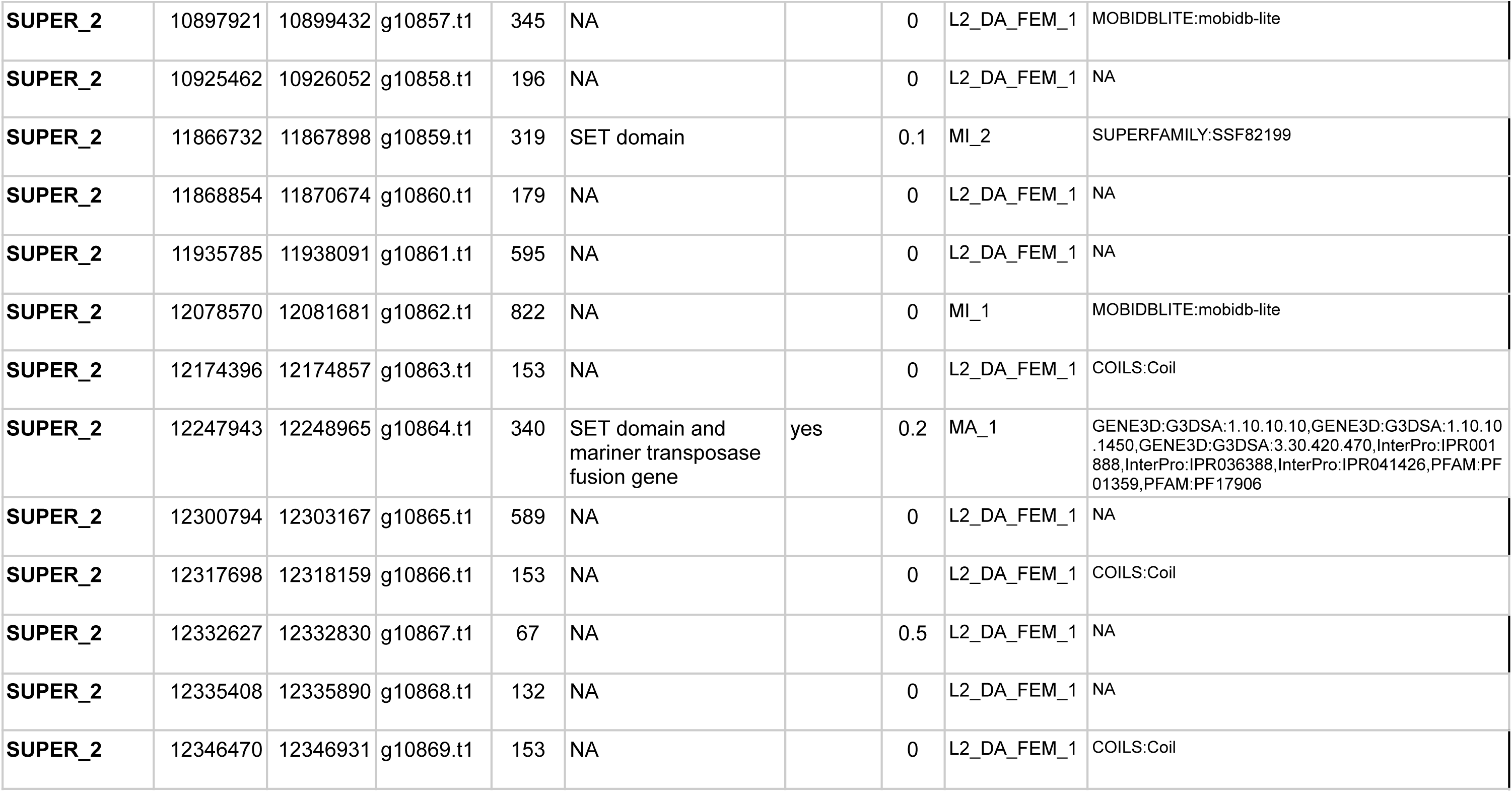
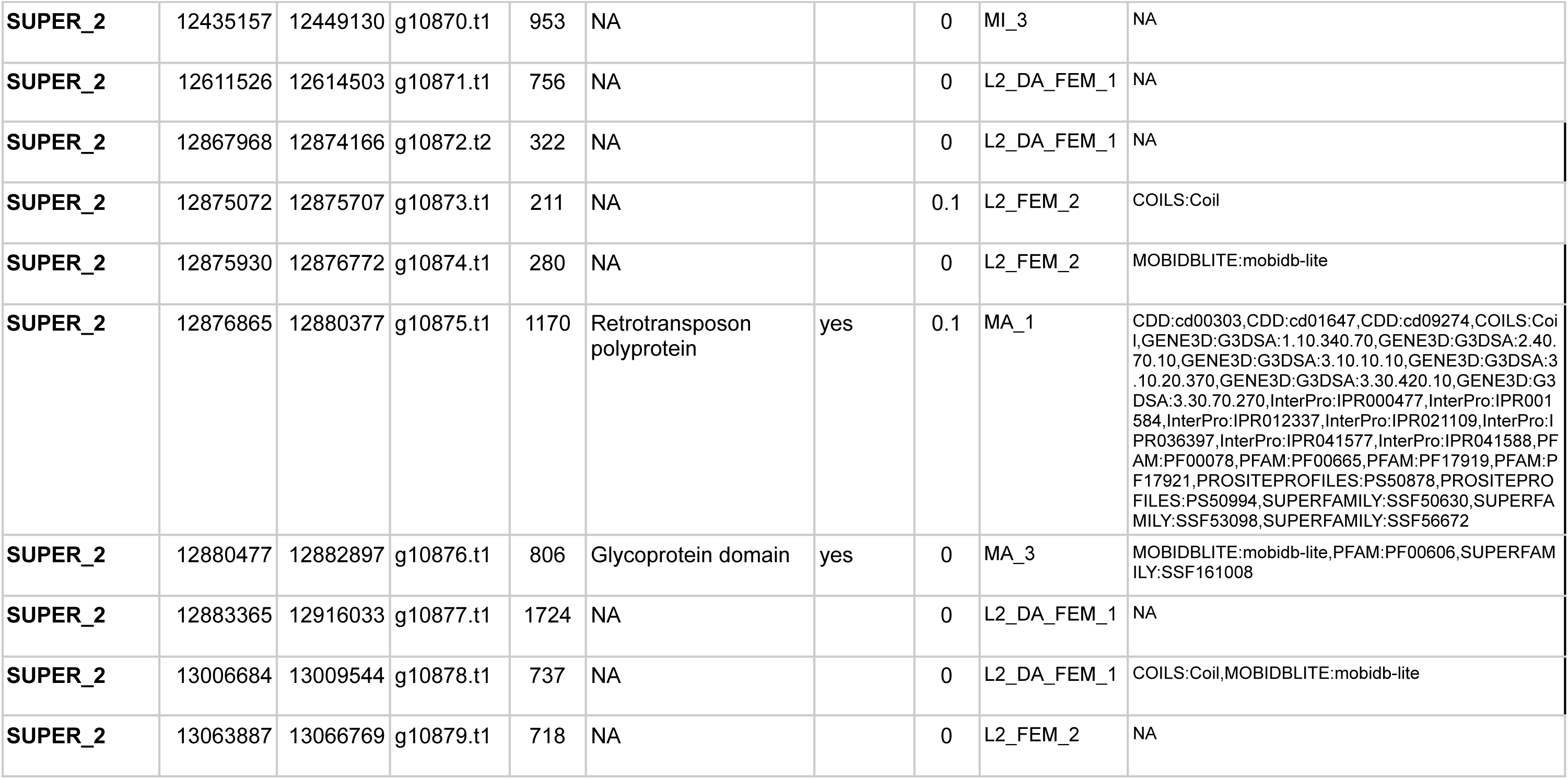
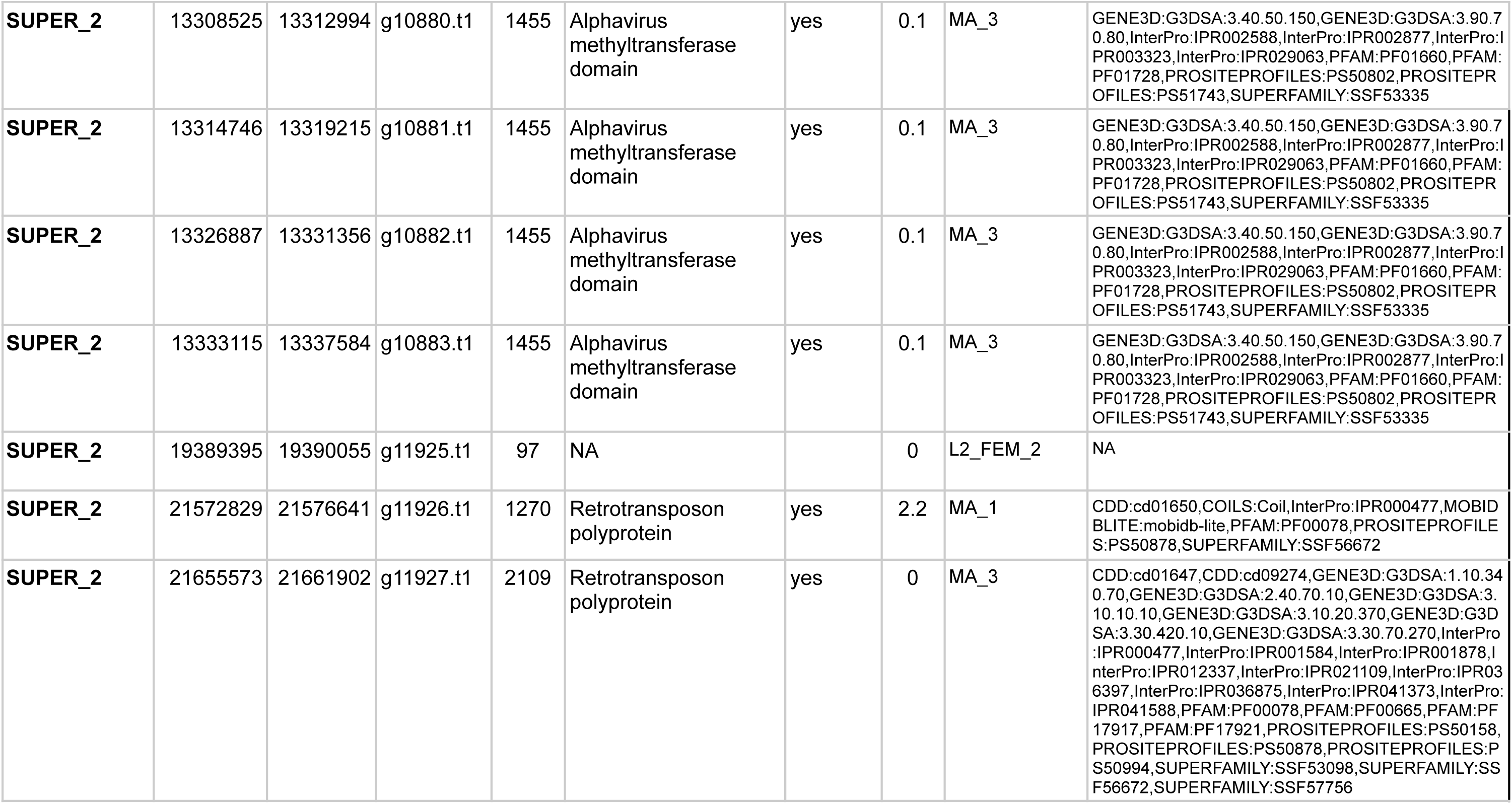
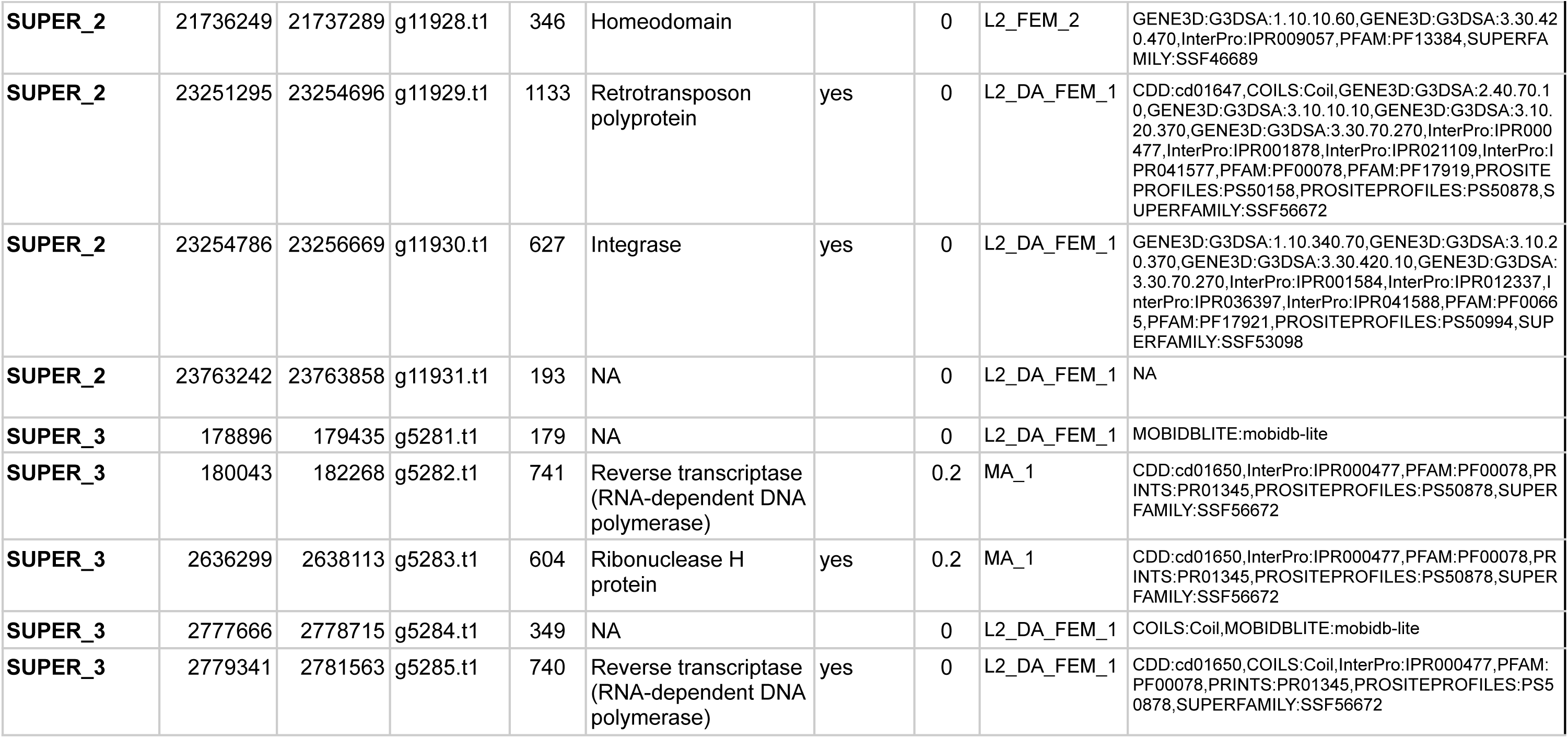
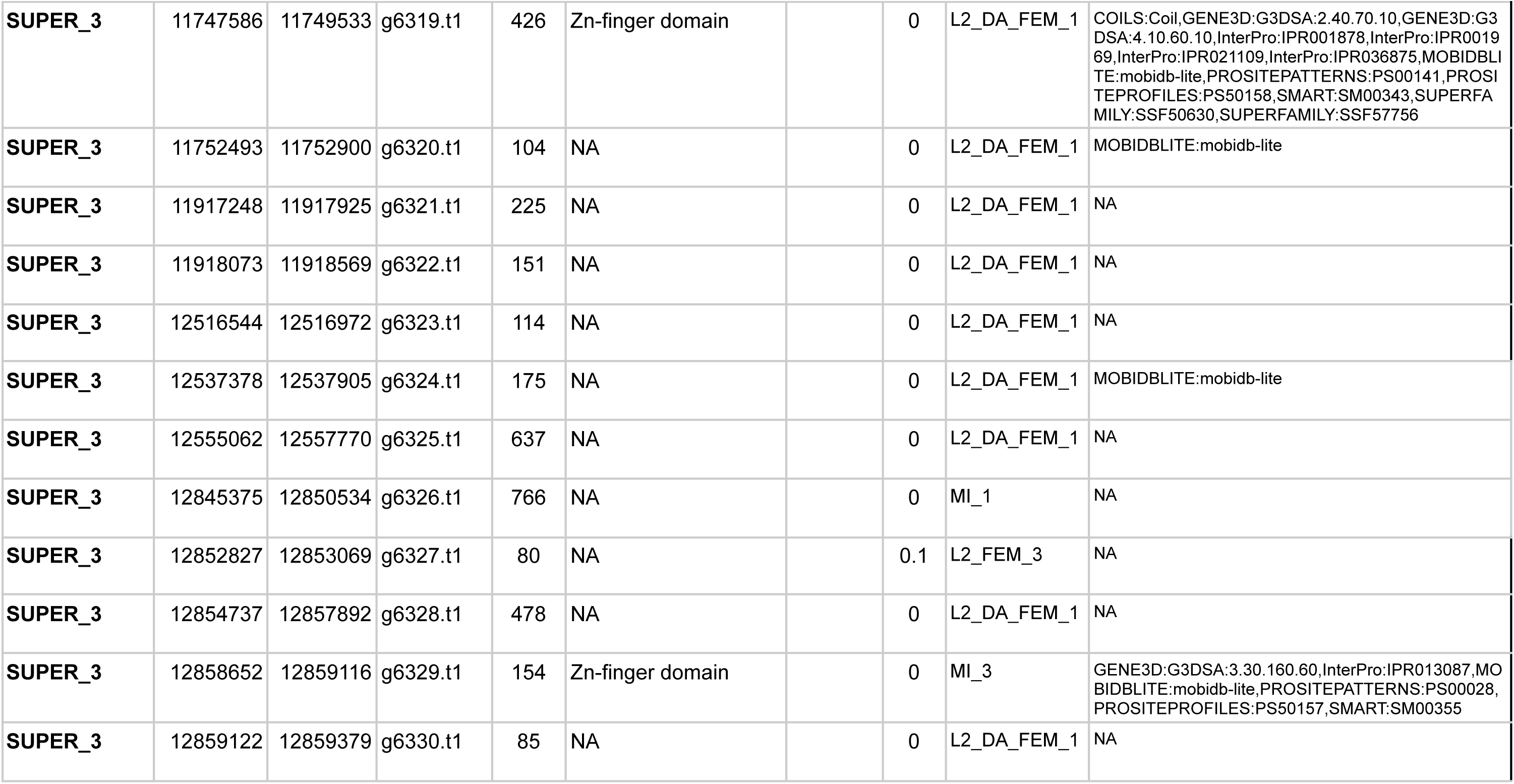
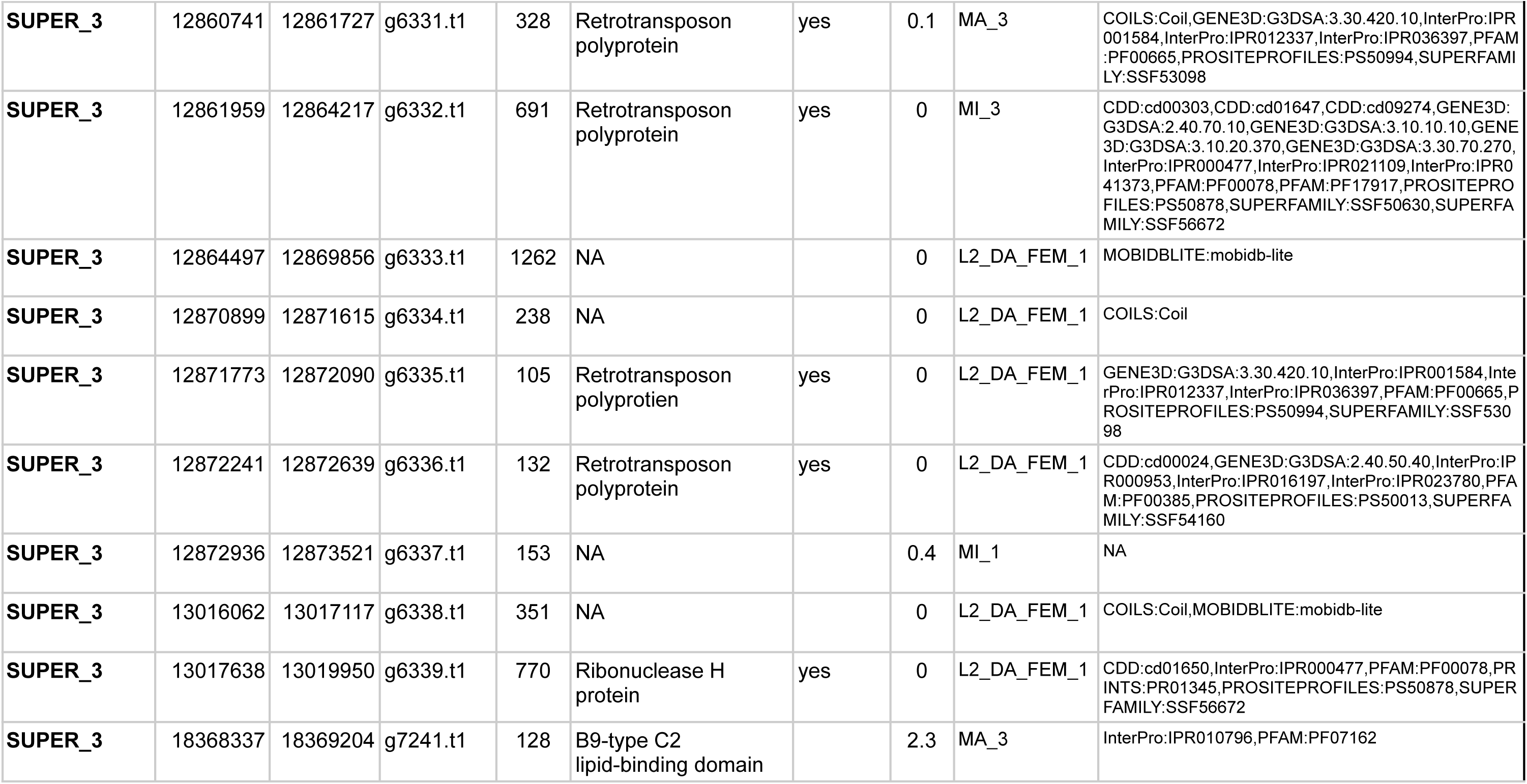
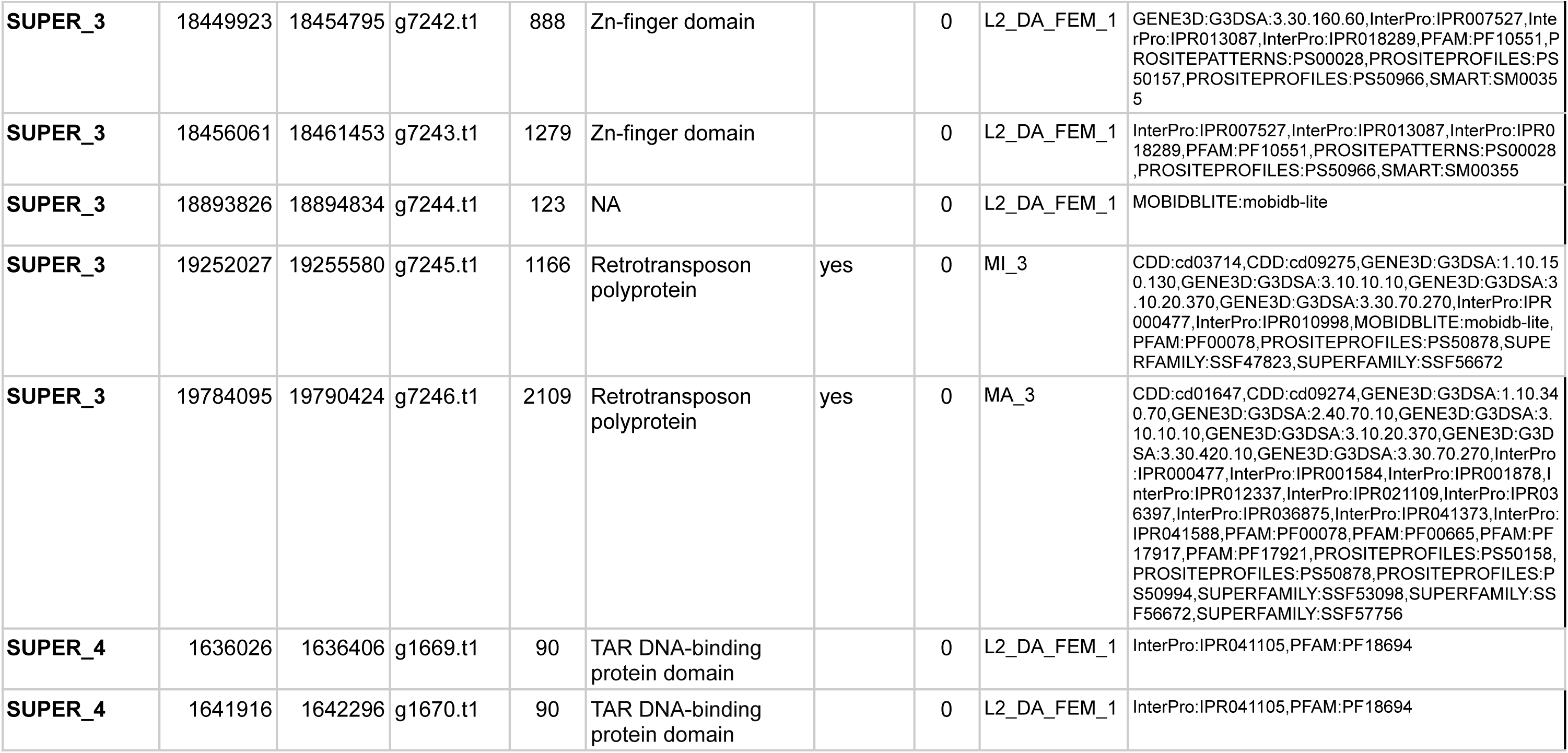
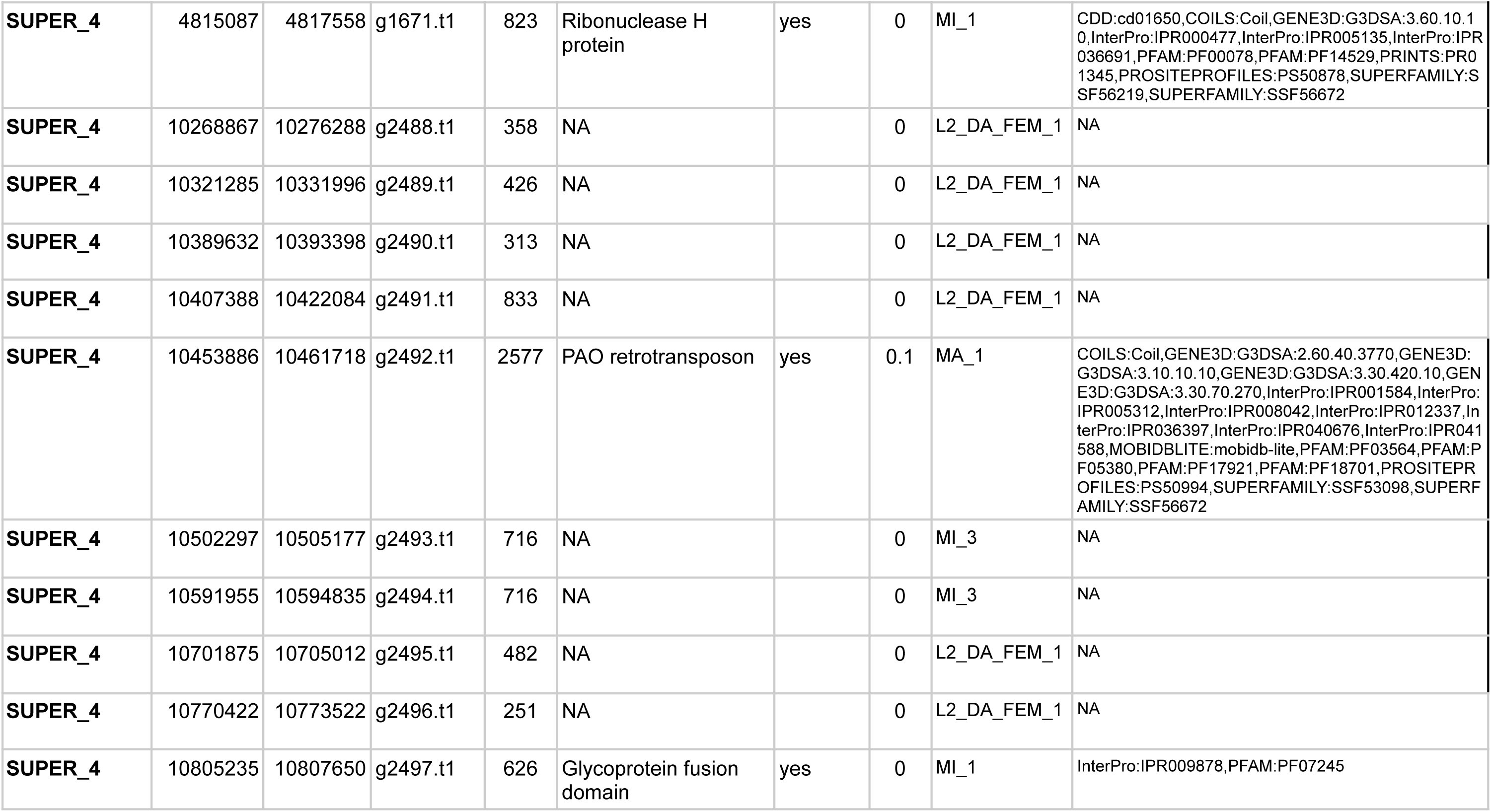
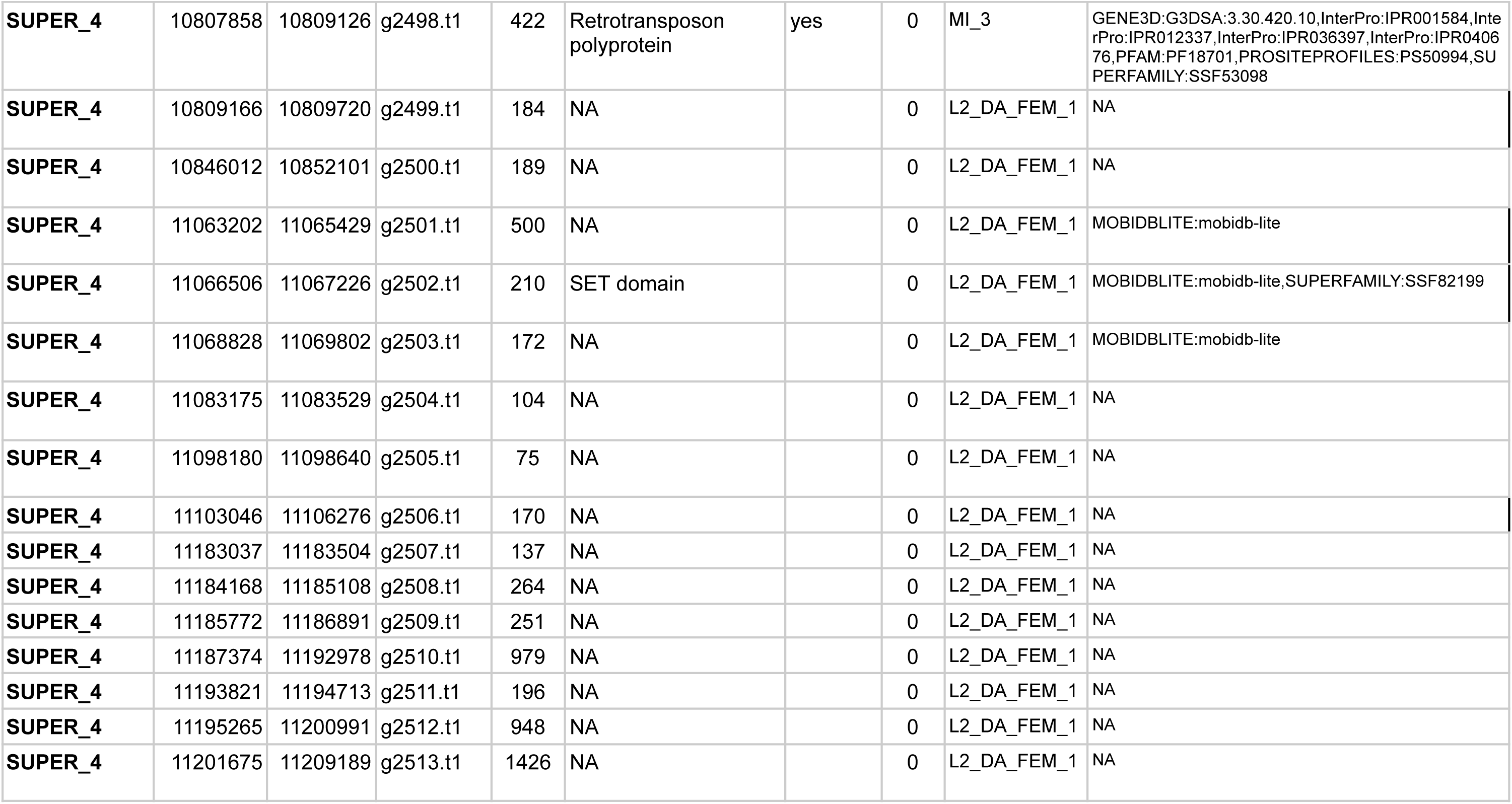
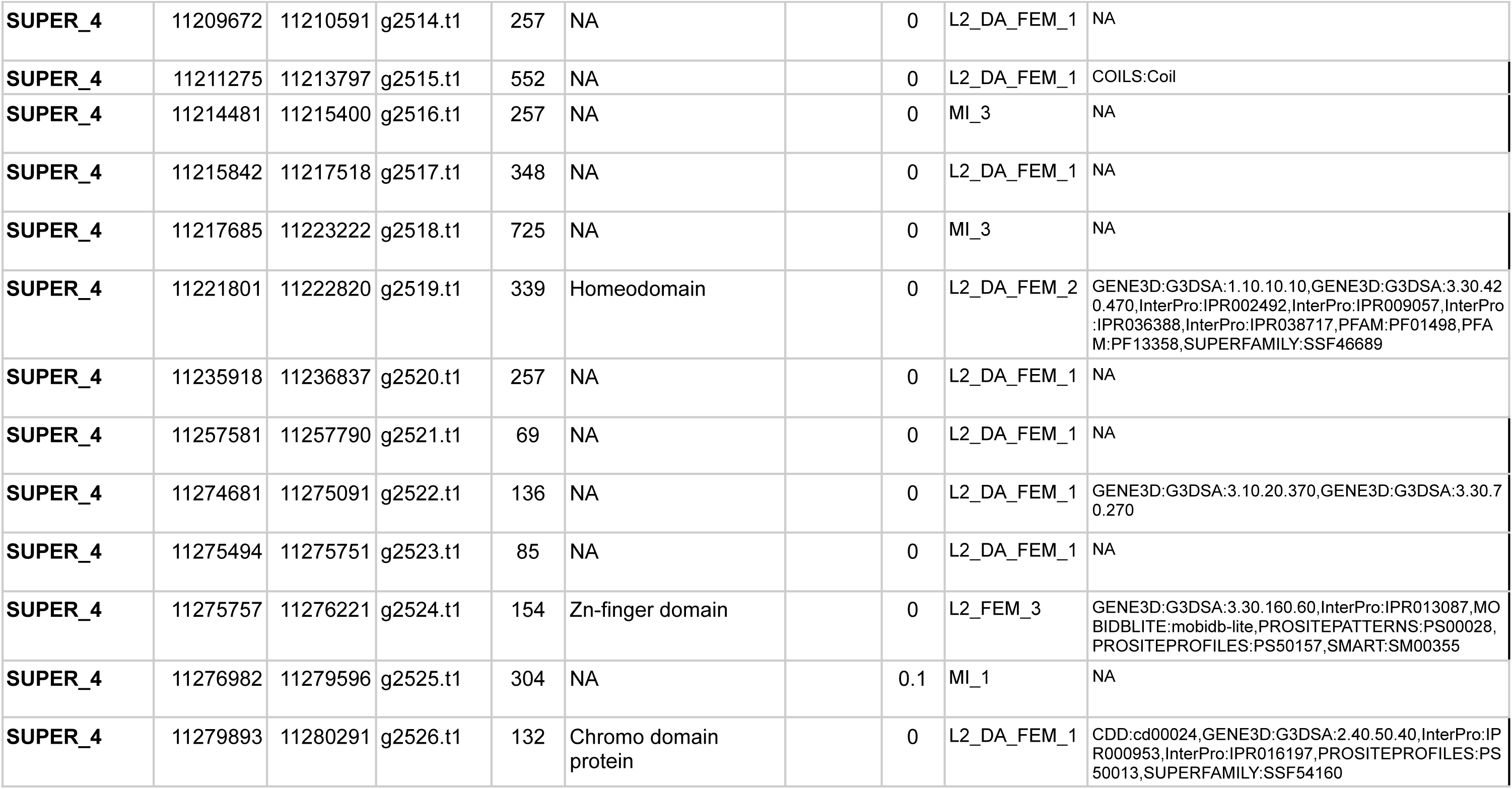
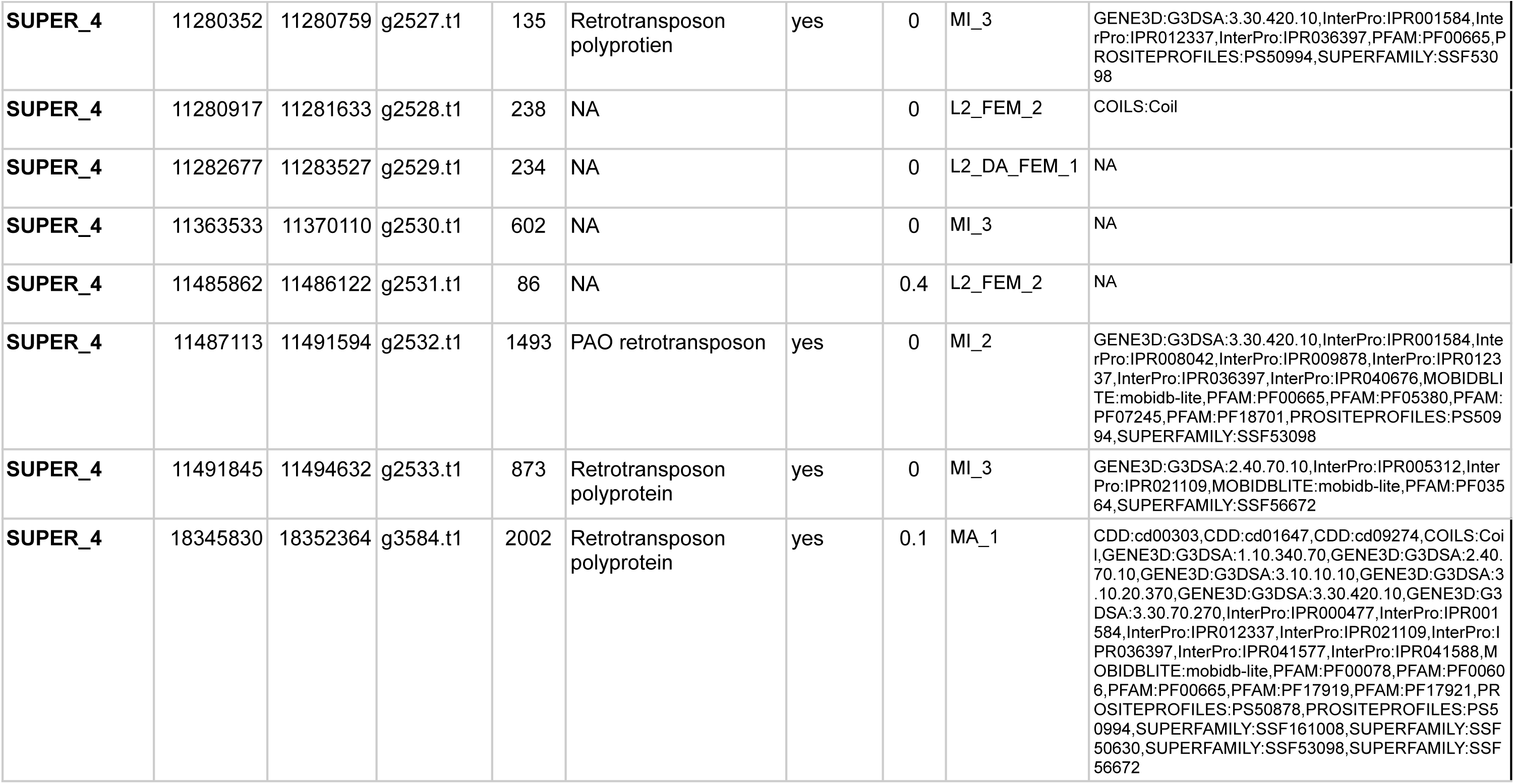
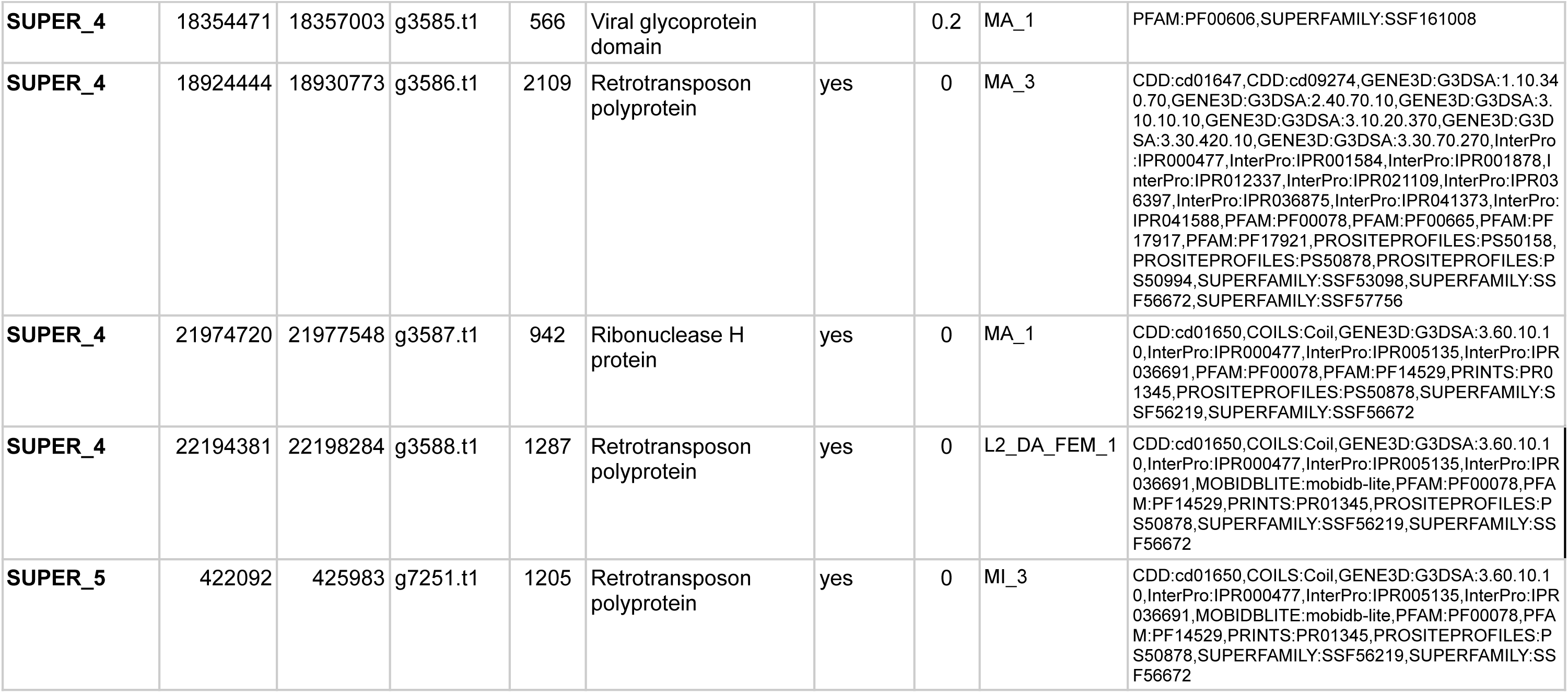
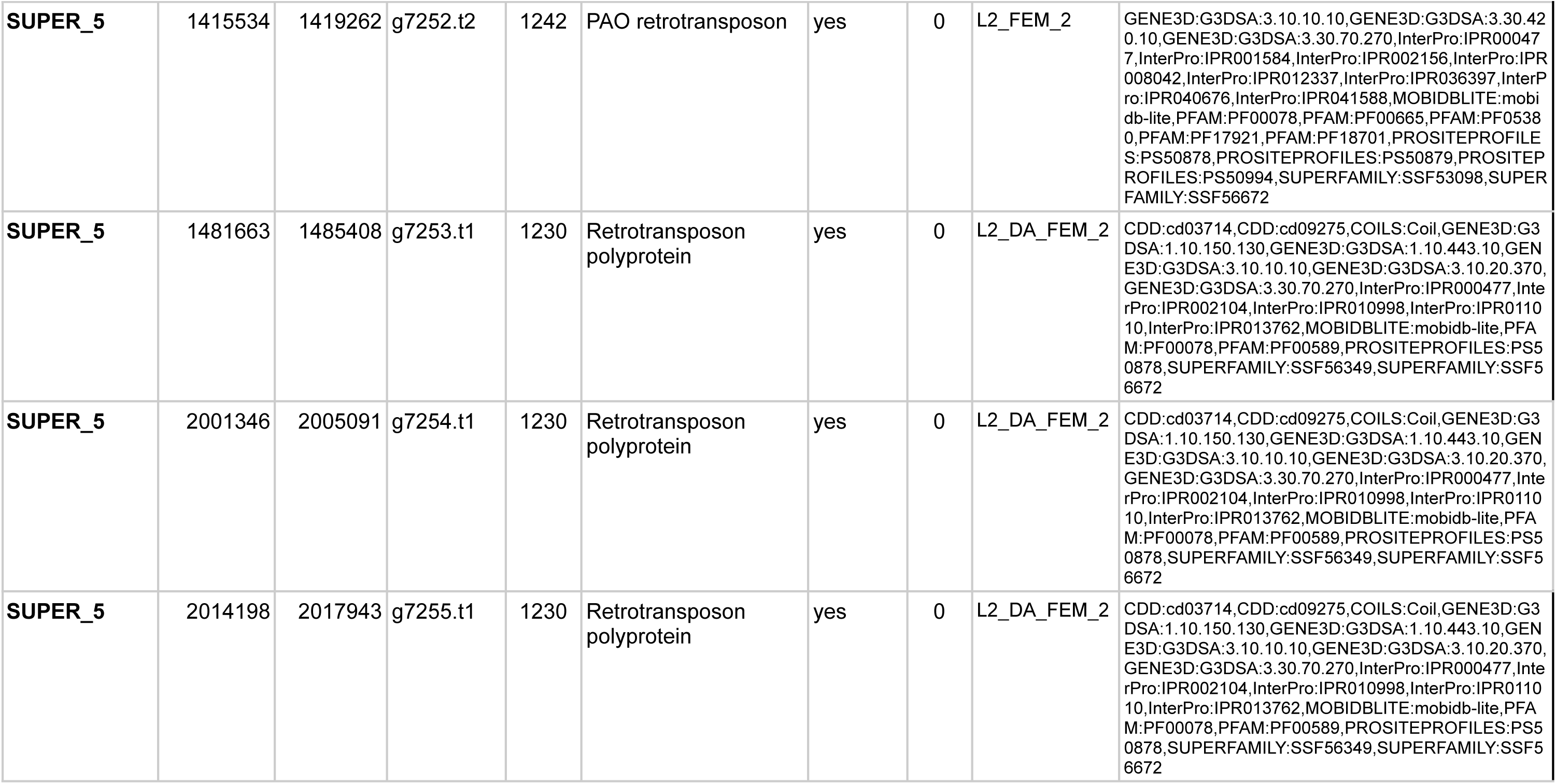
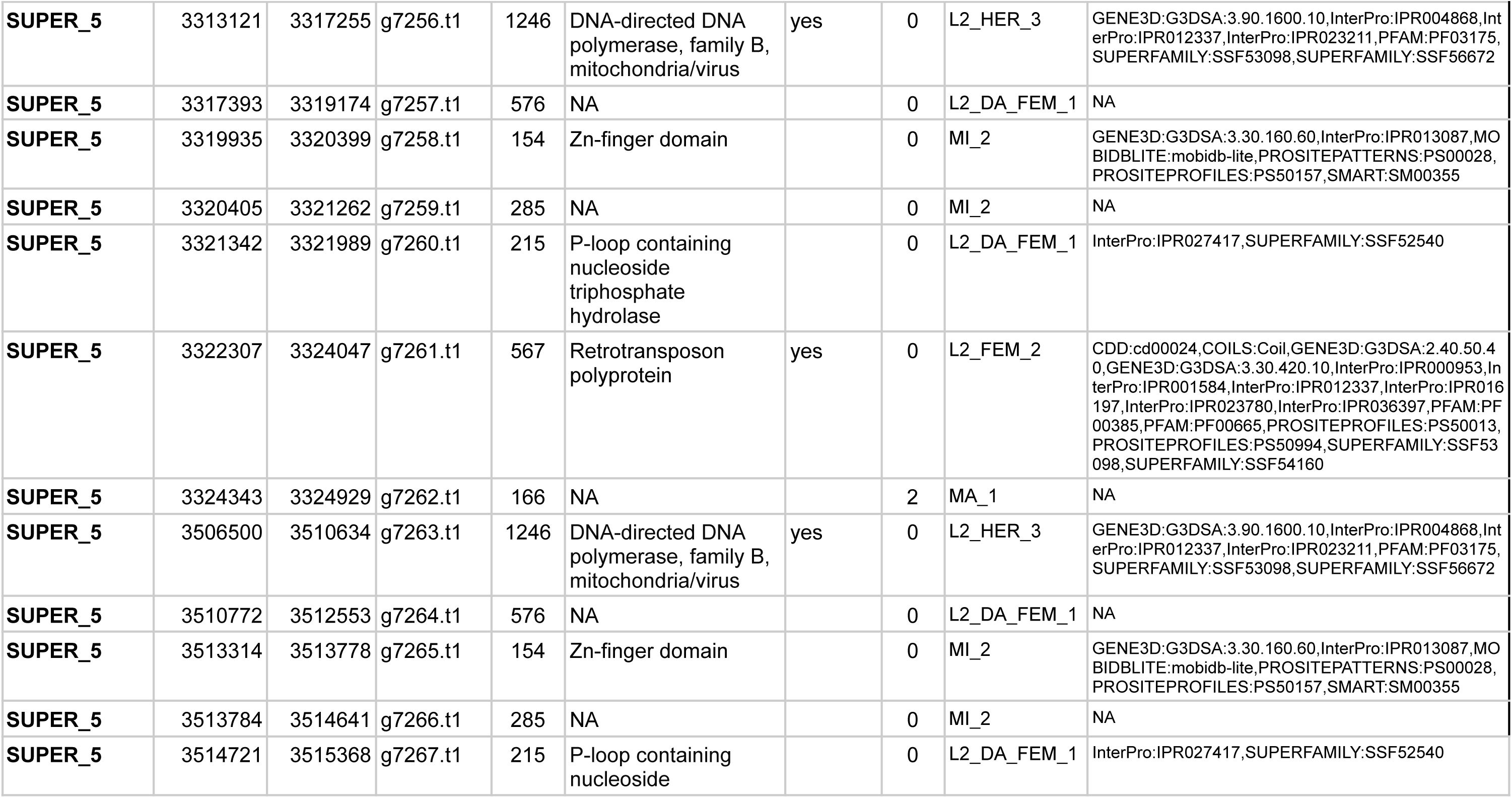
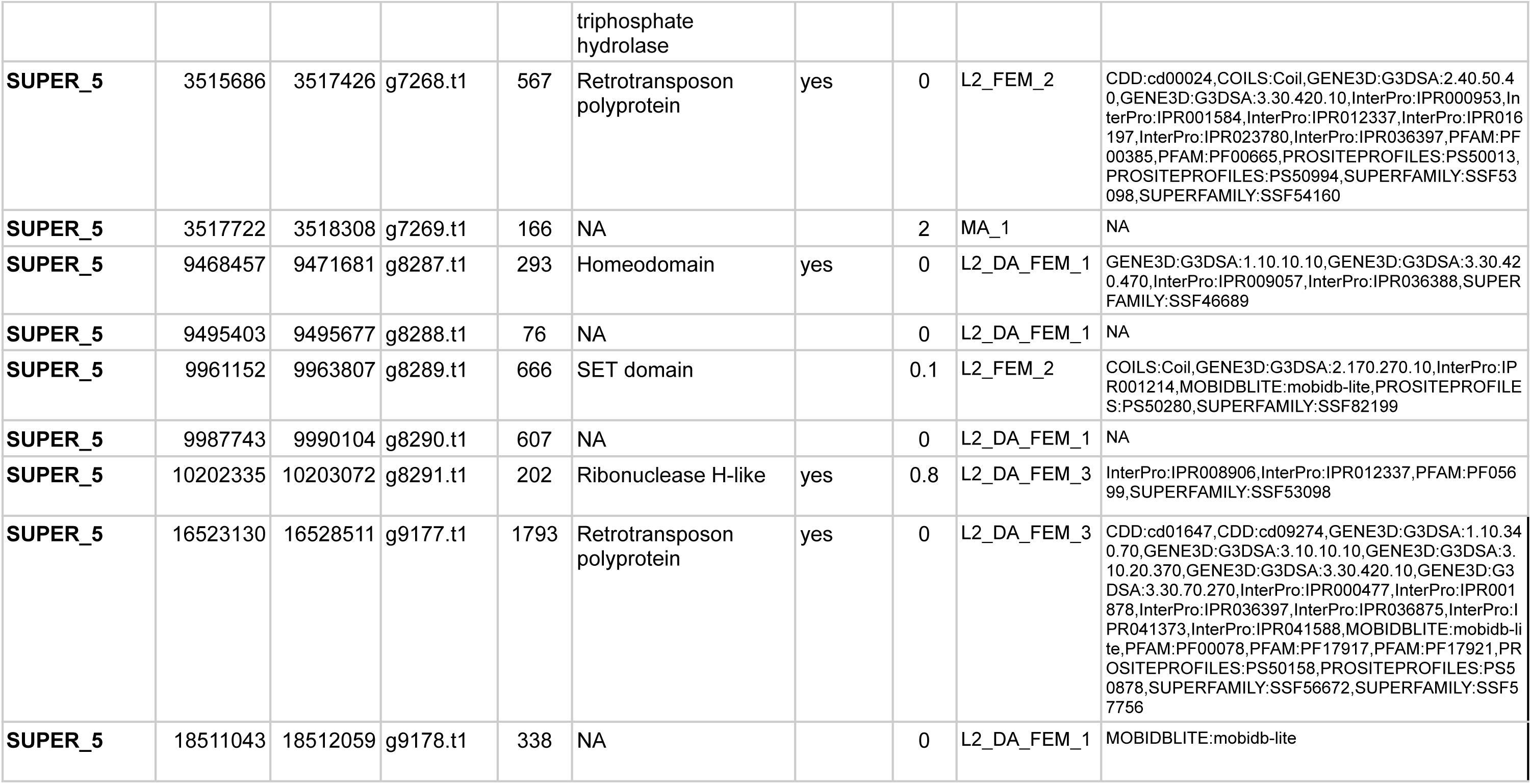
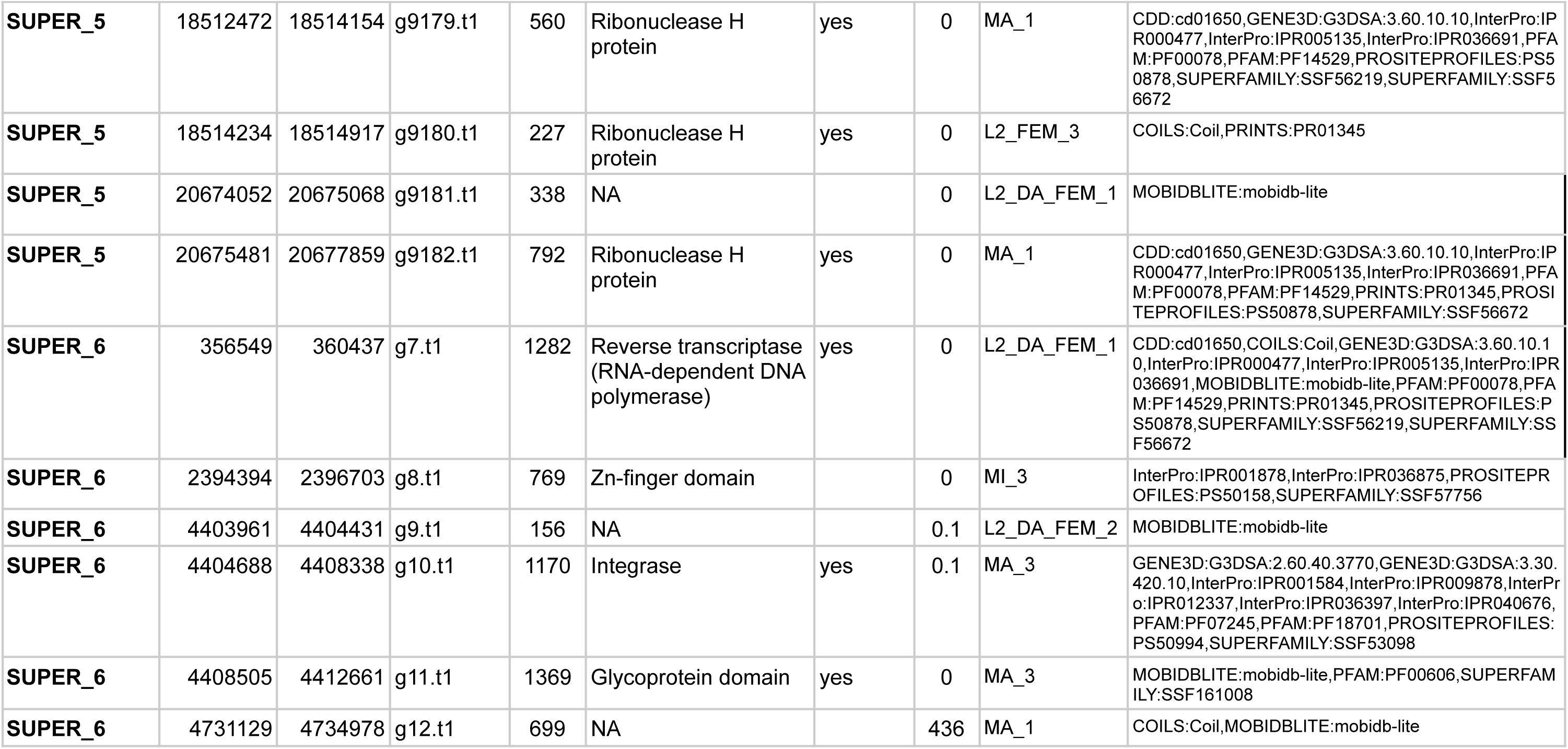
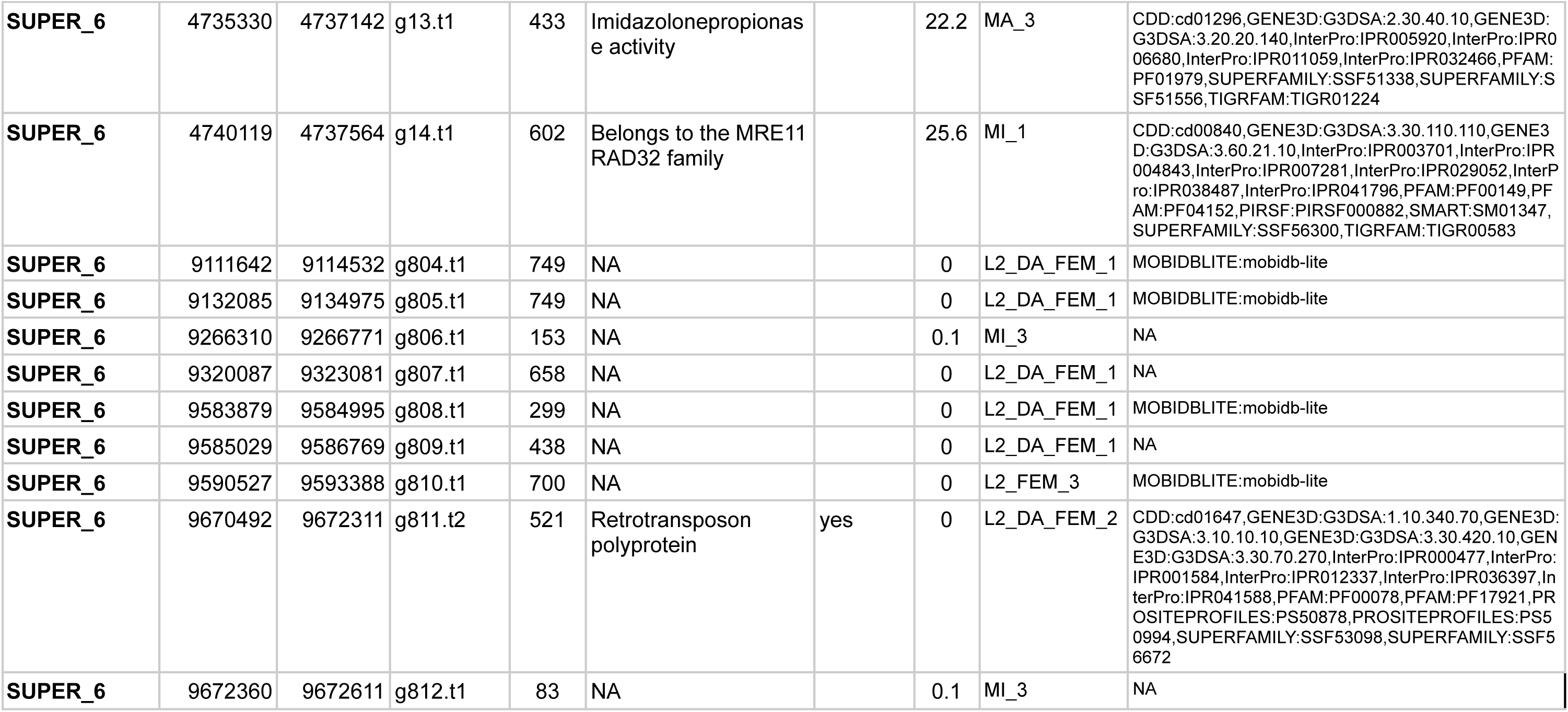
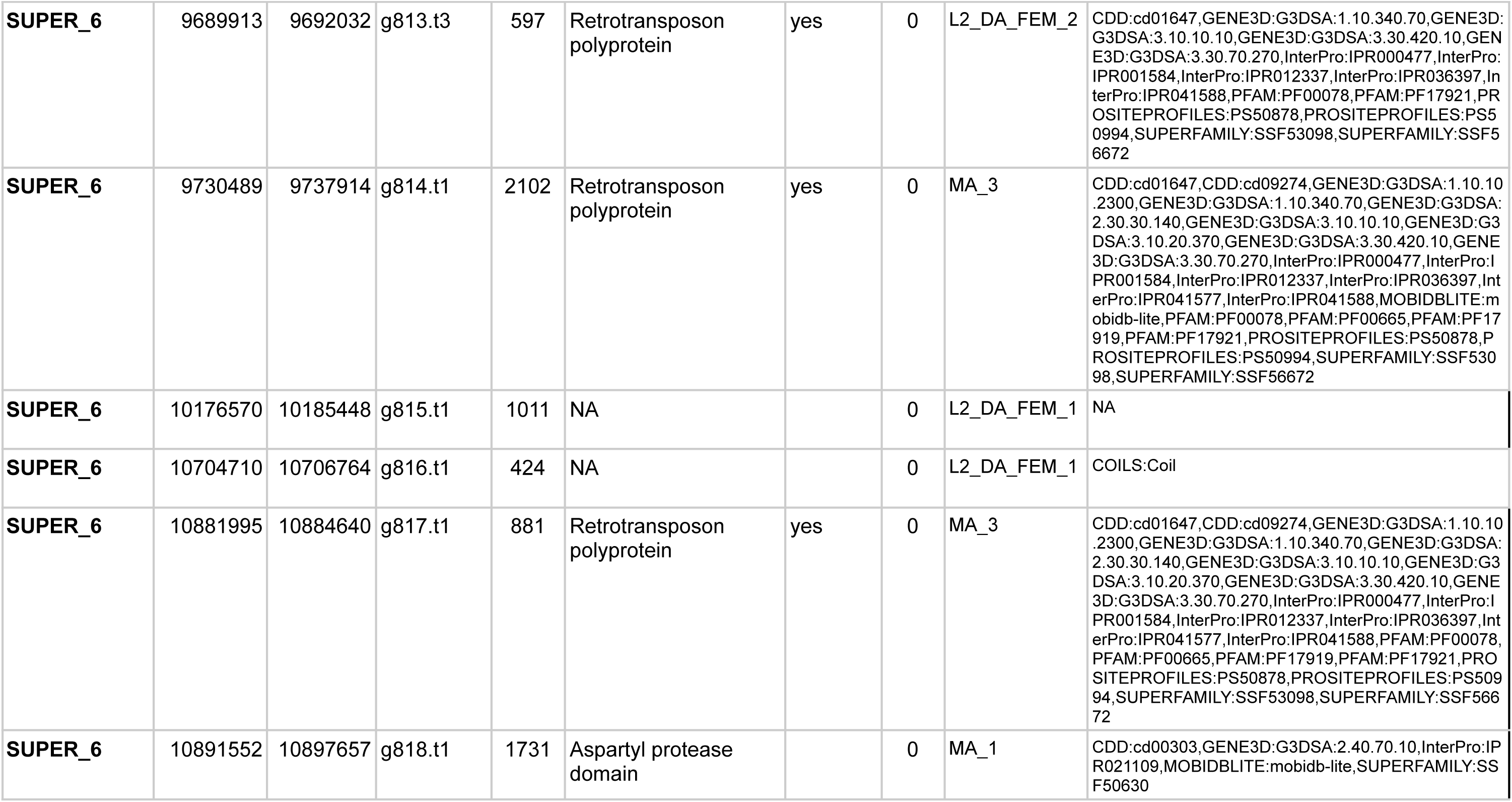
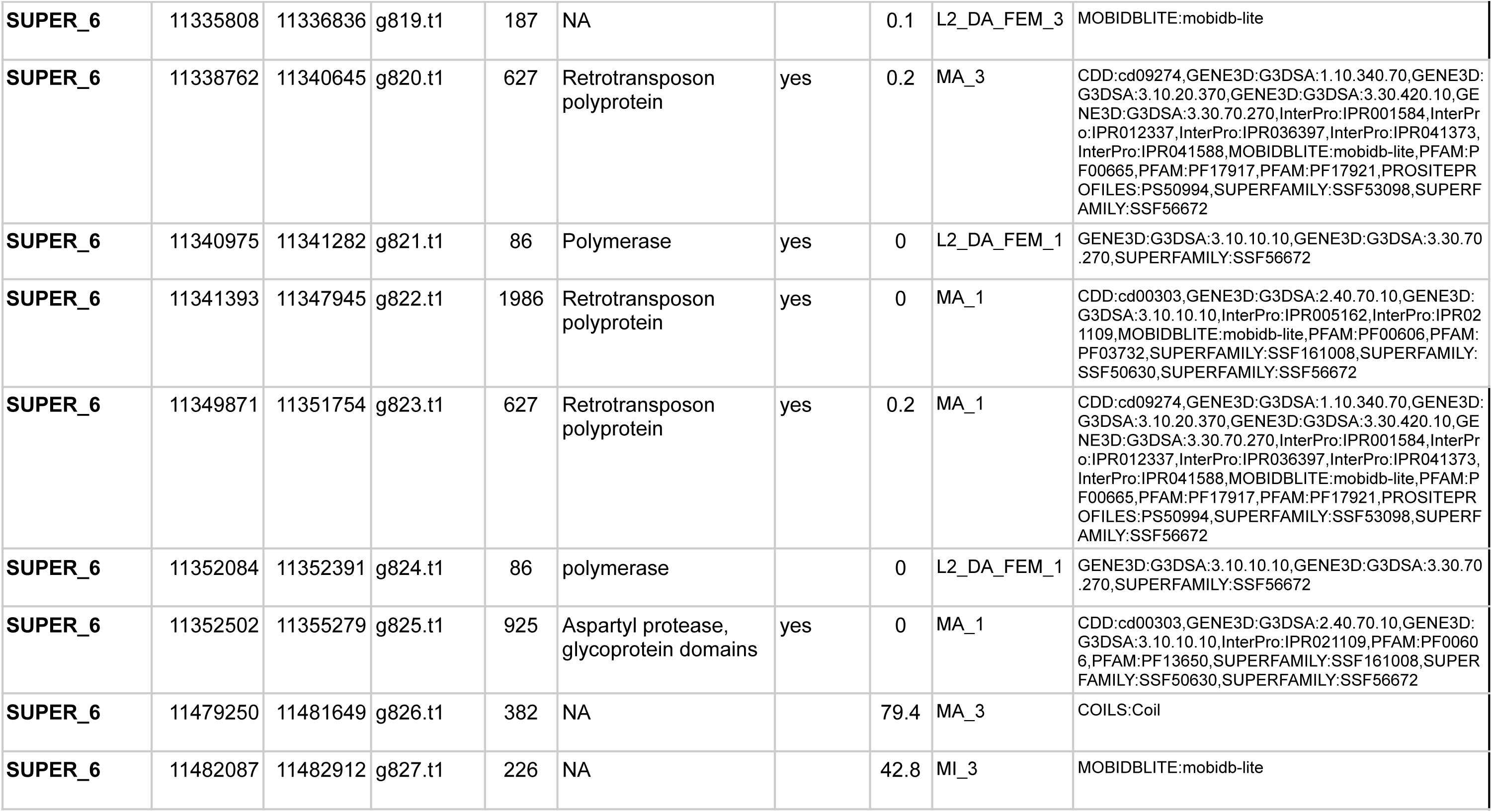
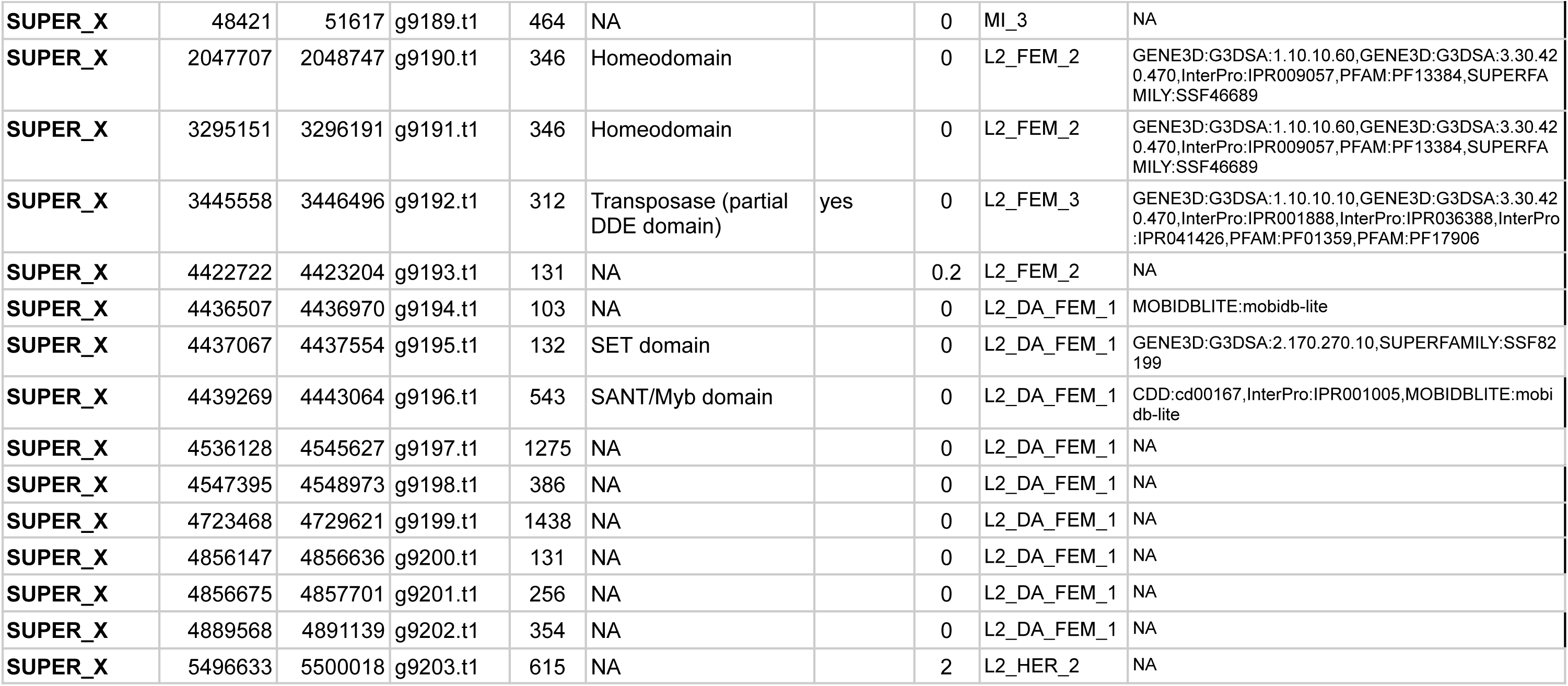
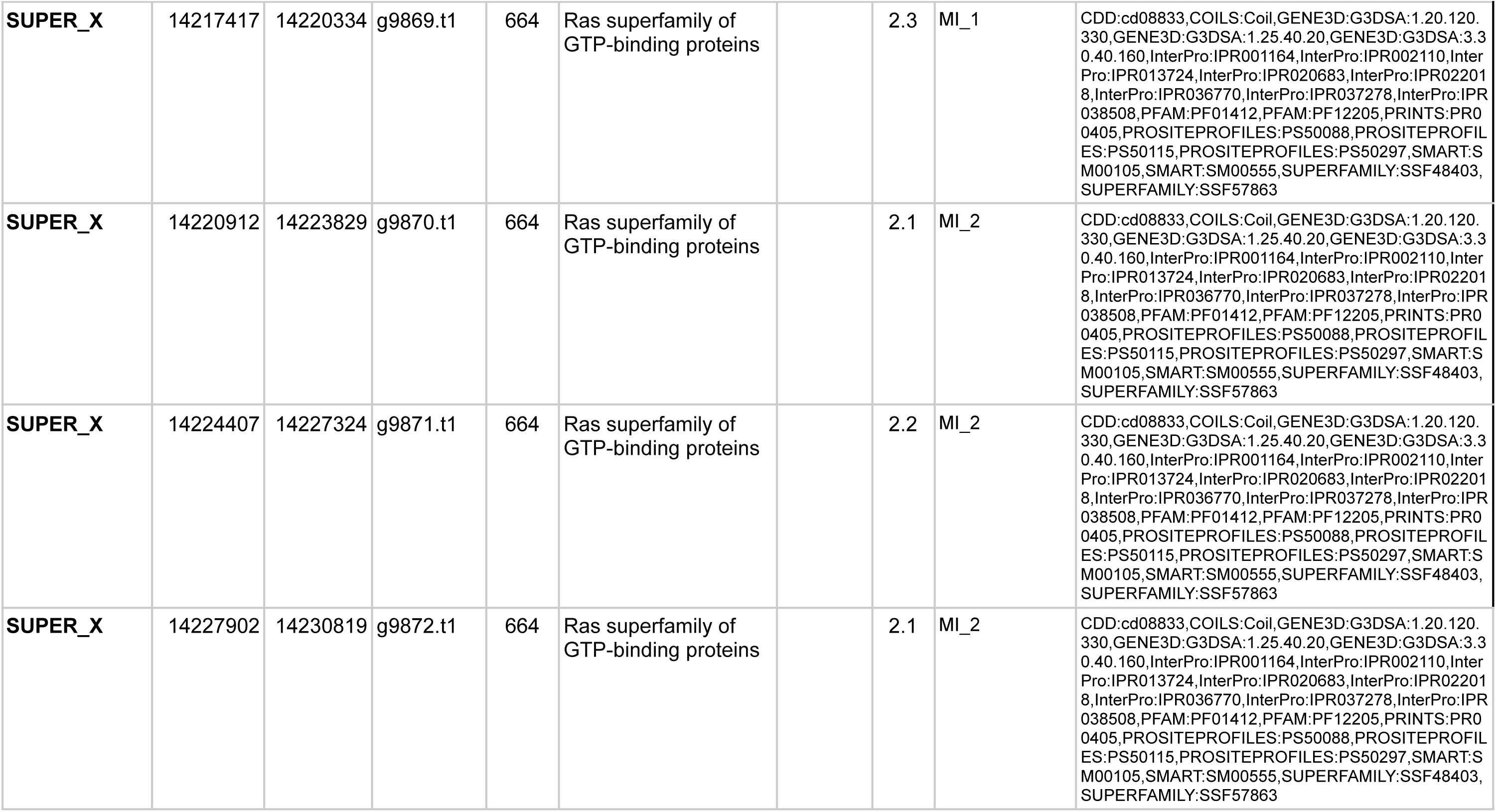
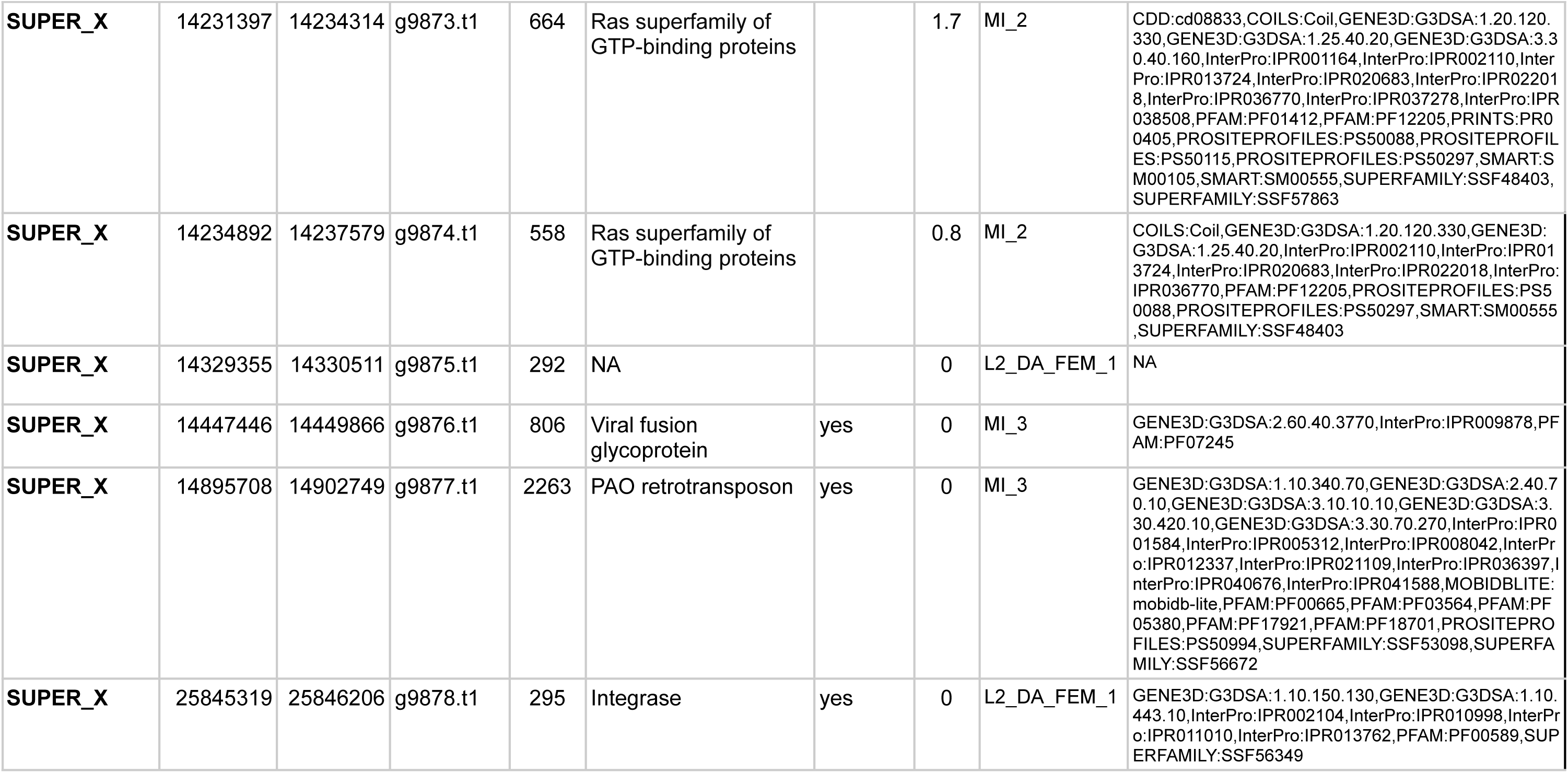
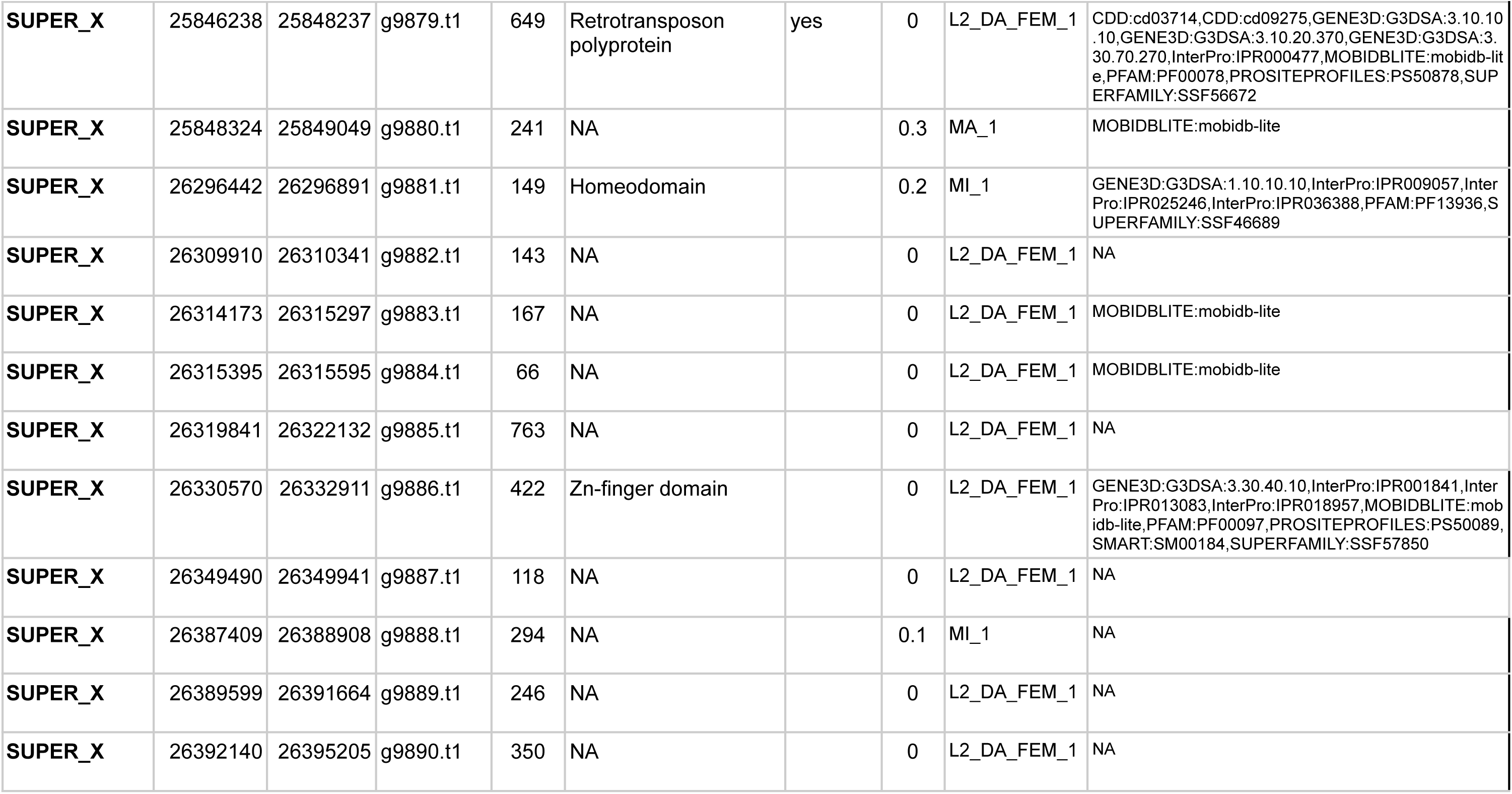
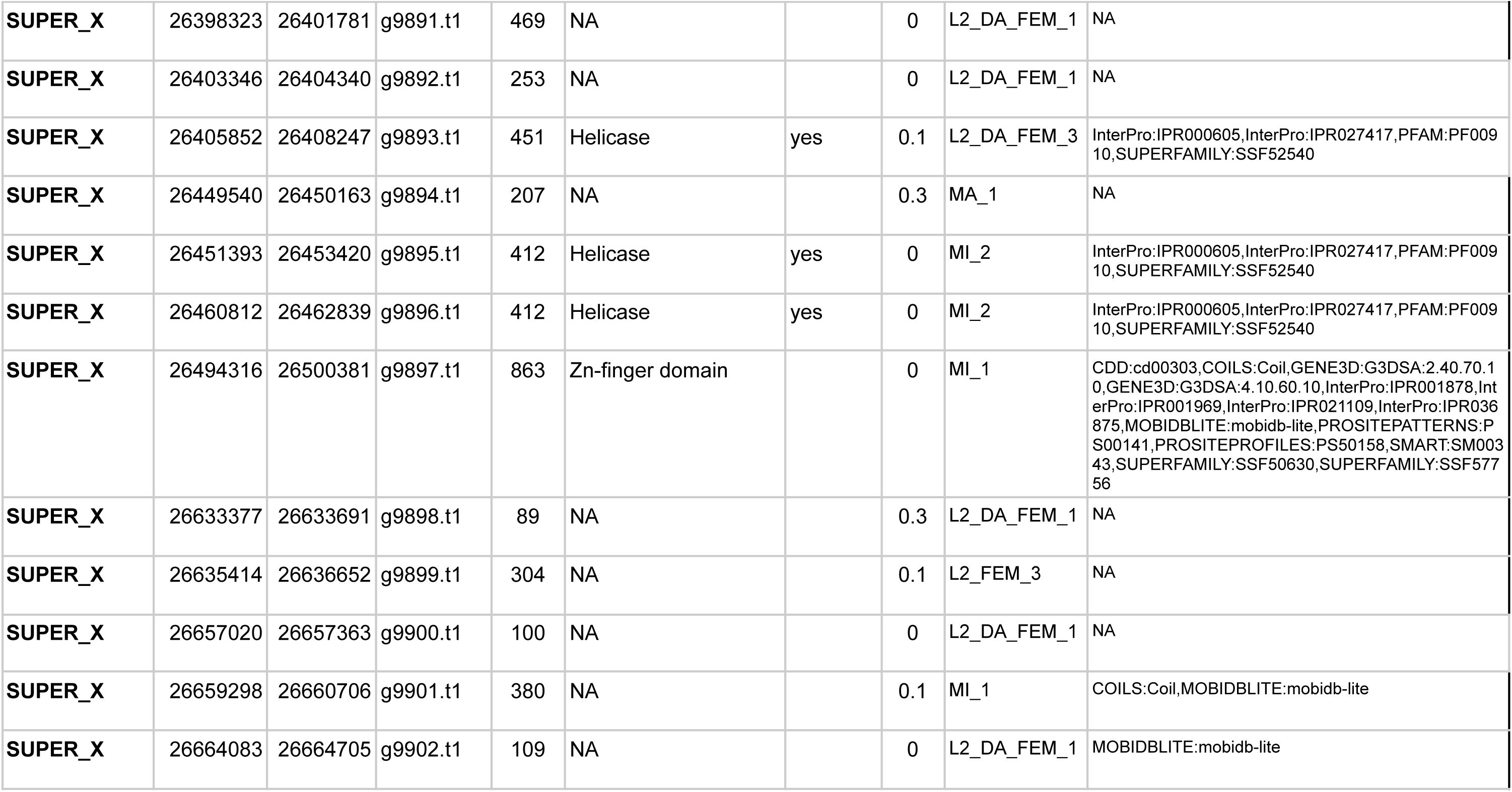
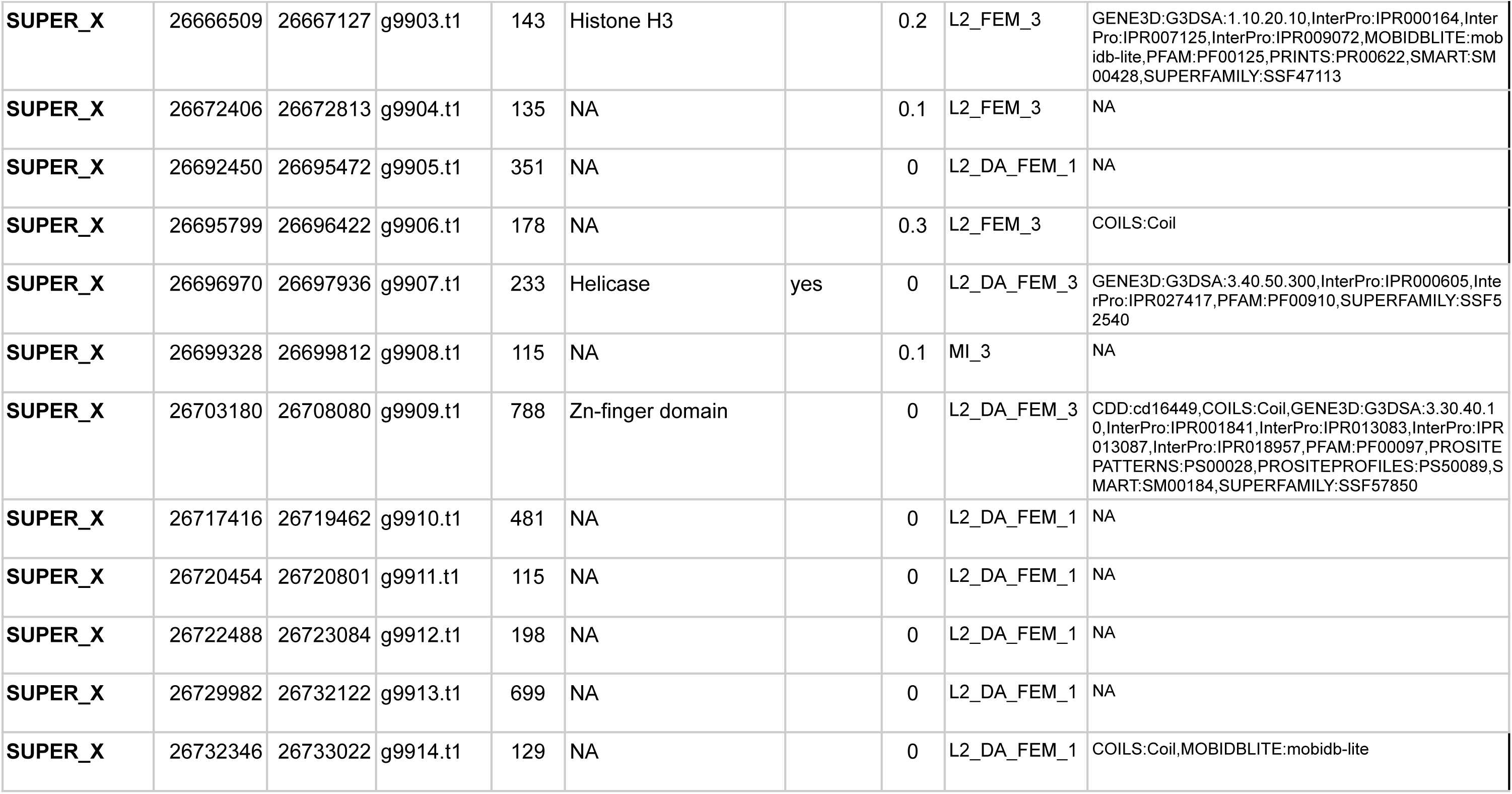
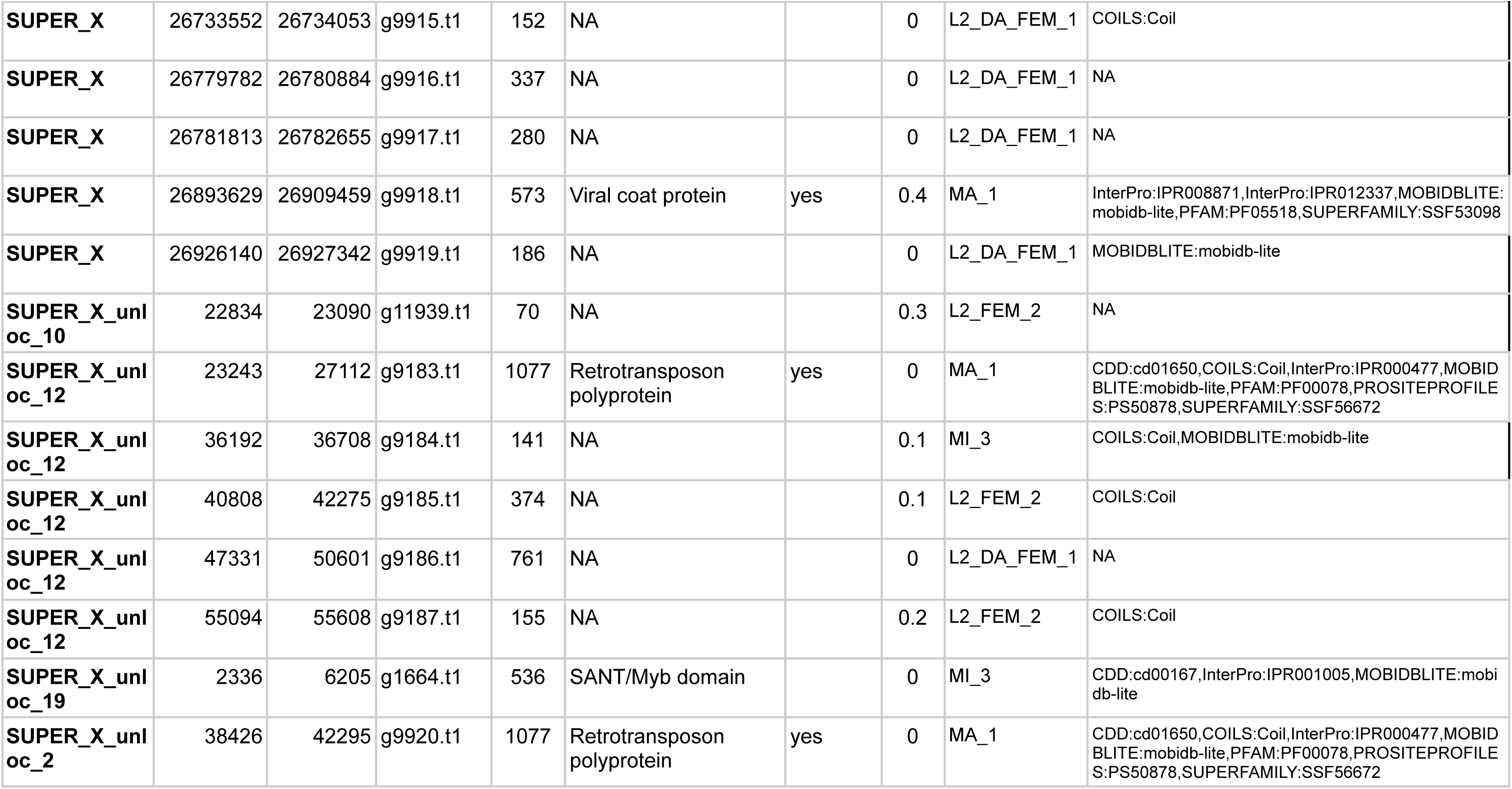
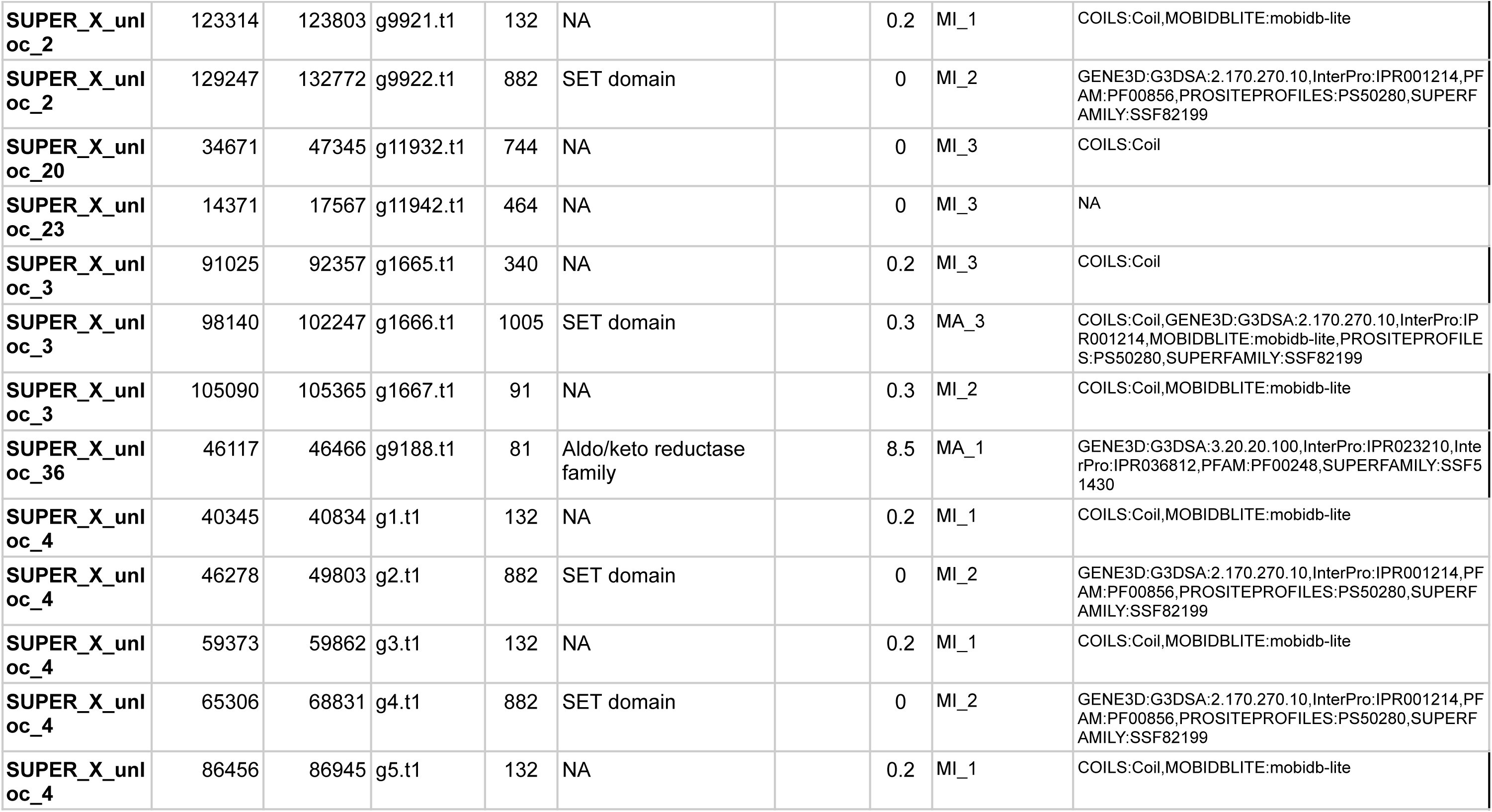
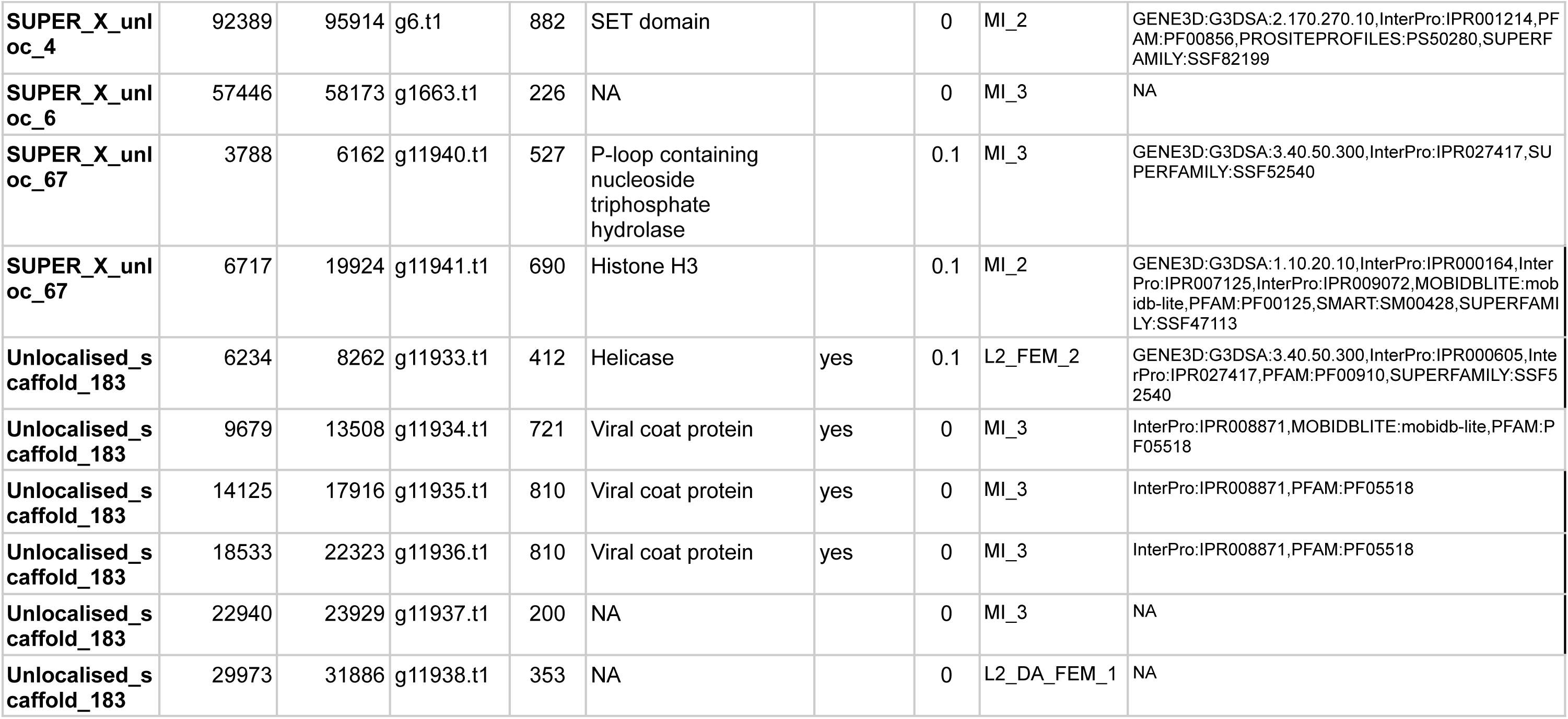
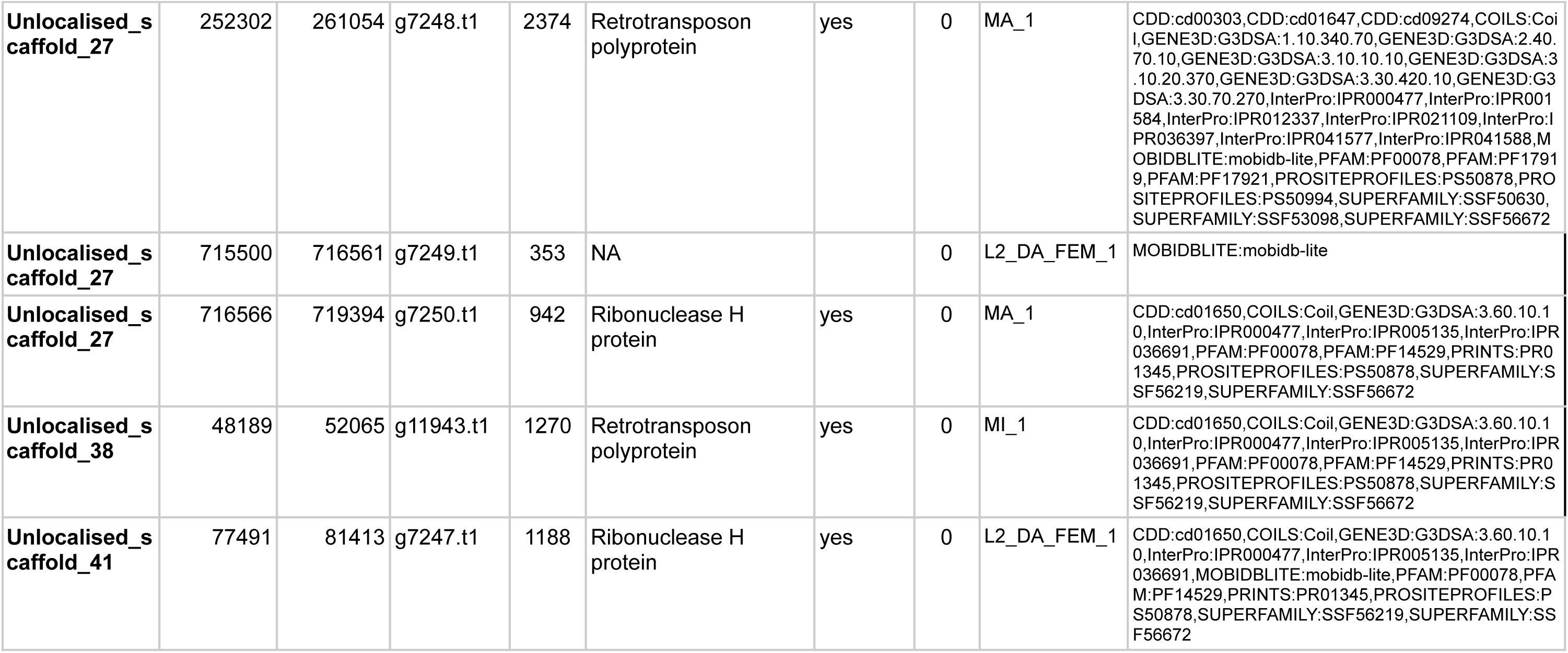
Genes present in eliminated DNA of *Auanema rhodense*. **Column 1:** Under “Sequence name”, “SUPER” stands for chromosomal super scaffold, “SUPER_X_unloc” stands for ulocalised scaffold assigned to SUPER_X (chromosome X). Unlocalised scaffolds were not uniquely associated with any chromosome; all are eliminated. **Column 7:** NA means no functional annotation found; the sources of the annotations are given in **Column 10**. **Column 8:** TPM = transcripts per million mapped reads. **Column 9:** Sample names are described in Table S8.

**Table S6.**
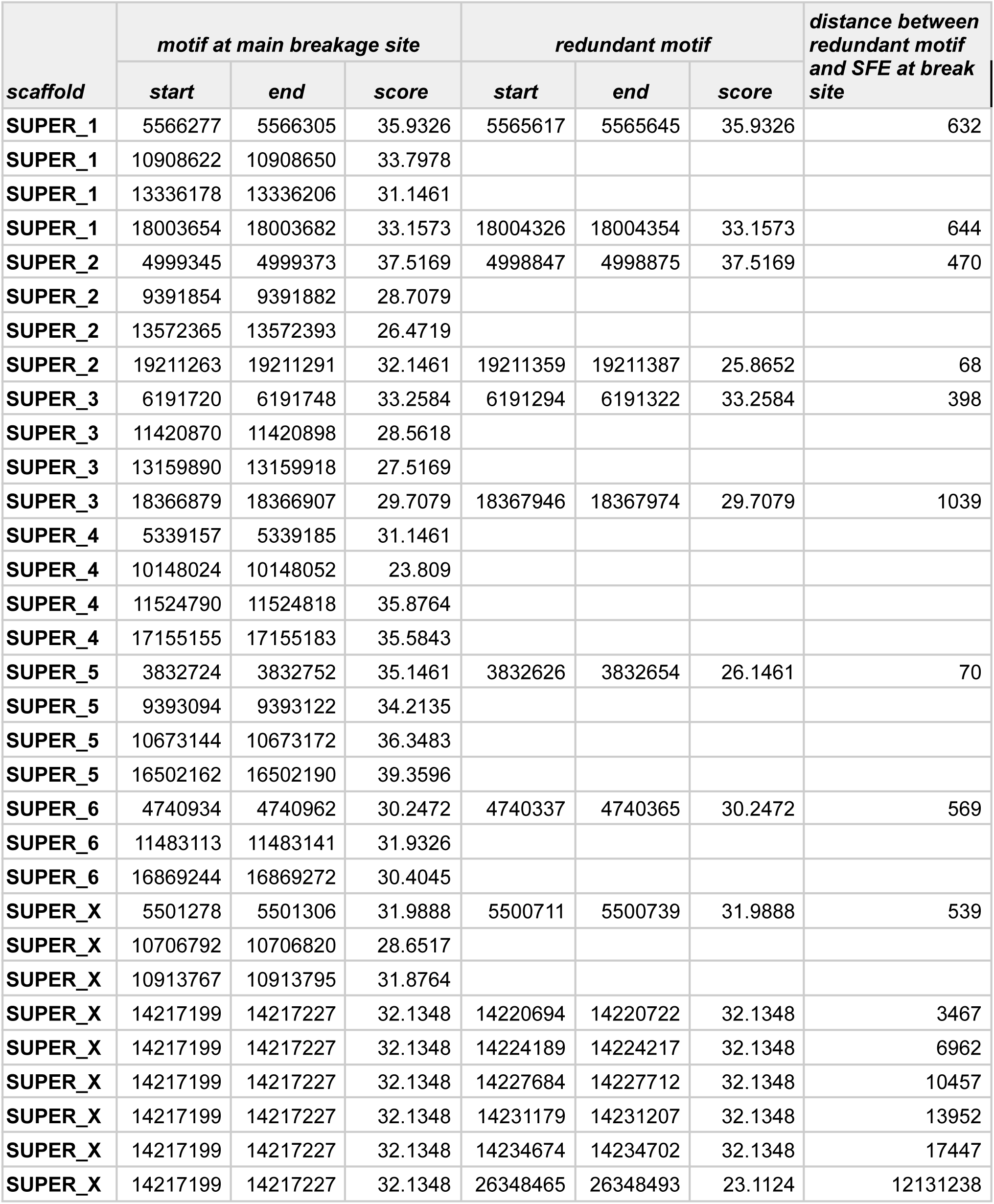
Coordinates of Sequences for Elimination (SFEs) at the breakage sites in *Auanema rhodense* and neighboring SFE occurrences. “SUPER_” is equivalent to “Chromosome”.

**Table S7.**
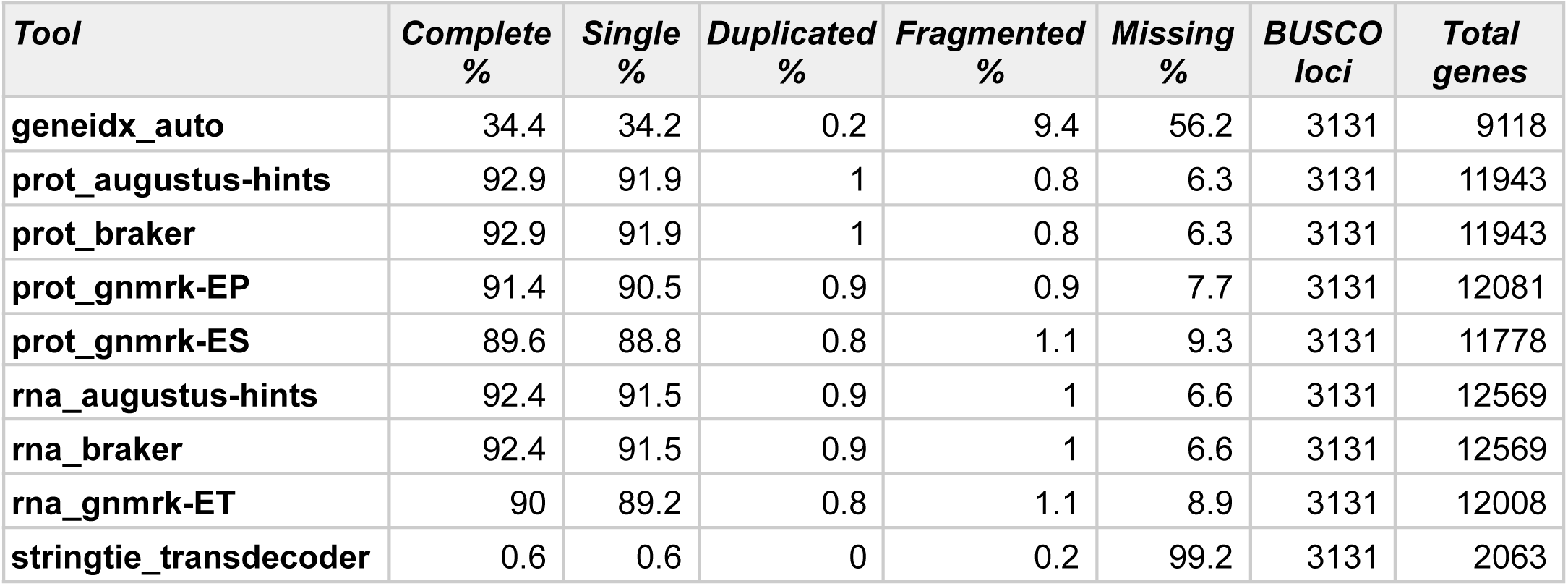
Comparison of *Auanema rhodense* gene finding outcomes assessed with BUSCO Nematoda_odb10.

**Table S8.**
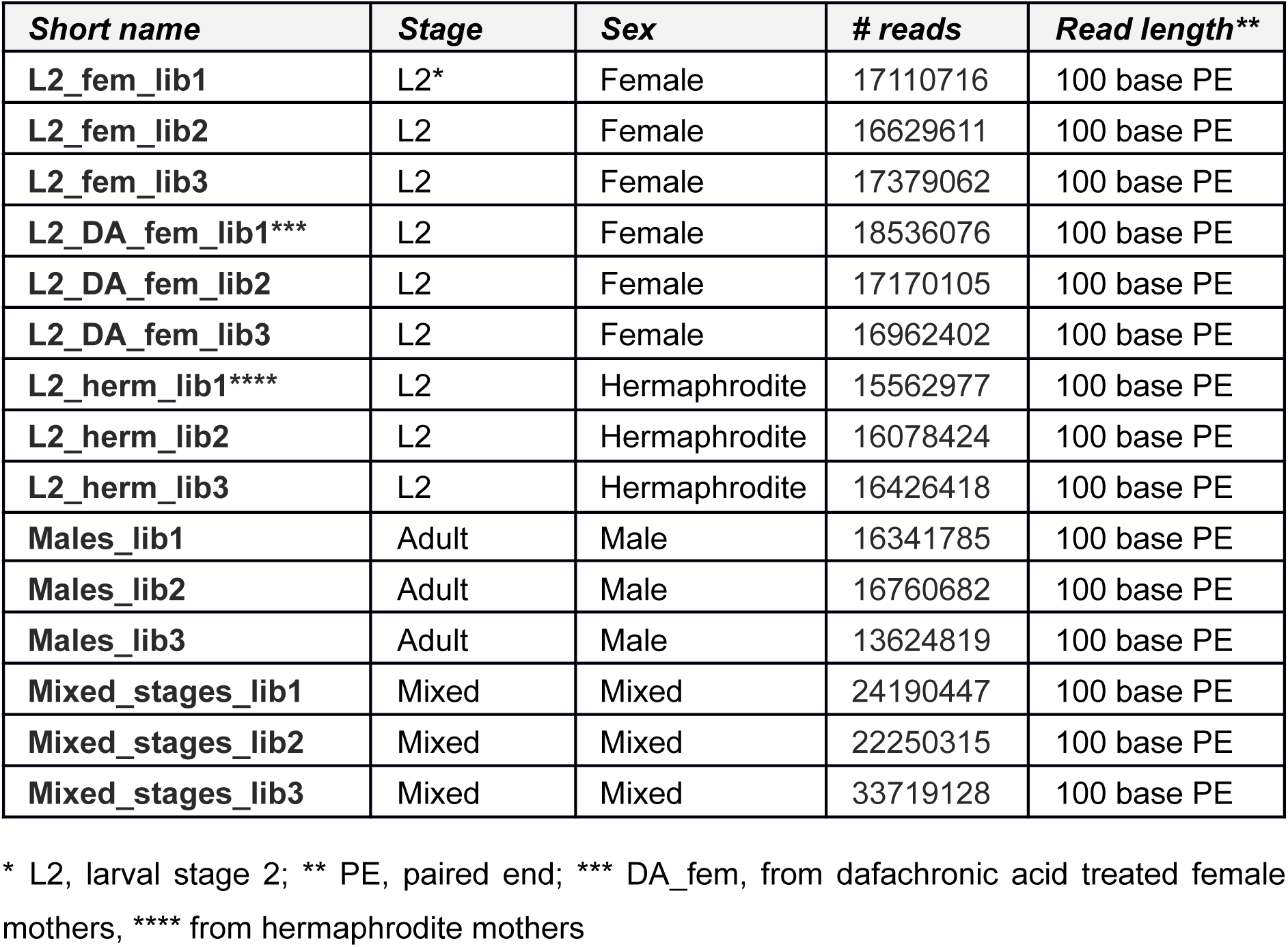
Stage-specific transcriptomic data from Tandonnet, S., Koutsovoulos, G.D., Adams, S., Cloarec, D., Parihar, M., Blaxter, M.L., and Pires-daSilva, A. (2019). Chromosome-Wide Evolution and Sex Determination in the Three-Sexed Nematode *Auanema rhodensis*. G3 9, 1211–1230. https://doi.org/10.1534/g3.119.0011. **ArrayExpress accession E-MTAB-7667** https://www.ebi.ac.uk/biostudies/ArrayExpress/studies/E-MTAB-7667

**Table S9.**
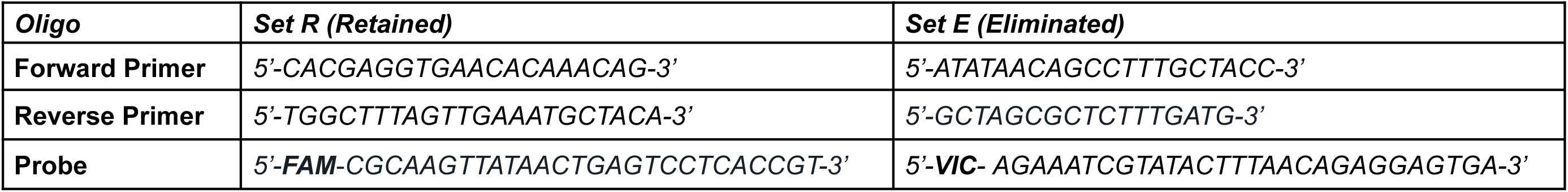
Oligonucleotide sequence of probe and primer sets used for qPCR.

## References

1. Weismann, A. (1892). Das Keimplasma: eine Theorie der Vererbung (Fischer).

2. Cory, S., and Adams, J.M. (1980). Deletions are associated with somatic rearrangement of immunoglobulin heavy chain genes. Cell 19, 37–51. 10.1016/0092-8674(80)90386-4.

3. Schatz, D.G., Oettinger, M.A., and Schlissel, M.S. (1992). V(D)J recombination: molecular biology and regulation. Annu Rev Immunol 10, 359–383. 10.1146/annurev.iy.10.040192.002043.

4. Kim, S., Davis, M., Sinn, E., Patten, P., and Hood, L. (1981). Antibody diversity: somatic hypermutation of rearranged VH genes. Cell 27, 573–581. 10.1016/0092-8674(81)90399-8.

5. Stephens, P.J., Greenman, C.D., Fu, B., Yang, F., Bignell, G.R., Mudie, L.J., Pleasance, E.D., Lau, K.W., Beare, D., Stebbings, L.A., et al. (2011). Massive genomic rearrangement acquired in a single catastrophic event during cancer development. Cell 144, 27–40. 10.1016/j.cell.2010.11.055.

6. Boveri, T. (1887). Uber Differenzierung der Zellkerne wahrend der Furchung des Eies von Ascaris megalocephala. Anat. Anz. 2, 688–693.

7. Wang, J., Gao, S., Mostovoy, Y., Kang, Y., Zagoskin, M., Sun, Y., Zhang, B., White, L.K., Easton, A., Nutman, T.B., et al. (2017). Comparative genome analysis of programmed DNA elimination in nematodes. Genome Res. 27, 2001–2014. 10.1101/gr.225730.117.

8. Wang, J., Veronezi, G.M.B., Kang, Y., Zagoskin, M., O’Toole, E.T., and Davis, R.E. (2020). Comprehensive Chromosome End Remodeling during Programmed DNA Elimination. Curr. Biol. 30, 3397–3413.e4. 10.1016/j.cub.2020.06.058.

9. Wang, J.B., Mitreva, M., Berriman, M., Thorne, A., Magrini, V., Koutsovoulos, G., Kumar, S., Blaxter, M.L., and Davis, R.E. (2012). Silencing of Germline-Expressed Genes by DNA Elimination in Somatic Cells. Dev. Cell 23, 1072–1080. 10.1016/j.devcel.2012.09.020.

10. Müller, F., Bernard, V., and Tobler, H. (1996). Chromatin diminution in nematodes. Bioessays 18, 133–138.

11. Muller, F., and Tobler, H. (2000). Chromatin diminution in the parasitic nematodes Ascaris suum and Parascaris univalens. Int. J. Parasitol. 30, 391–399.

12. Gonzalez de la Rosa, P.M., Thomson, M., Trivedi, U., Tracey, A., Tandonnet, S., and Blaxter, M. (2021). A telomere-to-telomere assembly of Oscheius tipulae and the evolution of rhabditid nematode chromosomes. G3 11, jkaa020.

13. Dockendorff, T.C., Estrem, B., Reed, J., Simmons, J.R., Zadegan, S.B., Zagoskin, M.V., Terta, V., Villalobos, E., Seaberry, E.M., and Wang, J. (2022). The nematode Oscheius tipulae as a genetic model for programmed DNA elimination. Curr Biol 32, 5083–5098.e6. 10.1016/j.cub.2022.10.043.

14. Rey, C., Launay, C., Wenger, E., and Delattre, M. (2023). Programmed DNA elimination in Mesorhabditis nematodes. Curr Biol 33, 3711–3721.e5. 10.1016/j.cub.2023.07.058.

15. Stevens, L., Sun, S., Haruta, N., Xiao, L., Uwatoko, N., Kieninger, M., Sato, K., Yoshida, A., Absolon, D., Collins, J., et al. (2025). Programmed DNA elimination was present in the last common ancestor of Caenorhabditis nematodes. bioRxiv, 2025.10.23.681605. 10.1101/2025.10.23.681605.

16. Launay, C., Wenger, E., Letcher, B., and Delattre, M. (2026). Somatic Programmed DNA Elimination is widespread in free-living Rhabditidae nematodes. eLife. 10.7554/elife.111501.1.

17. Maupas, E. (1900). Modes et formes de reproduction des nématodes. Ann Zool Exp Gen, 463–624.

18. Félix, M.-A. (2004). Alternative morphs and plasticity of vulval development in a rhabditid nematode species. Development Genes and Evolution 214, 55–63. 10.1007/s00427-003-0376-y.

19. Kanzaki, N., Kiontke, K., Tanaka, R., Hirooka, Y., Schwarz, A., Müller-Reichert, T., Chaudhuri, J., and Pires-daSilva, A. (2017). Description of two three-gendered nematode species in the new genus Auanema (Rhabditina) that are models for reproductive mode evolution. Sci. Rep. 7, 11135. 10.1038/s41598-017-09871-1.

20. Adams, S., Tandonnet, S., and Pires-daSilva, A. (2025). Balancing selfing and outcrossing: the genetics and cell biology of nematodes with three sexual morphs. Genetics 229. 10.1093/genetics/iyae173.

21. Chaudhuri, J., Bose, N., Tandonnet, S., Adams, S., Zuco, G., Kache, V., Parihar, M., von Reuss, S.H., Schroeder, F.C., and Pires-daSilva, A. (2015). Mating dynamics in a nematode with three sexes and its evolutionary implications. Sci Rep 5, 17676. 10.1038/srep17676.

22. Tandonnet, S., Farrell, M.C., Koutsovoulos, G.D., Blaxter, M.L., Parihar, M., Sadler, P.L., Shakes, D.C., and Pires-daSilva, A. (2017). Sex- and Gamete-Specific Patterns of X Chromosome Segregation in a Trioecious Nematode. Curr. Biol. 28, 93–99 e3. 10.1016/j.cub.2017.11.037.

23. Tandonnet, S., Koutsovoulos, G.D., Adams, S., Cloarec, D., Parihar, M., Blaxter, M.L., and Pires-daSilva, A. (2019). Chromosome-Wide Evolution and Sex Determination in the Three-Sexed Nematode Auanema rhodensis. G3 9, 1211–1230. 10.1534/g3.119.0011.

24. Manni, M., Berkeley, M.R., Seppey, M., Simão, F.A., and Zdobnov, E.M. (2021). BUSCO Update: Novel and Streamlined Workflows along with Broader and Deeper Phylogenetic Coverage for Scoring of Eukaryotic, Prokaryotic, and Viral Genomes. Mol. Biol. Evol. 38, 4647–4654. 10.1093/molbev/msab199.

25. Girgis, H.Z. (2015). Red: an intelligent, rapid, accurate tool for detecting repeats de-novo on the genomic scale. BMC Bioinformatics 16, 227. 10.1186/s12859-015-0654-5.

26. MacQueen, A.J., Phillips, C.M., Bhalla, N., Weiser, P., Villeneuve, A.M., and Dernburg, A.F. (2005). Chromosome Sites Play Dual Roles to Establish Homologous Synapsis during Meiosis in C. elegans. Preprint, 10.1016/j.cell.2005.09.034 10.1016/j.cell.2005.09.034.

27. Phillips, C.M., and Dernburg, A.F. (2006). A family of zinc-finger proteins is required for chromosome-specific pairing and synapsis during meiosis in C. elegans. Dev. Cell 11, 817–829. 10.1016/j.devcel.2006.09.020.

28. Phillips, C.M., Meng, X., Zhang, L., Chretien, J.H., Urnov, F.D., and Dernburg, A.F. (2009). Identification of chromosome sequence motifs that mediate meiotic pairing and synapsis in C. elegans. Nat. Cell Biol. 11, 934–942. 10.1038/ncb1904.

29. Hillers, K.J., Jantsch, V., Martinez-Perez, E., and Yanowitz, J.L. (2017). Meiosis. WormBook 2017, 1–43. 10.1895/wormbook.1.178.1.

30. Phillips, C.M., Wong, C., Bhalla, N., Carlton, P.M., Weiser, P., Meneely, P.M., and Dernburg, A.F. (2005). HIM-8 binds to the X chromosome pairing center and mediates chromosome-specific meiotic synapsis. Cell 123, 1051–1063. 10.1016/j.cell.2005.09.035.

31. Felix, M.A. (2004). Alternative morphs and plasticity of vulval development in a rhabditid nematode species. Dev. Genes Evol. 214, 55–63.

32. Hope, I.A. ed. (1999). C. elegans - a practical approach (Oxford Univerisity Press).

33. Ranallo-Benavidez, T.R., Jaron, K.S., and Schatz, M.C. (2020). GenomeScope 2.0 and Smudgeplot for reference-free profiling of polyploid genomes. Nat. Commun. 11, 1432. 10.1038/s41467-020-14998-3.

34. Cheng, H., Concepcion, G.T., Feng, X., Zhang, H., and Li, H. (2021). Haplotype-resolved de novo assembly using phased assembly graphs with hifiasm. Nat. Methods 18, 170–175. 10.1038/s41592-020-01056-5.

35. Guan, D., McCarthy, S.A., Wood, J., Howe, K., Wang, Y., and Durbin, R. (2020). Identifying and removing haplotypic duplication in primary genome assemblies. Bioinformatics 36, 2896–2898. 10.1093/bioinformatics/btaa025.

36. Challis, R., Richards, E., Rajan, J., Cochrane, G., and Blaxter, M. (2020). BlobToolKit - Interactive Quality Assessment of Genome Assemblies. G3 *10*, 1361–1374. 10.1534/g3.119.400908.

37. Uliano-Silva, M., Gabriel R. N. Ferreira, J., Krasheninnikova, K., Formenti, G., Abueg, L., Torrance, J., W Myers, G., Durbin, R., Blaxter, M. A. McCarthy, S., et al. (2022). MitoHiFi: a python pipeline for mitochondrial genome assembly from PacBio High Fidelity reads. bioRxiv. 10.1101/2022.12.23.521667.

38. Zhou, C., McCarthy, S.A., and Durbin, R. (2023). YaHS: yet another Hi-C scaffolding tool. Bioinformatics 39. 10.1093/bioinformatics/btac808.

39. Howe, K., Chow, W., Collins, J., Pelan, S., Pointon, D.-L., Sims, Y., Torrance, J., Tracey, A., and Wood, J. (2021). Significantly improving the quality of genome assemblies through curation. Gigascience 10. 10.1093/gigascience/giaa153.

40. Steinbiss, S., Willhoeft, U., Gremme, G., and Kurtz, S. (2009). Fine-grained annotation and classification of de novo predicted LTR retrotransposons. Nucleic Acids Res. 37, 7002–7013. 10.1093/nar/gkp759.

41. Mistry, J., Finn, R.D., Eddy, S.R., Bateman, A., and Punta, M. (2013). Challenges in homology search: HMMER3 and convergent evolution of coiled-coil regions. Nucleic Acids Res. 41, e121. 10.1093/nar/gkt263.

42. Flynn, J.M., Hubley, R., Goubert, C., Rosen, J., Clark, A.G., Feschotte, C., and Smit, A.F. (2020). RepeatModeler2 for automated genomic discovery of transposable element families. Proc. Natl. Acad. Sci. U. S. A. 117, 9451–9457. 10.1073/pnas.1921046117.

43. Wheeler, T.J., Clements, J., Eddy, S.R., Hubley, R., Jones, T.A., Jurka, J., Smit, A.F., and Finn, R.D. (2013). Dfam: a database of repetitive DNA based on profile hidden Markov models. Nucleic Acids Res. 41, D70–D82. 10.1093/nar/gks1265.

44. Rognes, T., Flouri, T., Nichols, B., Quince, C., and Mahé, F. (2016). VSEARCH: a versatile open source tool for metagenomics. PeerJ 4, e2584. 10.7717/peerj.2584.

45. Howe, K.L., Bolt, B.J., Shafie, M., Kersey, P., and Berriman, M. (2017). WormBase ParaSite - a comprehensive resource for helminth genomics. Mol. Biochem. Parasitol. 215, 2–10. 10.1016/j.molbiopara.2016.11.005.

46. Camacho, C., Coulouris, G., Avagyan, V., Ma, N., Papadopoulos, J., Bealer, K., and Madden, T.L. (2009). BLAST+: architecture and applications. BMC Bioinformatics 10, 421. 10.1186/1471-2105-10-421.

47. Benson, G. (1999). Tandem repeats finder: a program to analyze DNA sequences. Nucleic Acids Res. 27, 573–580. 10.1093/nar/27.2.573.

48. Chan, P.P., Lin, B.Y., Mak, A.J., and Lowe, T.M. (2021). tRNAscan-SE 2.0: improved detection and functional classification of transfer RNA genes. Nucleic Acids Res. 49, 9077–9096. 10.1093/nar/gkab688.

49. Lagesen, K., Hallin, P., Rødland, E.A., Staerfeldt, H.-H., Rognes, T., and Ussery, D.W. (2007). RNAmmer: consistent and rapid annotation of ribosomal RNA genes. Nucleic Acids Res. 35, 3100–3108. 10.1093/nar/gkm160.

50. Nawrocki, E.P., Kolbe, D.L., and Eddy, S.R. (2009). Infernal 1.0: inference of RNA alignments. Bioinformatics 25, 1335–1337. 10.1093/bioinformatics/btp157.

51. Kalvari, I., Nawrocki, E.P., Ontiveros-Palacios, N., Argasinska, J., Lamkiewicz, K., Marz, M., Griffiths-Jones, S., Toffano-Nioche, C., Gautheret, D., Weinberg, Z., et al. (2021). Rfam 14: expanded coverage of metagenomic, viral and microRNA families. Nucleic Acids Res. 49, D192–D200. 10.1093/nar/gkaa1047.

52. Lawrence, M., Gentleman, R., and Carey, V. (2009). rtracklayer: an R package for interfacing with genome browsers. Bioinformatics 25, 1841–1842. 10.1093/bioinformatics/btp328.

53. Quinlan, A.R., and Hall, I.M. (2010). BEDTools: a flexible suite of utilities for comparing genomic features. Bioinformatics 26, 841–842. 10.1093/bioinformatics/btq033.

54. Lawrence, M., Huber, W., Pagès, H., Aboyoun, P., Carlson, M., Gentleman, R., Morgan, M.T., and Carey, V.J. (2013). Software for Computing and Annotating Genomic Ranges. Preprint, 10.1371/journal.pcbi.1003118 10.1371/journal.pcbi.1003118.

55. Brůna, T., Hoff, K.J., Lomsadze, A., Stanke, M., and Borodovsky, M. (2021). BRAKER2: automatic eukaryotic genome annotation with GeneMark-EP+ and AUGUSTUS supported by a protein database. NAR Genom Bioinform 3, lqaa108. 10.1093/nargab/lqaa108.

56. Dobin, A., Davis, C.A., Schlesinger, F., Drenkow, J., Zaleski, C., Jha, S., Batut, P., Chaisson, M., and Gingeras, T.R. (2013). STAR: ultrafast universal RNA-seq aligner. Bioinformatics 29, 15–21. 10.1093/bioinformatics/bts635.

57. Stanke, M., and Waack, S. (2003). Gene prediction with a hidden Markov model and a new intron submodel. Preprint, 10.1093/bioinformatics/btg1080 10.1093/bioinformatics/btg1080.

58. Brůna, T., Lomsadze, A., and Borodovsky, M. (2020). GeneMark-EP+: eukaryotic gene prediction with self-training in the space of genes and proteins. NAR Genom Bioinform 2, lqaa026. 10.1093/nargab/lqaa026.

59. Paysan-Lafosse, T., Blum, M., Chuguransky, S., Grego, T., Pinto, B.L., Salazar, G.A., Bileschi, M.L., Bork, P., Bridge, A., Colwell, L., et al. (2022). InterPro in 2022. Nucleic Acids Res. 10.1093/nar/gkac993.

60. Jones, P., Binns, D., Chang, H.Y., Fraser, M., Li, W., McAnulla, C., McWilliam, H., Maslen, J., Mitchell, A., Nuka, G., et al. (2014). InterProScan 5: genome-scale protein function classification. Bioinformatics 30, 1236–1240. 10.1093/bioinformatics/btu031.

61. Cantalapiedra, C.P., Hernández-Plaza, A., Letunic, I., Bork, P., and Huerta-Cepas, J. (2021). eggNOG-mapper v2: Functional Annotation, Orthology Assignments, and Domain Prediction at the Metagenomic Scale. Mol. Biol. Evol. 38, 5825–5829. 10.1093/molbev/msab293.

62. Li, H. (2018). Minimap2: pairwise alignment for nucleotide sequences. Bioinformatics 34, 3094–3100. 10.1093/bioinformatics/bty191.

63. Bonfield, J.K., and Whitwham, A. (2010). Gap5--editing the billion fragment sequence assembly. Bioinformatics 26, 1699–1703. 10.1093/bioinformatics/btq268.

64. Seah, B.K.B., and Swart, E.C. (2021). BleTIES: Annotation of natural genome editing in ciliates using long read sequencing. Bioinformatics. 10.1093/bioinformatics/btab613.

65. Bailey, T.L., Johnson, J., Grant, C.E., and Noble, W.S. (2015). The MEME Suite. Preprint, 10.1093/nar/gkv416 10.1093/nar/gkv416.

66. Bailey, T.L., and Elkan, C. (1995). Unsupervised learning of multiple motifs in biopolymers using expectation maximization. Preprint, 10.1007/bf00993379 10.1007/bf00993379.

67. Grant, C.E., Bailey, T.L., and Noble, W.S. (2011). FIMO: scanning for occurrences of a given motif. Bioinformatics 27, 1017–1018. 10.1093/bioinformatics/btr064.

68. Ou, J., Wolfe, S.A., Brodsky, M.H., and Zhu, L.J. (2018). motifStack for the analysis of transcription factor binding site evolution. Nat. Methods 15, 8–9. 10.1038/nmeth.4555.

69. Wagih, O. (2017). ggseqlogo: a versatile R package for drawing sequence logos. Bioinformatics 33, 3645–3647. 10.1093/bioinformatics/btx469.

70. Lee, R.Y.N., Howe, K.L., Harris, T.W., Arnaboldi, V., Cain, S., Chan, J., Chen, W.J., Davis, P., Gao, S., Grove, C., et al. (2018). WormBase 2017: molting into a new stage. Nucleic Acids Res 46, D869–D874. 10.1093/nar/gkx998.

71. Emms, D.M., and Kelly, S. (2015). OrthoFinder: solving fundamental biases in whole genome comparisons dramatically improves orthogroup inference accuracy. Genome Biol. 16, 157. 10.1186/s13059-015-0721-2.

72. Buchfink, B., Reuter, K., and Drost, H.-G. (2021). Sensitive protein alignments at tree-of-life scale using DIAMOND. Nat Methods 18, 366–368. 10.1038/s41592-021-01101-x.

73. Nakamura, T., Yamada, K.D., Tomii, K., and Katoh, K. (2018). Parallelization of MAFFT for large-scale multiple sequence alignments. Bioinformatics 34, 2490–2492. 10.1093/bioinformatics/bty121.

74. Naser-Khdour, S., Minh, B.Q., Zhang, W., Stone, E.A., and Lanfear, R. (2019). The Prevalence and Impact of Model Violations in Phylogenetic Analysis. Genome Biol. Evol. 11, 3341–3352. 10.1093/gbe/evz193.

75. Martinez-Perez, E., and Villeneuve, A.M. (2005). HTP-1-dependent constraints coordinate homolog pairing and synapsis and promote chiasma formation during C. elegans meiosis. Genes Dev 19, 2727–2743. 10.1101/gad.1338505.

76. Adilardi, R.S., and Dernburg, A.F. (2022). Robust, versatile DNA FISH probes for chromosome-specific repeats in Caenorhabditis elegans and Pristionchus pacificus. G3 (Bethesda) 12. 10.1093/g3journal/jkac121.

77. Preibisch, S., Saalfeld, S., and Tomancak, P. (2009). Globally optimal stitching of tiled 3D microscopic image acquisitions. Bioinformatics 25, 1463–1465. 10.1093/bioinformatics/btp184.

